# Enamel proteins reveal biological sex and genetic variability within southern African *Paranthropus*

**DOI:** 10.1101/2023.07.03.547326

**Authors:** Palesa P. Madupe, Claire Koenig, Ioannis Patramanis, Patrick L. Rüther, Nomawethu Hlazo, Meaghan Mackie, Mirriam Tawane, Johanna Krueger, Alberto J. Taurozzi, Gaudry Troché, Job Kibii, Robyn Pickering, Marc Dickinson, Yonatan Sahle, Dipuo Kgotleng, Charles Musiba, Fredrick Manthi, Liam Bell, Michelle DuPlessis, Catherine Gilbert, Bernhard Zipfel, Lukas F. K. Kuderna, Esther Lizano, Frido Welker, Pelagia Kyriakidou, Jürgen Cox, Catherine Mollereau, Caroline Tokarski, Jonathan Blackburn, Jazmín Ramos-Madrigal, Tomas Marques-Bonet, Kirsty Penkman, Clément Zanolli, Lauren Schroeder, Fernando Racimo, Jesper V. Olsen, Rebecca R. Ackermann, Enrico Cappellini

## Abstract

The evolutionary relationships among extinct African hominin taxa are highly debated and largely unresolved, due in part to a lack of molecular data. Even within taxa, it is not always clear, based on morphology alone, whether ranges of variation are due to sexual dimorphism versus potentially undescribed taxonomic diversity. For *Paranthropus robustus*, a Pleistocene hominin found only in South Africa, both phylogenetic relationships to other taxa ^1,2^ and the nature of intraspecific variation ^3–6^ are still disputed. Here we report the mass spectrometric (MS) sequencing of enamel proteomes from four ca. 2 million year (Ma) old dental specimens attributed morphologically to *P. robustus,* from the site of Swartkrans. The identification of AMELY-specific peptides and semi-quantitative MS data analysis enabled us to determine the biological sex of all the specimens. Our combined molecular and morphometric data also provide compelling evidence of a significant degree of variation within southern African *Paranthropus*, as previously suggested based on morphology alone ^6^. Finally, the molecular data also confirm the taxonomic placement of *Paranthropus* within the hominin clade. This study demonstrates the feasibility of recovering informative Early Pleistocene hominin enamel proteins from Africa. Crucially, it also shows how the analysis of these proteins can contribute to understanding whether hominin morphological variation is due to sexual dimorphism or to taxonomic differences. We anticipate that this approach can be widely applied to geologically-comparable sites within South Africa, and possibly more broadly across the continent.

---

While our understanding of the evolutionary relationships among Middle to Late Pleistocene hominins is becoming increasingly clear, in large part due to ancient DNA (aDNA) sequencing data, the relationships among earlier Plio-Pleistocene hominins remain unresolved. The genus *Paranthropus* evolved ca. 2.8 million years ago (Ma) and persisted until 1 Ma, coexisting in time and space with a number of other hominins, including *Australopithecus* species and members of the genus *Homo*. Resolving the relationships among these taxa is key to understanding the origins of our lineage. Even within *Paranthropus*, the phylogenetic relationships among the three currently identified species have been the subject of considerable discussion. Most researchers consider *Paranthropus* taxa to be monophyletic ^7^, however, morphological similarities between *P. robustus* and *Au. africanus* in a South African context ^1,2,8^, and between *P. aethiopicus* and *Au. afarensis* in an eastern African context ^9–11^ (Fig. 1A), have raised the possibility of paraphyly or even admixture between species ^12,13^. Furthermore, analyses of the enamel-dental junction (EDJ) of southern African *Paranthropus* indicate significant variation, suggesting the possibility of substructuring within *P. robustus* ^5^ or even the presence of more than a single species of this genus in paleoanthropological record of this region ^6,14^. Determining to what extent the variation within and between Plio-Pleistocene hominins is due to evolutionary diversification versus sexual dimorphism is therefore fundamental to resolving these relationships.

**Figure 1:**
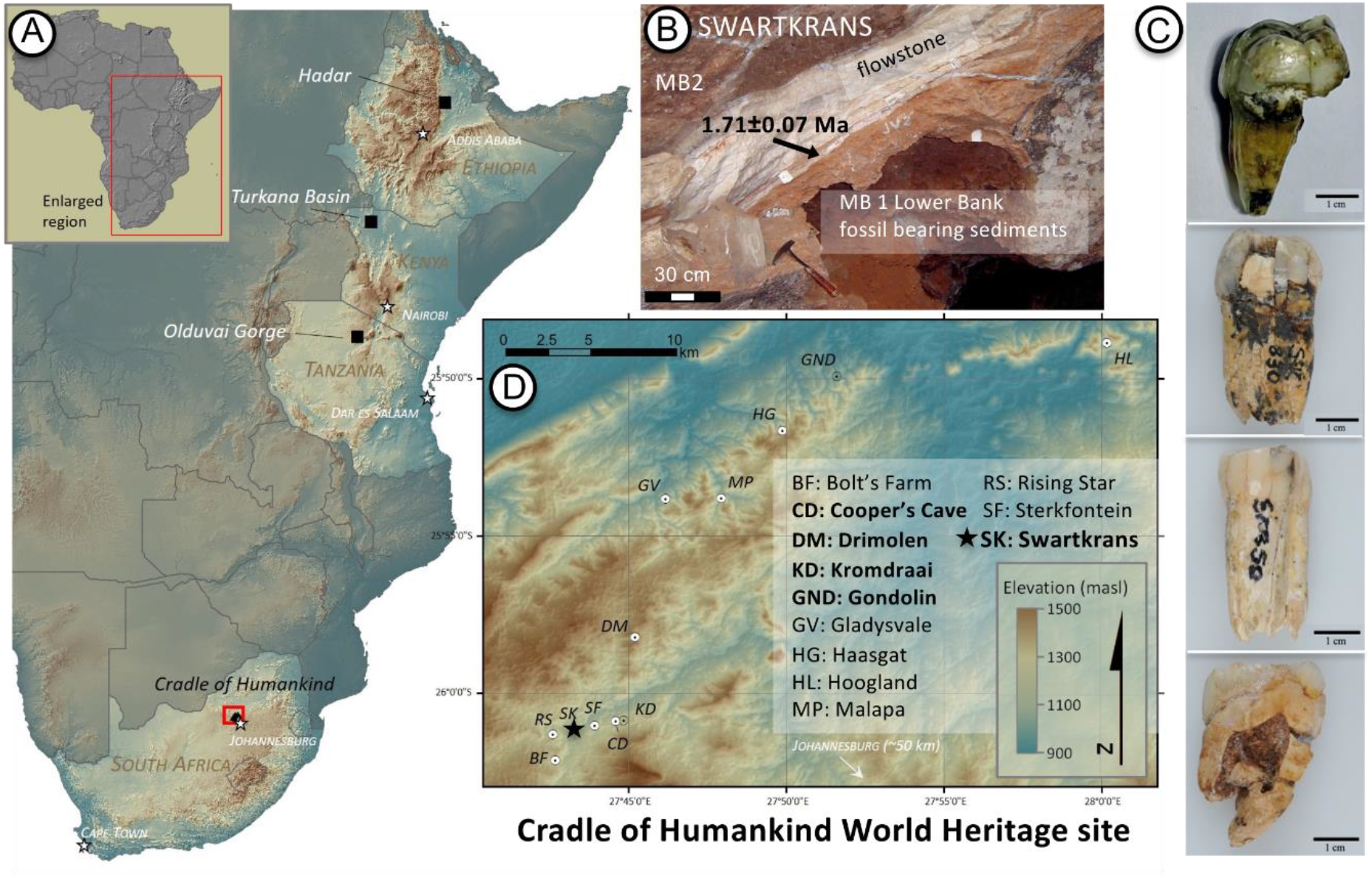
Location and cave structure of the site of Swartkrans, South Africa. A - Topographical map of the African continent (inset) showing the major early hominin fossil bearing regions. B - Photograph of the *Paranthropus* bearing palaeocave in Swartkrans, showing the Member 1 and dated flowstone ^74^. C - *Paranthropus* teeth analysed, from top to bottom: SK 835, a left M^3^; SK 830, a left P_4_; SK 850, a right P_3_; SK 14132, a right M^3^ (Supplementary Information 2.3). D - Enlarged view of the Cradle of Humankind World Heritage Site in South Africa (shown in A), with *Paranthropu*s fossil locality names in bold. Swartkrans is marked with a star.

Genetic data have provided insights of unprecedented resolution into human demography and evolution. Although aDNA has been crucial to clarify human history and admixture ^15–22^, it has never been successfully recovered from African hominin material older than ~ 0.018 Ma ^20^. As phylogenetically informative ancient proteins have been retrieved beyond the limits of aDNA preservation in Eurasia ^23–25^, we attempted their recovery to investigate Plio-Pleistocene hominin variation in Africa. Accordingly, we used liquid chromatography (LC) coupled to high-resolution orbitrap tandem mass spectrometry (LC-MS/MS) to sequence dental enamel protein remains from four southern African hominin specimens usually attributed to *P. robustus* (Fig. 1C and Supplementary Information 2.3). The LC-MS/MS data allowed the determination of the biological sex for all four specimens, as well as the exploratory characterisation of intraspecific genetic diversity and evolutionary relationships with other hominins. The downstream integration of proteomics and morphological analysis further highlighted taxon diversity within the sample set.

The four hominin fossils analysed originated from Swartkrans cave (Fig. 1B), located approximately 40 km northwest of Johannesburg, in South Africa’s Cradle of Humankind World Heritage Site (Fig. 1D). The teeth are from the oldest deposits at Swartkrans, Member 1 (MB1), which is dated to between 1.8 and 2.2 Ma ^26,27^. Swartkrans has produced the largest collection of specimens attributed to *P. robustus* in the world, but the taxonomic placement of this material has been the subject of various debates ^7,28–33^.

To maximise the breadth and depth of amino acid sequence coverage, manual off-line high-pH reversed-phase fractionation was carried out on stage-tips ^34^. This strategy extended the dynamic range of the less complex fractions for subsequent MS analysis ^35–37^, increasing peptide identifications in all four *Paranthropus* samples (Extended Data Fig. 2). The number of recovered amino acid positions increased up to 17%, and the number of PSMs (Peptide-Spectrum Matches) increased up to 3-fold (Supplementary Information 9.3). Further methodological development was achieved with the creation of an automated and open-source sequence assembly pipeline (Extended Data Fig. 4). A site-based sequence reconstruction approach^38^ was developed to generate consensus sequences directly from MaxQuant’s output tables. This sequence assembly pipeline enabled faster, more reproducible, and transparent data analysis processes. The generated outputs can be traced back to the original fragmentation spectra, thereby simplifying manual validation of ambiguous hits.

The combined analysis of the LC-MS/MS data obtained from fractionated and single-shot samples of each of the four individuals resulted in 4,600 to 8,500 PSMs covering 540 to 780 amino acid positions from 8 to 10 enamel-specific proteins (Extended Data Fig. 1A), 6 of which (ALB, AMELX, COL17A1, ENAM and MMP20) appear in all specimens (Ext. Data Fig. 1B). A total of 425 amino acid positions were consistently identified in all four *Paranthropus* specimens (Extended Data Fig. 1C), indicating that the majority of the covered positions were shared across all the samples. For validation, the MS workflow was successfully replicated in a proteomics laboratory in South Africa (Supplementary Information 9.9). The authenticity of the recovered sequences was supported by multiple lines of evidence. First, expected levels of racemisation and extent of peptide bond hydrolysis between free and bound amino acids indicate that, in all the samples, the dental enamel behaved as a closed system (Extended Data Fig. 3, Supplementary Information 8.1). Equally, no or negligible exogenous contamination across all four specimens is also supported by the high similarity of the amino acid composition profiles observed both within our sample set, and in comparison to other ancient dental enamel specimens previously investigated (Supplementary Information 8.1). Second, all the samples showed advanced rates of diagenetically-induced amino acid modifications, such as glutamine and asparagine deamidation and arginine to ornithine conversion, compatible with the age and the geographic origin of the *Paranthropus* specimens (Extended Data Fig. 1E-F). Additionally, we also observed extended oxidative degradation of histidine, phenylalanine, tyrosine and tryptophan (Supplementary Information 9.7). Third, the peptide length distribution was skewed towards shorter amino acid chains, as previously observed in other palaeoanthropological material and as supported by the chiral amino acid data (Supplementary Information 8.1), most likely due to extensive terminal hydrolysis (Extended Data Fig. 1D). Altogether, this evidence supports the authenticity of the ancient protein sequences we report. The attempt to detect protein-protein crosslinks did not lead to any confident identification (Supplementary Information 9.8).

Specimens SK 850 and SK 835 were unambiguously identified as male *Paranthropus* individuals based on the observation of multiple overlapping AMELY-specific peptides (Fig. 2A and Supplementary Information 9.6). No AMELY-specific peptide was detected in SK 830 and SK 14132. This absence, however, cannot alone lead to a female identification. Such an observation is also compatible with the possibility that these specimens originated from male individuals whose signal for the AMELY-specific peptides falls under the instrument’s detection limit. Here, assuming constant AMELX and AMELY ratios across the entire sample set investigated and using a site-based semi-quantitative approach, we defined the AMELX intensity threshold above which we would expect to detect AMELY-specific peptides, if present in the sample^39^. Since the AMELX site intensities of both SK 830 and SK 14132 were measured above the defined intensity threshold, we demonstrated that both specimens most likely originated from female individuals (Supplementary Information 9.6). We then integrated our molecular-based sex identifications with the conclusions of more established morphological methods, largely based on overall size. Specifically, we compared the available buccolingual (BL) and mesiodistal (MD) measurements of SK 830, SK 835, and SK 850 (SK 14132 not included due to missing data) to other relevant dental specimens of southern African *Paranthropus* (Supplementary Information 11.1). SK 830, assigned to a female individual based on molecular evidence, has MD and BL measurements consistent with previously identified female specimens (Extended Data Fig. 9B). SK 850, assigned to a male based on AMELY-peptides, has a MD measurement falling within the lower range of size variability seen among specimens previously assigned to males (Extended Data Fig. 10C). SK 835, also confidently identified through AMELY-peptides as originating from a male, shows a MD measurement falling toward the lower end of the entire size variability considered and a BL measurement falling at the centre of both the size distribution and the bivariate plot (Extended Data Fig. 9A). Interestingly, its protein-based male attribution is consistent with a previous study supporting philopatry in *P. robustus*, where SK 835 was shown to have a local (male) strontium isotope signal^40^. Our result however contradicts the recent assignment of SK 835 to a female individual based on its small size^41^.

**Figure 2:**
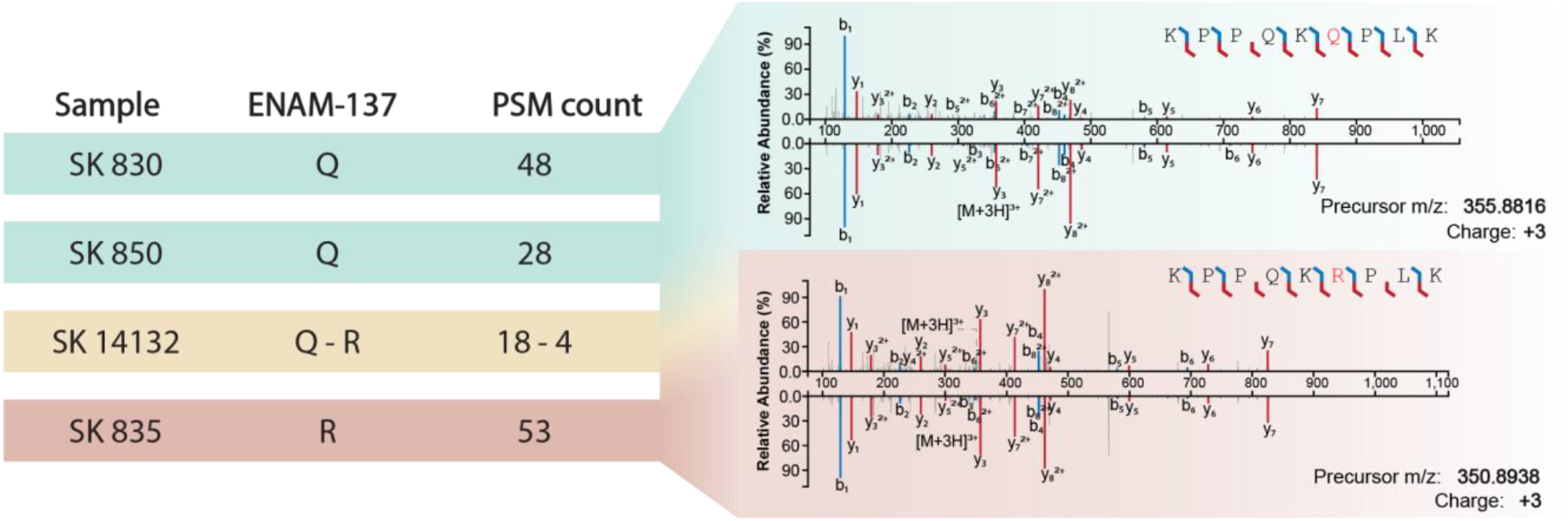
Sex identification of the four *Paranthropus* specimens. A - MS2 spectra of AMELY-specific peptides detected in the two male individuals: SK 850 and SK 830. In both spectra the detection of the methionine in position 59, characteristic to males, is well supported by the ion series. B - Site intensities of AMELY-59 as a function of AMELX-60 for all four *Paranthropus* samples. The site intensity represents the sum of the intensities of all PSMs covering that site. The horizontal cut-off represents the minimum intensity of an AMELY-specific peptide covering the AMELY-59 position in a male individual. The vertical cut-off represents the minimum intensity of a peptide covering the AMELX-60 position in a male individual.

To attempt molecular-based phylogenetic reconstruction, the assembled protein sequences from all four *Paranthropus* specimens were aligned to their orthologs from a large reference dataset including multiple individuals of extant and extinct Hominidae species. We also created a subset of that reference database, keeping a single individual for each species (Supplementary Information 10.1). The reconstructed sequences were analysed using a semi-automated sequence assembly workflow, allowing for full reproducibility and transparency across the entire analysis (Extended Data Fig. 5) ^42^. Based on the phylogenetically informative sites identified (Table S13, S18-19), all four *Paranthropus* sequences were closer to those in the *Homo* clade than to any other primate (Extended Data Fig. 6). In the four *Paranthropus* individuals, we detected a total of 17 Hominidae species-informative single amino acid polymorphisms (SAPs) (Table S13), only two of which showing an allelic state different from the present-day humans, Neanderthals, and Denisovans’.

The first SAP site, COL17A1-636, i.e. position 635 in the *Homo sapiens* canonical Ensembl transcript of COL17A1 - ENST00000648076.2, was observed in SK 835 and SK 14132. Both specimens carried the ancestral allele shared with non-hominins, while present-day humans, Neanderthals, and Denisovans display the derived allele (Extended Data Fig. 7). The second SAP site, ENAM–137, i.e. position 137 in the *Homo sapiens* canonical Ensembl transcript of ENAM - ENST00000396073.4, was recovered in all four specimens and appears to be variable between them. More specifically, SK 830 and SK 850 bear what we parsimoniously interpret as a fully deamidated glutamine (Q), while SK 835 bears an arginine (R) in that position (Fig. 3). Interestingly, in SK 14132 the ENAM-137 site appeared to be heterozygous, with the Q allele covered in 80% of the PSMs (18 vs 4). The confident identification of the two ENAM-137 alleles was further confirmed by synthetic peptide analysis (Fig. 3). Arginine at ENAM-137 is shared with all African apes, while glutamine is reported only in the *Pongo* clade, where it is fixed. Looking at a primate-wide dataset, the R to Q substitution appears in ENAM-137 in different branches of the tree (Extended Data Fig. 7). Although the probability of this substitution varies across different models ^43–45^, in modern humans a single non-synonymous G to A transition from the codon CGG to CAG is sufficient for it to occur on ENAM-137. Accordingly, our results indicate that this substitution most likely occurred independently in *Paranthropus*, but did not fixate within all the wider populations or groups of this genus.

**Figure 3:**
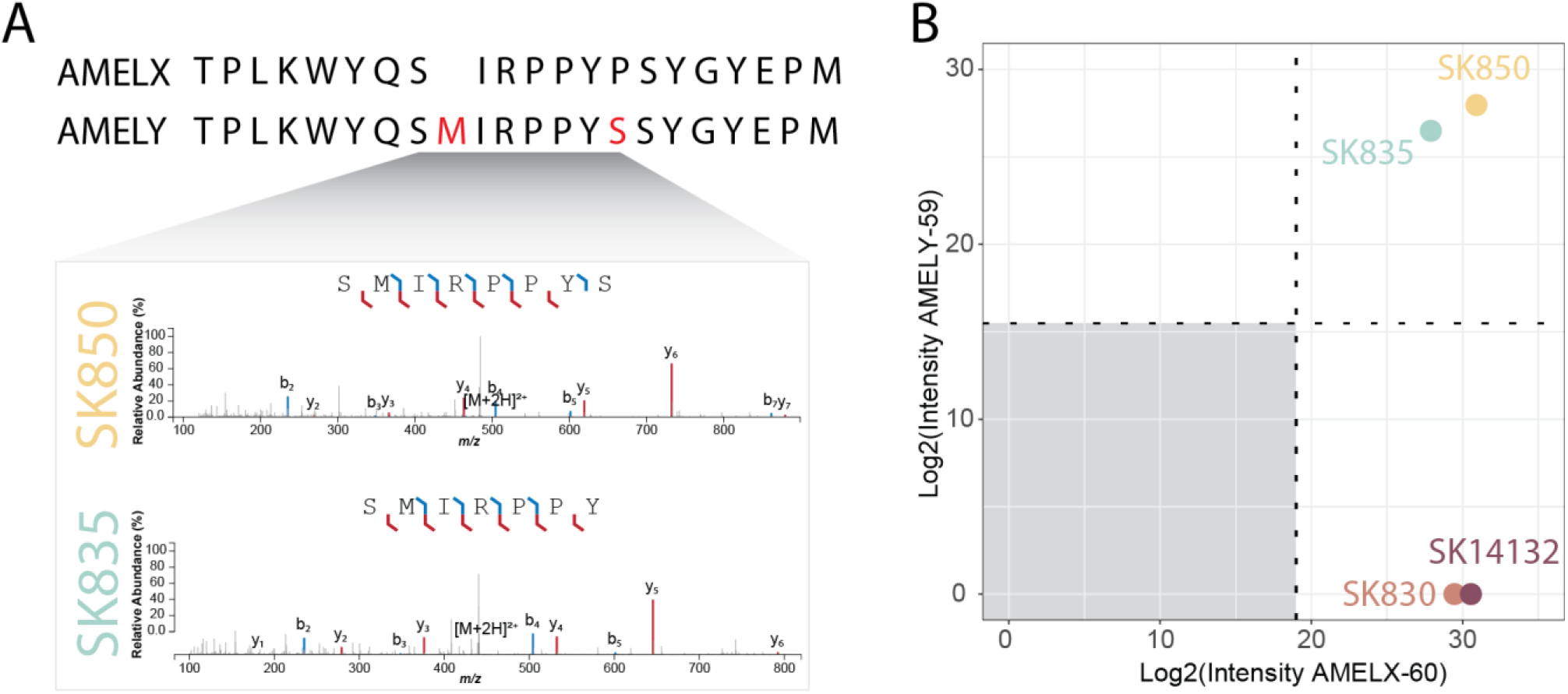
Sequence variation within the *Paranthropus* individuals. Amino acid sequence variation at ENAM-137 in the four *Paranthropus* samples and number of PSMs supporting their detection. The detection of glutamine (Q) and arginine (R) at ENAM-137 was validated with synthetic peptides. On the right, the mirror plots represent the MS2 spectra covering glutamine, in peptide KPPQKQPLK, and arginine, in peptide KPPQKRPLK, in the *Paranthropus* samples (top) compared to the MS2 spectra of the corresponding synthetic peptides (bottom).

Within-species variation similar to that observed on ENAM-137 is expected for any large enough dataset of amino acid sequences, but might be unexpected for a sample size of 4 individuals. To assess this, we repeatedly sampled, as a proxy, 4 randomly selected individuals from a global sample of present-day humans, leveraging the abundance of available genetic data, and estimated the probability of observing variation on the protein level in the region of interest (Methods). We found it plausible that genetic variants segregating within a given species could manifest as amino acid differences in a sample of the same size as the one we have for *Paranthropus*, though we note that the effective population size of humans today likely differs from that of *Paranthropus* (Supplementary Information 10.4), leaving any taxonomic conclusions on the basis of this genetic variation alone still premature.

To explore the taxonomic placement of the *Paranthropus* individuals, we utilised our aligned reference datasets to generate phylogenetic trees via a maximum likelihood ^46^ and a Bayesian ^47^ approach (Fig. 4A, Fig. S18-S21). The phylogenetic reconstructions place the *Paranthropus* individuals as outgroups to the clade containing present-day humans and available Pleistocene hominins from Eurasia (Neanderthal and Denisovan). All these individuals, including *Paranthropus*, form a clade to the exclusion of other members of present-day Hominidae. Three of the *Paranthropus* specimens, i.e. SK 850, SK 14132 and SK 830, form a clade, while SK 835 appears to be more distantly related to them. These results were also independently validated by repeating the phylogenetic analysis using an alternative workflow and reference dataset (Supplementary Information 10.3.3).

**Figure 4:**
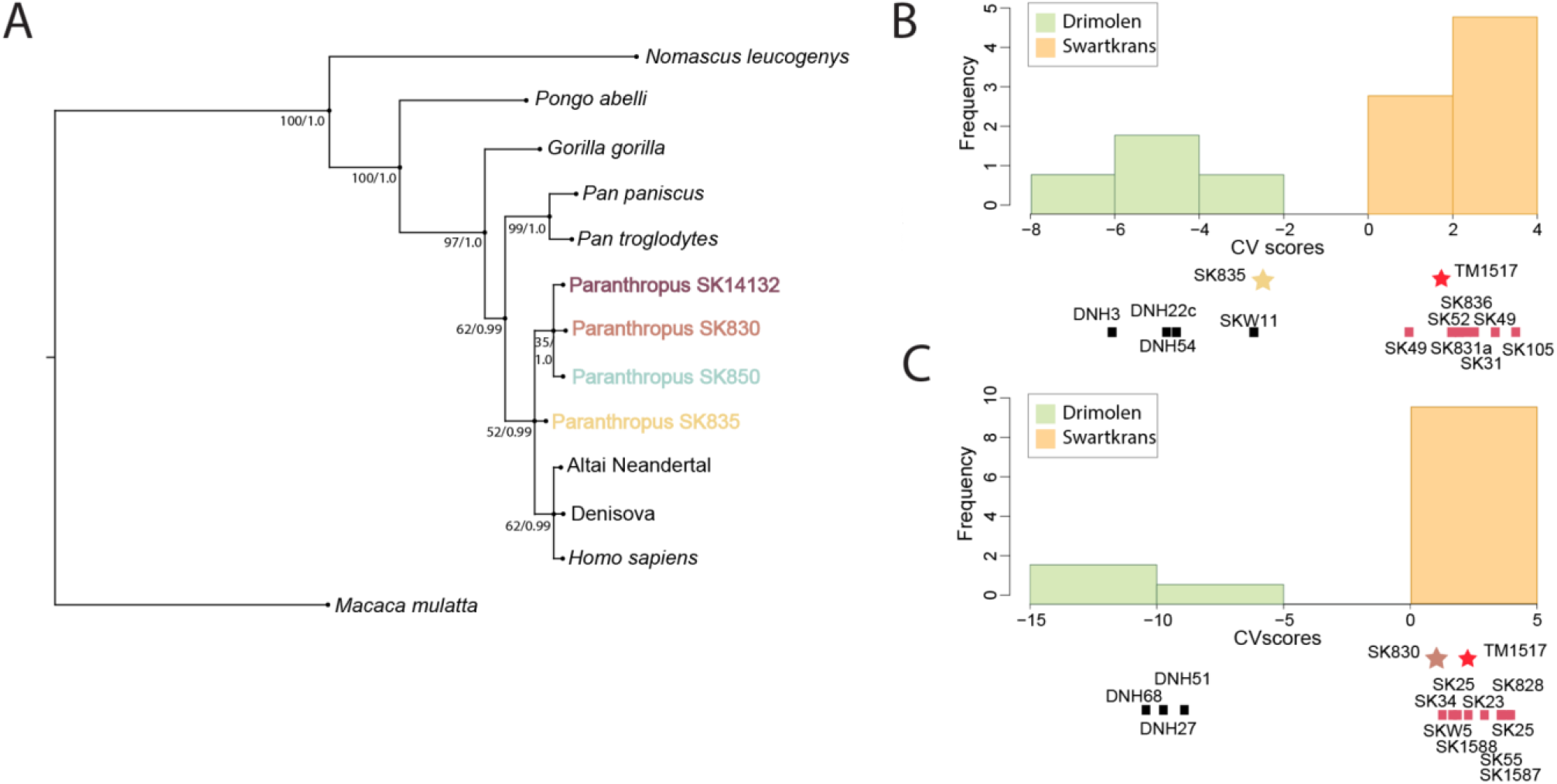
Phylogenetic tree and geometric morphometric analysis of the enamel-dentine junction. A - Bayesian and maximum likelihood phylogeny and divergence of the four *Paranthropus* samples among extant apes, with *Macaca mulatta* shown as outlier. The node values indicate the maximum likelihood values (0– 100) from PhyML and the Bayesian probability (0–1) from MrBayes. B - Frequency and distribution of canonical variate scores of the Swartkrans and Kromdraai, and Drimolen M^3^s enamel-dentine junction shape analysis. This analysis shows SK 835 as statistically closer to specimens from Drimolen compared to other specimens from Swartkrans and the holotype of *P. robustus* TM 1517 from Kromdraai. C - Same as in B but using P_4_s, showing that SK 830 is closer to Swartkrans and Kromdraai specimens compared to other specimens from Drimolen.

Of note, while the Bayesian tree places high posterior probabilities on the reconstructed divergences within *Paranthropus*, the bootstrap resampling of the maximum likelihood approach shows only moderate support for these divergences. Repeating the same phylogenetic workflow with a larger dataset containing several present-day Hominidae individuals also showed only moderate support for these relationships (Fig. S22-S24). When looking at each protein individually, none of the gene trees exactly match the topology of the concatenated phylogenetic trees obtained from all the sequenced proteins (Supplementary Document 4). These results are likely due to a combination of missing data as well as incomplete lineage sorting.

To integrate the evidence obtained with palaeoproteomics and more established approaches, geometric morphometric analysis of the enamel-dentine junction (EDJ) was carried out on the two morphologically best-preserved specimens (SK 835 and SK 830). The results unambiguously show that they both belonged to *Paranthropus* and differ from *Australopithecus* and early *Homo* (Extended Data Fig. 8; Supplementary Information 11.2). Noticeably, the EDJ shape of SK 835 M^3^, the individual bearing an arginine at ENAM-137, was statistically more similar to the *Paranthropus* specimens from the site of Drimolen than to those from Swartkrans and Kromdraai assemblages, which include the holotype of *P. robustus* (TM 1517). The EDJ shape of the P_4_ SK 830, in contrast, more closely resembles the latter group and statistically differed from the material from Drimolen (Fig. 4 A-B). The BL and MD measures further indicated high levels of variation within samples currently referred to as *P. robustus* (Supplementary Information 11). In the bivariate plot, specimens from Drimolen and Swartkrans largely grouped separately (Extended Data Fig. 9), with some overlap among larger specimens. SK 835 fell within the convex hull of Swartkrans, but close to the Drimolen grouping, while SK 830 grouped with other Swartkrans specimens (Extended Data Fig. 9).

## Discussion and broader implications

Here we report the recovery of Early Pleistocene hominin proteins from Africa. The *Paranthropus* specimens we studied were recovered from cave sediments mostly composed of remobilized soil from outside the cave ^48^. Altogether, their rapid accumulation through episodic flash floods and the relative aridity ^49^, coupled with extensive cementation, favourably contributed to the preservation of the proteins within the fossil teeth. Whether or not protein preservation would be comparable in other early hominin-bearing deposits, including open-air sites such as those found in other places in Africa, is an open question, and future work should focus on testing the feasibility of palaeoproteomic analysis while minimising damage to precious African heritage.

The application of manual off-line reversed-phase high-pH fractionation improved the dental enamel protein sequence coverage (Extended Data Fig. 2), revealing the existence of diversity at the protein level within southern African *Paranthropus*. In addition, using spectral prediction software and sequencing synthetic peptides helped validate spectra that provided mass spectrometric evidence to confirm this diversity and detect heterozygosity. Mass spectrometry has been previously utilised as a method of choice for the detection of heterozygosity in modern human individuals ^50–52^. To our knowledge, however, this has never been previously applied in the context of palaeoproteomics. Future studies should explore this aspect, as heterozygosity is an additional source of genetic information.

Our results provide an initial insight into the genetic relationships between *Paranthropus* and other hominins. We find that the sequences we recover place *Paranthropus* within hominins and as an outgroup to the clade including *Homo sapiens*, Neanderthals and Denisovans. The trees presented here, however, only provide a tentative phylogenetic placement of *Paranthropus*, as they are based on a small set of proteins, whose genes may be subject to incomplete lineage sorting ^25,53–60^. Similarly, the individual gene trees can only be inferred partially, as we can only observe mutations that have resulted in amino acid substitutions and that are located on peptides covered by mass spectrometry, while the strict manual filtering scheme required for preventing false positives further limits the number of phylogenetically-informative sites available to us (see Methods). Despite these caveats, the recovery of ca. 2Ma phylogenetically-informative genetic material in African hominins can be considered a potentially transformative breakthrough for palaeoanthropology. Future methodological developments should focus on the recovery of richer proteomes, beyond dental enamel, and should be applied to the investigation of other Early Pleistocene hominin species, such as *P. boisei*, *Au. africanus*, *Au. sediba*, and *H. erectus*, to further refine the molecular-based reconstruction of hominin phylogeny.

Due to a single SAP (ENAM–137), our reconstructed trees suggest that one of the *Paranthropus* individuals (SK835) might be more distantly related to the other three individuals than they are to each other. Though potentially the result of incomplete lineage sorting, it is also possible that this individual may have belonged to a distinct *Paranthropus* group with a relatively recent separation from the group to which the other three individuals belong. The hypothesis of two *Paranthropus* groups is further supported when the palaeoproteomics results are integrated with the morphological variability observed using EDJ analysis. The *Paranthropus* assemblage within South Africa exhibits considerable size variation, most of which has previously been attributed to sexual dimorphism and possibly reflecting a gorilla-like pattern of extended growth for males (bimaturation) ^61^. However, recent studies have suggested that these morphological differences might indicate either different species ^6^, or site-related diversity within a single species over time, i.e. micro-evolutionary changes, following a morphocline ^3,4,62^. The four analysed specimens from Swartkrans are from Member 1, and here considered as penecontemporaneous, eliminating the possibility that the differences between them represent change over time within a species, and supporting instead greater taxon diversity, possibly at the species level. We couch this interpretation within the caveat that the time window in which Member 1 sediments accumulated is long (500,000 years at most), and due to the way in which the fossils were collected, we do not have exact provenience data for them. However, while Member 1 covers a long time interval, the fossil-bearing sedimentary deposits likely accumulated rapidly, supporting our penecontemporaneous interpretation of the *Paranthropus* teeth analysed here. If the hypothesis of two different *Paranthropus* groups holds, we would assume that individuals from the site of Drimolen, which are morphologically closer to SK 835, would also share the same SAP for ENAM-137. Given that the site of ENAM-137 was covered with high confidence in all four of our samples, we anticipate that future sampling of *Paranthropus* from Drimolen would cover this SAP as well, and anticipate that further research could test this hypothesis.

The confident identification of both male and female *Paranthropus* individuals demonstrates the limitation of sexing techniques based on morphological (tooth) size, highlighting the greater resolution offered by proteomics. This capability has clear implications for our understanding and interpretation of morphological variation in the deep-time human fossil record, as it enables us to control for sexual dimorphism and by implication the range of anatomical variation, in our identification of hominin species. However, the method we use is reliant on the identification of the protein product deriving from the expression of the Amelogenin-Y gene. Deletions of this gene have been recorded both in modern human populations ^63,64^ as well as in one Neanderthal individual ^65^. Male individuals bearing such deletions cannot express AMELY and would therefore be identified as females using our approach.

Previous palaeoproteomic studies of extinct organisms have largely focused on single individuals to explore the genetic history of a species ^23–25,66,67^. Here we showcase that the analysis of multiple individuals, in conjunction with morphological data, can better represent that history and illuminate variation that might be indicative of inter- or even intra-taxon diversity. The successful recovery of million years old proteins should be achievable in other South African cave sites of similar age and geology, making biological sex identification and intra-species analysis possible. This study also brings palaeoproteomics closer to extracting similar data from other African early hominin material, specifically taxa such as *Au. afarensis* that are represented by a plethora of isolated and fragmentary dental remains. The Cradle of Humankind has yielded an exceptionally large number of hominin fossils, yet the greatest diversity of hominin taxa is currently known from eastern African sites, mainly in the rift valley regions of Ethiopia, Kenya and Tanzania. Whether and how much of this phyletic diversity is real, and not the result of methodological limitations and research(er) bias, remains a hotly debated topic. The coherent results obtained from this study combining molecular and morphological data has profound implications for addressing such long-standing controversies surrounding the nature and extent of Plio-Pleistocene hominin diversity ^68–73^.

## Methods

### Sample selection and site

Prior to analysis of hominin material, four bovid teeth were sampled for palaeoproteomic analysis, three of them originating from Swartkrans and one from Cooper’s Cave, South Africa (Supplementary Information 3). Both are *Paranthropus*-bearing localities in South Africa’s Cradle of Humankind World Heritage Site. Swartkrans is located approximately 40 km northwest of Johannesburg, with its oldest fossil bearing deposits dating to 1.8 to 2.2 Ma. Cooper’s Cave is about 4 km away from Swartkrans and preserves fossils dated to around 1.3 Ma. Following successful identification of proteins from the bovids, four teeth previously assigned to *Paranthropus robustus* were selected for analysis. All four sampled *Paranthropus* teeth came from Member 1 of Swartkrans (Supplementary Information 1 & 2), and are housed at Ditsong Museum, South Africa.

### Biomolecular preservation

Chiral amino acid analysis was undertaken on the four *Paranthropus* teeth. Enamel samples were powdered, and oxidative treatment was used to isolate the intra-crystalline fraction of amino acids ^75^. Two subsamples per specimen were subjected to analysis. One subsample was demineralized for the free amino acid analysis, and the second subsample was treated to release the peptide-bound (total hydrolysable) amino acids. Samples were analysed in duplicate by reverse-phase high-performance liquid chromatography, with standards and blanks run alongside (Supplementary Information 4). The values of the D/L stereoisomers of aspartic acid plus asparagine, glutamic acid plus glutamine, phenylalanine and alanine were assessed to provide an overall estimate of intra-crystalline protein decomposition (Extended Data Fig. 3).

### Peptide extraction

All reagents were supplied by Sigma-Aldrich unless stated otherwise. Peptide extraction took place in the clean laboratory for ancient biomolecule extraction at the University of Copenhagen. Filtered ventilation and positive pressure were used in combination with personal protective equipment to minimise modern protein contamination. An extraction blank was prepared in parallel with, and processed in parallel to, the ancient samples. Enamel was cleaned and separated from dentine using a rotating cutting disc connected to an electrically operated rotary tool. Peptide extraction was performed following the procedure described in Cappellini et al.^23^. Briefly, powdered enamel was demineralized twice overnight at 4 °C in 10% trifluoroacetic acid (TFA) and used for single shot and fractionation analysis. Over all experiments, a total of 80-250 mg per tooth was used.

### Peptide clean-up and fractionation

Peptides were loaded on in-house built StageTips after conditioning with methanol and equilibration using 0.1% TFA. Salts were then removed with 0.1% TFA ^23^. The stage-tips were either directly eluted for single-shot LC-MS/MS analysis or subjected to high-pH reversed-phase offline fractionation. 60-200 mg of enamel were used for fractionation using an increasing gradient of acetonitrile (ACN). Buffer A was composed of 5 mM ammonium bicarbonate (pH 7.8) and the peptides were eluted with 0 % (pH adjustment fraction), 11 %, 15 %, 20 %, 70 % of buffer ACN. The fractions were acidified using 1 % TFA and the organic solvent was removed by vacuum evaporation. For separation by EASY-nLC, the peptides were reconstituted in 0.1 % TFA 5 % ACN in water, whereas Evosep chromatography required the peptides to be loaded on disposable C18 trap columns (Evotip, Evosep) following the manufacturer instructions.

### LC-MSMS

Peptides were separated on a liquid chromatographic system (EASY-nLC™ 1200 or Evosep One) coupled to an Exploris 480 mass spectrometer (Thermo Fisher Scientific, Bremen, Germany). Peptides were separated using an in-house packed column (15 cm x 75 μm, 1.9 μm) packed with C_18_ beads (Reprosil-AQ Pur, Dr. Maisch). The nanoLC was operated using a 77 min gradient. Buffer A was milliQ water, buffer B was 80% ACN, 0.1 % FA, and the flow rate was set at 250 nL/min. The Evosep One separation was performed using the predefined commercial gradient 20 SPD, operating at a flow rate of 200 nL/min. The mass spectrometer was operated in Data Dependent Acquisition (DDA) in positive mode. Full MS resolution was set at 120,000 at m/z 200 with a maximum injection time of 25 ms. The HCD fragment spectra resolution was set at 60,000 with a maximum injection time of 118 ms and a Top10 method with 30 seconds dynamic exclusion. The isolation window was set at 1.2 Th and the normalised collision energy at 30%. Injection blanks were run prior to every sample injection to limit the risk of carry over in the chromatographic column. Faunal samples were analysed on a Q Exactive HF-X (Thermo Fisher Scientific, Bremen, Germany) using the same parameters as described above, except for the normalised collision energy that was set at 28%.

### Database search strategy

Raw data was searched using MaxQuant (V. 1.6.0.17)^76^, Peaks Client 7.0 ^77^ and pFind (V. 3.1.5)^78^. A custom-built database, containing 622 entries of enamel specific proteins collected from Uniprot corresponding to human, chimpanzee, gorilla and orangutan, was used. This database was supplemented with Gigantopithecus, Homo antecessor, Denisovan enamel proteome and some Neanderthal enamel specific sequences ^24,25^ (Supplementary Information 9.1).

### PTM analysis

The extent of deamidation was estimated using the python script from Mackie et al. ^79^. Other modifications were assessed by PSM counting. For a given modification, the ratio between the number of amino acids that could be modified to the number of amino acids actually modified is calculated. The counts were normalised by the MS count.

### Protein sequence reconstruction

Protein sequences were reconstructed using an in-house built R script. In short, an aligned database was used for indexing the detected peptide sequences to the correct gene and position. Experimental annotations from the summary.txt file were added to the output tables. The delta score was recalculated and a global FDR control at 1 % at the PSM level was ensured. The filtered indexed sequences were then converted into site-level data by expanding the table from peptides to amino acids. At this stage, the consistency of every amino acid position detected can be evaluated with metrics such as the site intensity or the PSM coverage for each sample. Conflicting positions, where evidence for two different amino acids for one given position in one sample is available, were elucidated based on the most likely outcome for the generation of the consensus sequence. The covered positions were then aligned at the gene level, and exported as FASTA files. An exclusion list, containing sequences manually filtered can be included in the script (Extended Data Fig. 4). Peptide sequences were discarded if: (i) the ion coverage was not sufficient to validate the presence or absence of a SAP, (ii) the covered region was unspecific to the enamel proteome and could be from bacterial origin, (iii) the peptide was also be identified in the lab blank, or (iv) the fragment ion intensity would not match the machine learning generated predictions.

### Reference datasets

The phylogenetic analysis was performed using two assembled reference datasets. The first dataset included a single representative protein sequence from each of the following species: *Homo sapiens*, *Pan paniscus*, *Pan troglodytes*, *Gorilla gorilla*, *Pongo abelii*, *Macaca mulatta*, *Macaca nemestrina*, *Microcebus murinus*, *Nomascus leucogenys*. Protein sequences for this dataset were obtained from Ensembl, Uniprot ^80,81^ as well as in-silico translations (described below), when neither online database contained the required protein. The second dataset included the in-silico translated proteins from 217 individuals covering all extant and some extinct members of Hominidae along with some additional primates. This dataset aims to represent the true within-species diversity of the Hominidae clade including that of extinct populations such as Neanderthals, Denisovans and ancient modern humans. Protein sequences for this dataset were acquired from the ‘Hominid Palaeoproteomic Reference Dataset’ available at Zenodo (https://zenodo.org/record/7728060#.ZBMGT3bMKbh). A third reference dataset was created by the co-authors in Barcelona to corroborate the results. This dataset is labelled ‘independent’ and its creation is described in Supplementary Information 6.1.3.

### Alignment, I/L correction and Phylogenetic Tree generation

The two reference datasets were merged with the reconstructed proteins of the four *Paranthropus* individuals. PalaeoProPhyler ^42^ was utilised and each merged dataset was processed through the same workflow (Extended Data Fig. 5): Sequences were divided into individual protein datasets, which were aligned using Mafft and then corrected for leucine and isoleucine isobaric positions in the same manner as described in Welker et al.^25^. The individual protein datasets were concatenated and converted to each of the appropriate formats for the phylogenetic analysis. A single phylogenetic tree was generated for each individual protein alignment as well as multiple phylogenetic trees for the concatenation of all the available proteins, using different software. PhyML ^46^ was used for the generation of the individual protein trees and both PhyML and Mrbayes ^47^ for the generation of the concatenated trees. Additional phylogenetic trees were generated using the concatenated datasets using IQTree ^82^ and Beast2 ^83^ (Fig. S21). The full command for each software can be found in the Supplementary Information 6. All trees were plotted using FigTree v.1.4.4 (http://tree.bio.ed.ac.uk/software/figtree/). The “independent” dataset was also utilised to generate multiple phylogenetic trees, as described in Supplementary Information 10.3.3.

### Pairwise distance matrix and heatmap

PHYLIP’s protdist ^84^ was applied onto the concatenated representative dataset using the default parameters. The resulting distance matrix (Fig. S20) was utilised to generate the distance heatmap (Extended Data Fig. 6) using R ^85^ and the ‘gplots’ (https://cran.r-project.org/web/packages/gplots/) and ‘RColorBrewer’ (https://cran.r-project.org/web/packages/RColorBrewer/) packages.

### Within Species Amino Acid Diversity

A custom workflow was developed to assess the probability of sampling four individuals from a species’ population and identifying variation on the protein level. The workflow was built using snakemake and utilising bcftools ^86^, biopython ^87^ and Ensembl’s VEP ^80^. Gnomad’s ^88^ ‘HGDP+1KG’ modern human callset (https://gnomad.broadinstitute.org/downloads#v3-hgdp-1kg) was used as the input of the workflow. The full details of this workflow are described in Supplementary Information 6.5. The code as well as short tutorial on how to reproduce the results are both available in a Github repository: (https://github.com/johnpatramanis/Code_for_Genetic_Diversity_Sampling)

### Homologous amino acids among primates

The reconstructed protein sequences from *Paranthropus* were analysed with another independently created reference dataset that included protein sequences from multiple primates as well as the previously published hominid enamel proteomes ^24,25^. The full details of the creation of the dataset can be found in Supplementary Information 6.1.3. The dataset was aligned using MAFFT v7.490 ^89^ and trimmed using TrimAl v1.2rev59 ^90^. Homologous amino acids of other primates were extracted from the MSA using SeqIO from biopython ^87^. The distribution of amino acids per species was then plotted in R v4.1.2 using the packages reshape2, ggplot2, ape, and ggimage.

### Morphological measurements

To assess size variation, buccolingual (BL) and mesiodistal (MD) measures of three of the four teeth analysed in this study, SK 830, SK 835 and SK 850, were compared to other relevant southern African dental remains currently attributed to *P. robustus*. The fourth specimen, SK 14132, was not included as these measurements were not attainable on this tooth. Measurements were collated from the published literature, except for SK 850, and are provided in Supplementary Document 3 with results further described in Supplementary Information 11.1. For specimens with both left and right dentition, measurements were averaged, but for all other specimens, side information was disregarded. Given the fragmentary nature of SK 850, only the mesiodistal dimension of this tooth was measured, which was extracted from a 3D scan of this specimen. Protocols for most MD and BL diameters follow that of Wood 1991^91^.

### Morphometric analysis

The enamel-dentine junction (EDJ) of the P_4_ SK 830 and of the M^3^ SK 835 ^41^ were analysed and compared with those of *Homo*, *Australopithecus* and *Paranthropus* (including TM 1517, the holotype of *P. robustus*; Table S4). Both SK 830 and SK 835 were scanned by X-ray microtomography with the General Electric V|Tome|x s industrial microCT system at the PLACAMAT platform (University of Bordeaux, France), and with the EasyTom XL Duo instrument at the PLATINA platform of the IC2MP (University of Poitiers, France) ^41^. The scans were done according to the following parameters: 70-110 kV voltage; 340-350 mA current; 0.1 mm Cu and 1.2 mm Al filters. The final volumes were reconstructed with an isotropic voxel size of 27.5 µm and 25 µm for SK 830 and SK 835, respectively. The comparative specimens were scanned using either X-ray or neutron microtomography and the final volumes were reconstructed with an isotropic voxel size of 10-30 µm for isolated teeth and 40-60 µm for jaw fragments ^6^. For all specimens, image stacks were imported into Avizo v.8.0 (FEI Visualization Sciences Group), and the images were segmented using semiautomatic procedures and an adaptation of the half-maximum height method ^92–94^. All the EDJ surfaces were generated using the ‘constrained smoothing’ option. The output data were imported in R with the package RToolsForDeformetrica v.0.1 ^95^. Using the packages ade4 v.1.7-6 ^96^ and Morpho v.2.8 ^97^ for R v.4.0.4 ^85^, we started by computing principal component analyses (PCA). We then performed cross-validated canonical variates analysis (CVA) based first on three groups assigned with equal prior probabilities (Early-Middle Pleistocene *Homo*, *Australopithecus* and *Paranthropus*) using the R package Morpho v.2.8 ^97^.

## Data Availability

The mass spectrometry proteomics data have been deposited to the ProteomeXchange Consortium (http://proteomecentral.proteomexchange.org) via the PRIDE partner repository ^98^ with the dataset identifier **PXD040221**.

Reference Datasets, XML files and phylogenetic results files are available on Zenodo: https://zenodo.org/record/7801259

## Code Availability

- Custom R-code for sequence assembly is available on GitHub at: https://github.com/ClaireKoenig/ProteinSequenceAssembly
- Genetic Variation analysis code is available on GitHub at: https://github.com/johnpatramanis/Code_for_Genetic_Diversity_Sampling
- Commands for the generation of the phylogenetic workflow are available in the Supplementary

## Acknowledgements

PPM, CK, IP, JK, CG, PK, JC, KP, FR, TMB, JVO and EC are supported by the European Union’s Horizon 2020 research and innovation programme under the Marie Sklodowska-Curie “PUSHH” training network, grant agreement No. 861389. PLR, EC, and JVO were supported by the European Commission through the MSC European Training Network ‘TEMPERA’ (grant number 722606). EC, LS, FR, JVO, RRA, AJT, MM and GT are supported by the European Research Council (ERC) under the European Union’s Horizon 2020 research and innovation programme (grant agreement No. 101021361). MM was supported in part by a grant from the Danish National Research Foundation award to Matthew Collins (PROTEIOS, DNRF128). Work at The Novo Nordisk Foundation Center for Protein Research (CPR) is funded in part by a donation from the Novo Nordisk Foundation (NNF14CC0001). RRA and NH are supported by the African Origins Platform, National Research Foundation of South Africa (grant no. 117670 and 136512). LS is funded by a Discovery Grant from the Natural Sciences and Engineering Research Council of Canada (NSERC Discovery Grant No. RGPIN-2020-04159). FW and GT are supported by the European Research Council (ERC) under the European Union’s Horizon 2020 research and innovation programme (grant agreement No. 948365). FR is supported by a Villum Young Investigator Grant (project no. 00025300), a Novo Nordisk Fonden Data Science Ascending Investigator Award (NNF22OC0076816) and by the European Research Council (ERC) under the European Union’s Horizon Europe programme (grant agreements No. 101077592 and 951385). TMB is supported by funding from the European Research Council (ERC) under the European Union’s Horizon 2020 research and innovation programme (grant agreement No. 864203). LB, MDP and JB would like to acknowledge the funding support received from DIPLOMICS, a research infrastructure initiative of the Department of Science and Innovation in South Africa for the validation work conducted at D-CYPHR. We acknowledge Ricardo Fong-Zazueta for creating and providing the protein translations of the ‘independent’ reference dataset. This research contributes towards the output of the Biogeochemistry Research Infrastructure Platform (BIOGRIP), supported by the Department of Science and Innovation, South Africa.

## Author Contributions

EC and RRA designed the study. PPM, CK, PLR, MM, AJT, GBT, NH, FW, LB, MP, MD, CZ, and LS performed experiments. PPM, CK, PLR, MM, IP, JK, CG, MD, LFKK, EL, FW, PK, JRM, CL, and LS analysed the data. JVO, RRA, EC, MT, BZ, TMB, KP and CZ provided material, reagents, or research infrastructure. PPM, CK, IP, RRA CZ, LS and EC wrote the manuscript with input from all the other authors.

## Additional information

### Supplementary Information

**Supplementary Document 1**-MS2 spectra supporting the detected SAP with spectral prediction.

**Supplementary Document 2**-Benchmark of spectral predictions for palaeoproteomics.

**Supplementary Document 3**-Paranthropus robustus standard dental measurements.

**Supplementary Document 4-**Maximum likelihood gene trees.

**Correspondence and requests for materials** should be addressed to JVO, RRA and/or EC.

### Extended data

**Extended Data Fig. 1.**
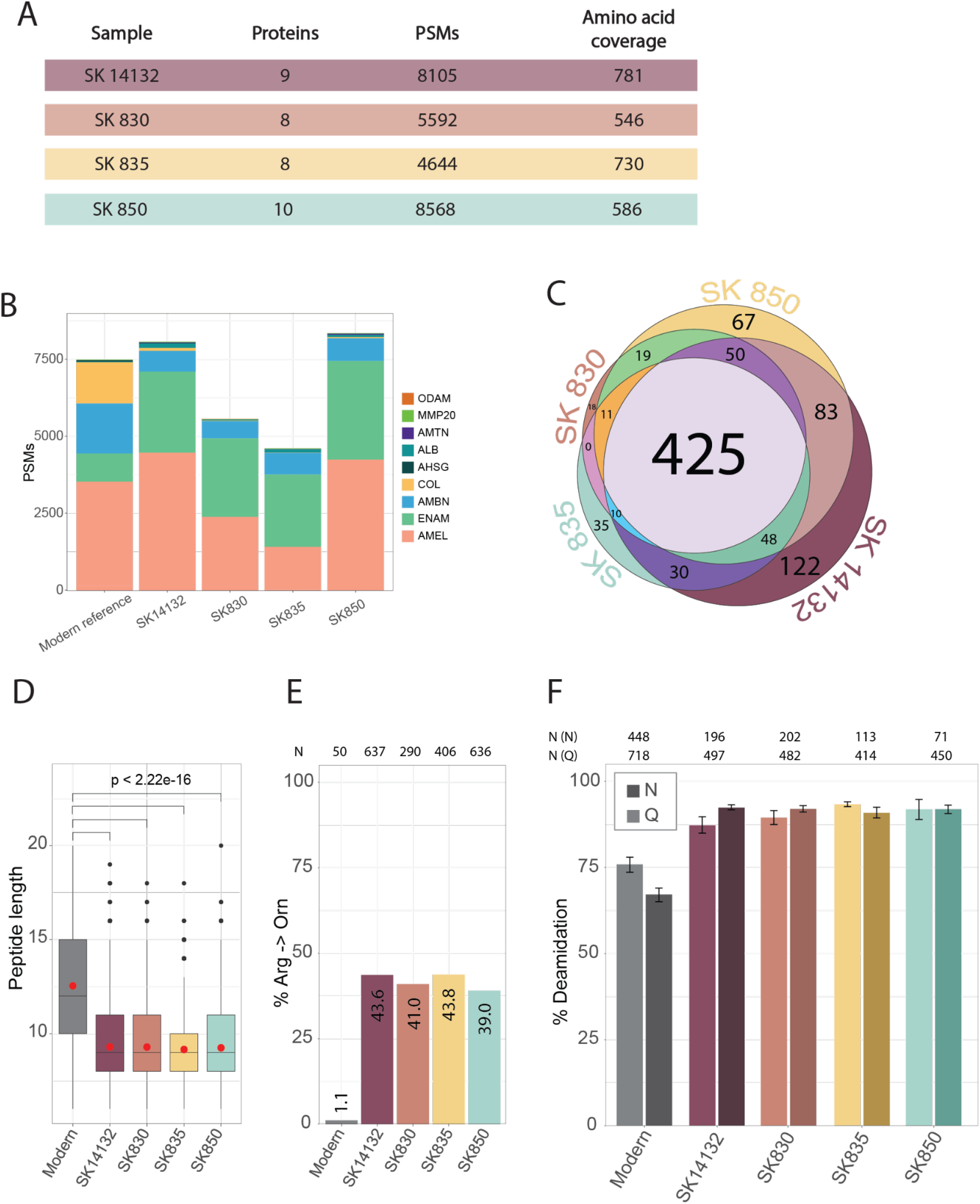
2-Ma recovered peptide sequences are consistent with ancient enamel material. A - Summary of the protein recovery of each *Paranthropus* sample. PSM = Peptide spectrum matches. B - PSM recovery per gene per sample shows consistent proteome composition between the *Paranthropus* samples and the modern human reference, corresponding to enamel. C - Venn diagram displaying the amino acid coverage per specimen, showing high overlap across samples. D - Diagenetic alterations lead to shorter peptides. Boxplot showing the peptide length distribution for modern *Homo sapiens* and *Paranthropus*. The red dot represents the mean. P = p-value. E - Arginine to ornithine conversion can attest to the peptide endogeneity. The modification rate was calculated based on PSM counts. N represents the number of PSMs containing an ornithine instead of an arginine per sample. F - Asparagine (N) and glutamine (Q) deamidation showed higher rates in the *Paranthropus* samples compared to the modern reference. The deamidation rate was calculated based on unique and razor peptides. N (N) and N (Q) represent the number of peptides used for calculation of asparagine and glutamine deamidation respectively. Error-bars are based on 1000 bootstrap replicates.

**Extended Data Fig. 2.**
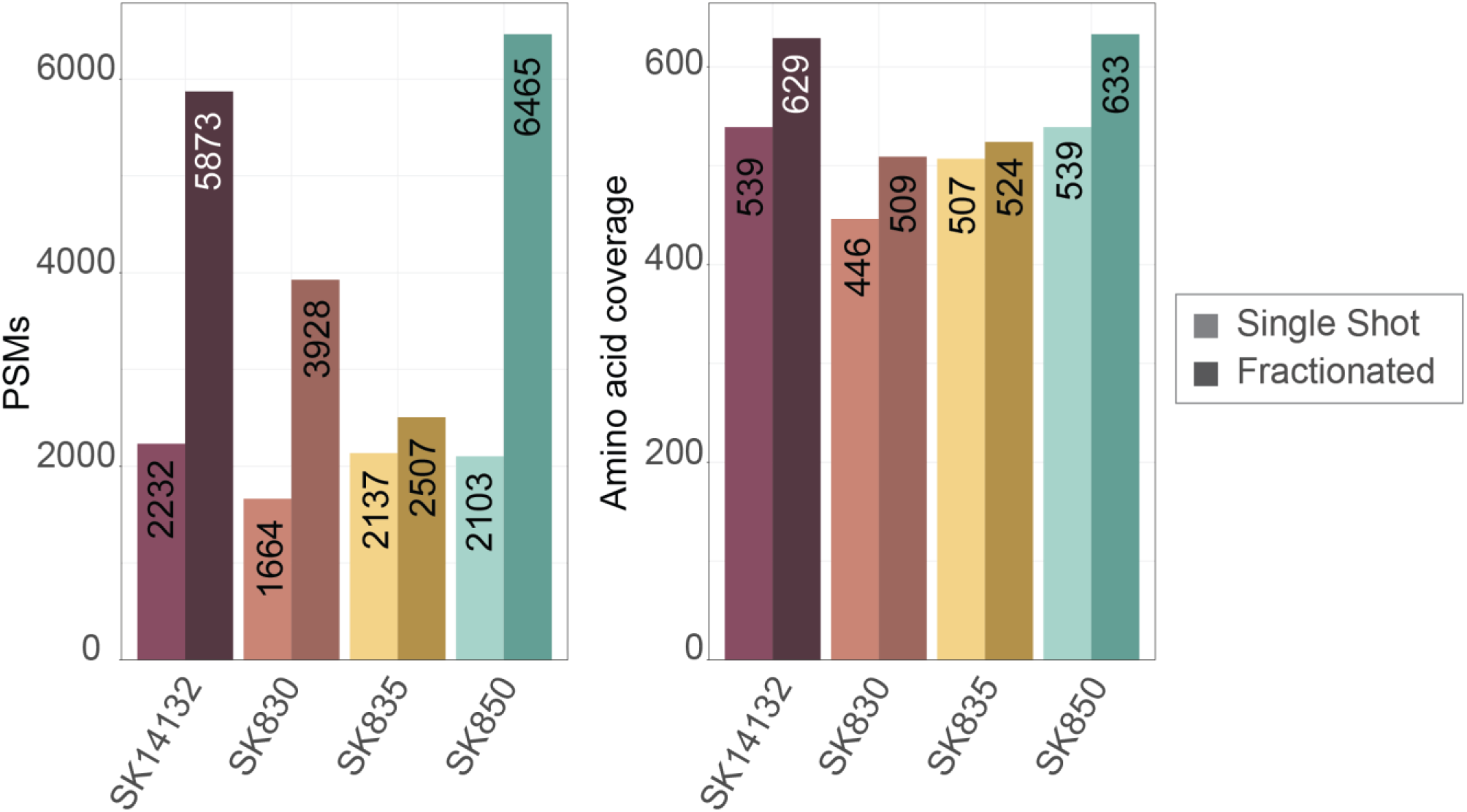
Fractionation enables increased coverage. PSM and amino acid coverage comparison between single shot analysis and fractionation for all four samples. The lighter colour displays the single-shot injection and the darker colour represents the fractionated samples.

**Extended Data Fig. 3.**
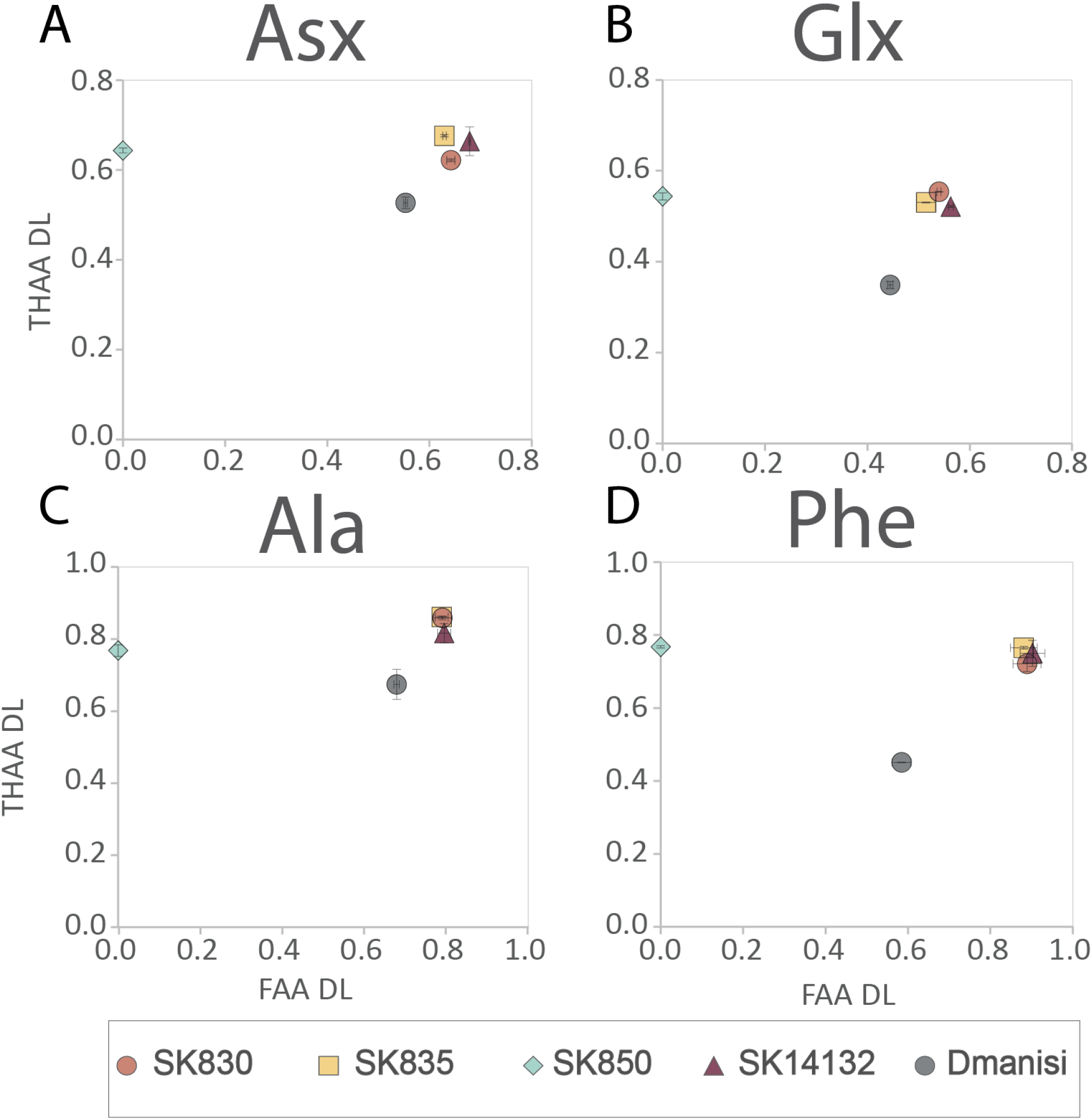
Amino acid racemisation analysis showed that the enamel displayed a closed system behaviour, supporting endogeneity of the peptides recovered by MS. Comparison of the *Paranthropus* enamel intra-crystalline FAA (free amino acids) vs THAA (total hydrolysable amino acids) covariance of four amino acids: A - asparagine/aspartic acid (Asx), B - glutamine/glutamic acid (Glx), C - alanine (Ala) and D - phenylalanine (Phe), to *H. erectus* enamel from Dmanisi. Note: The Swartkrans data point plotting on the y-axis is due to a lack of FAA data for that sample and does not represent D/L values of 0.

**Extended Data Fig. 4.**
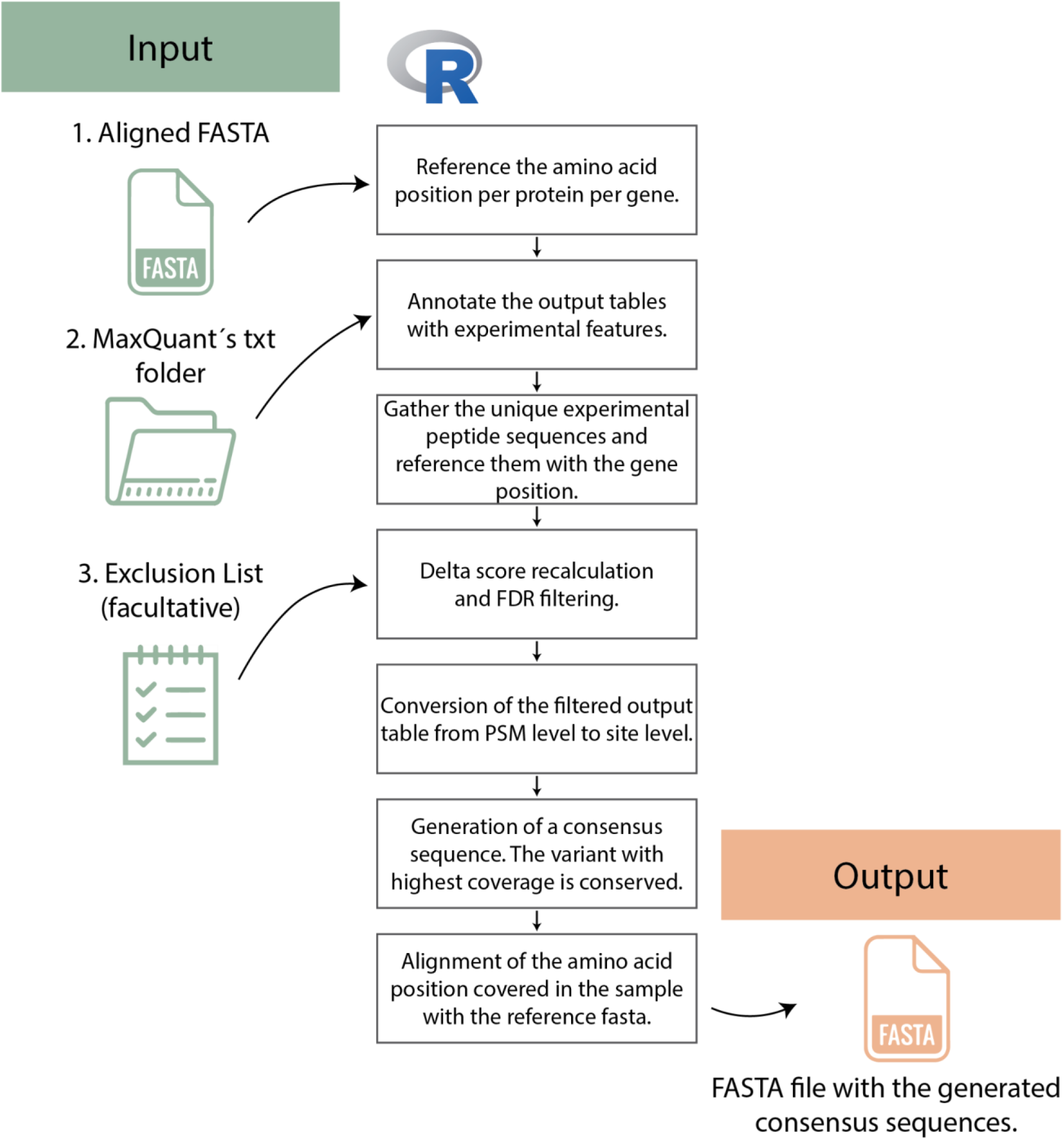
Semi-automated and transparent pipeline was developed for sequence reconstruction. Flow diagram of the sequence reconstruction pipeline. PSM = Peptide Spectrum Matches, FDR = False Discovery Rate.

**Extended Data Fig. 5.**
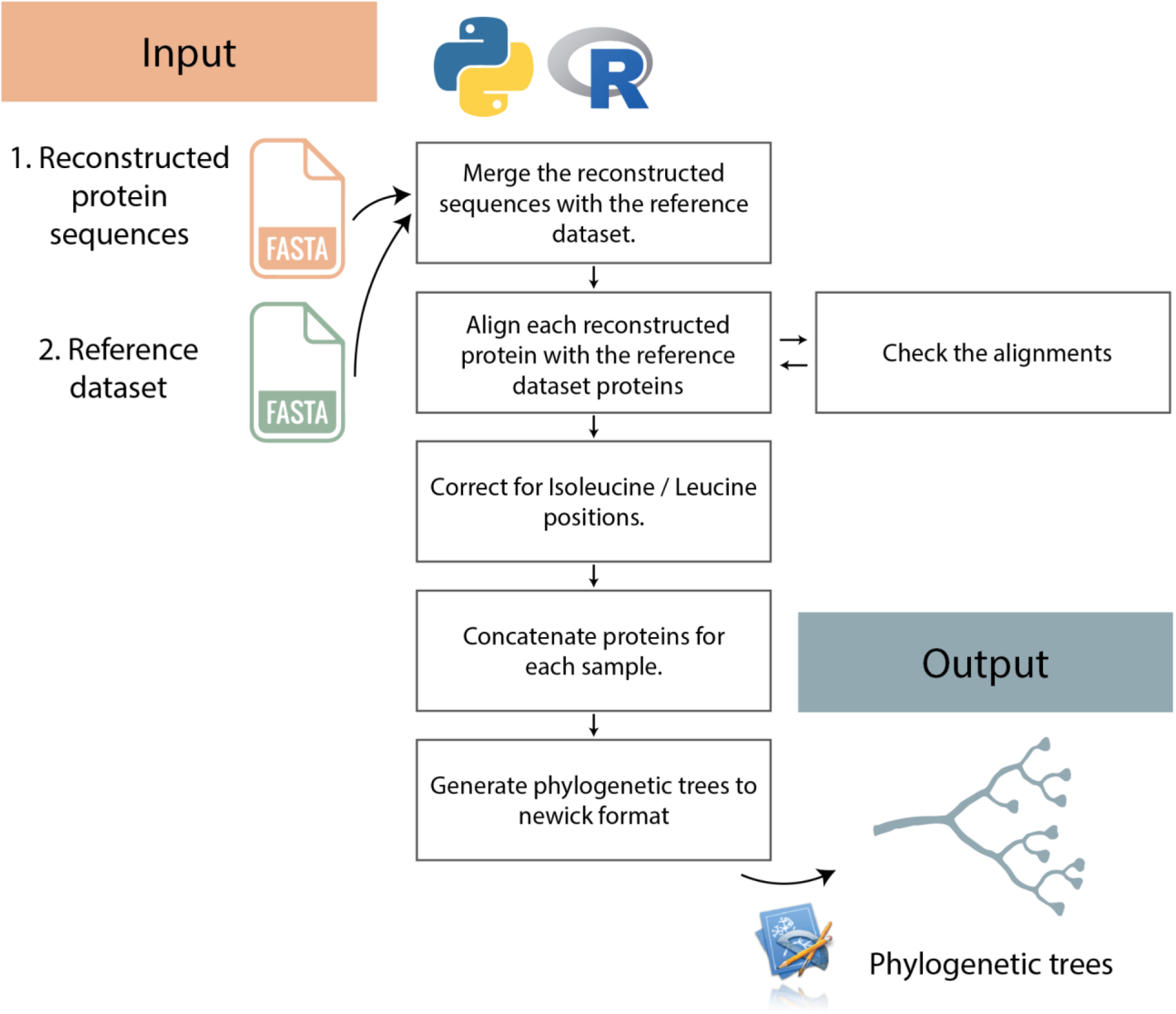
Semi-automated pipeline that was used for the phylogenetic analysis of the reconstructed sequences. Flow diagram of the sequence reconstruction pipeline.

**Extended Data Fig. 6.**
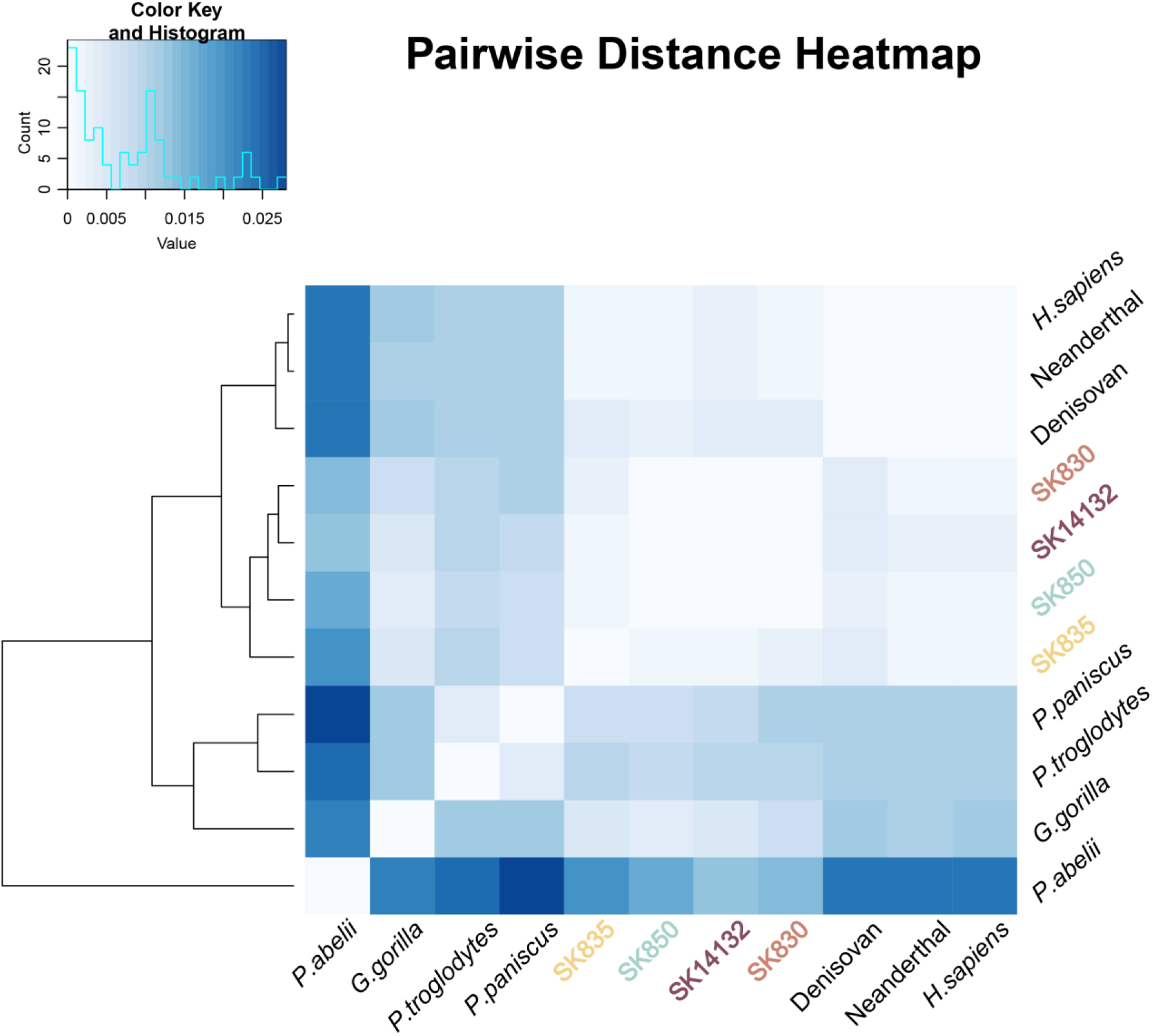
Heatmap of Pairwise distances. Heatmap of the pairwise distances matrix. Lighter colours depict lower distances.

**Extended Data Fig. 7.**
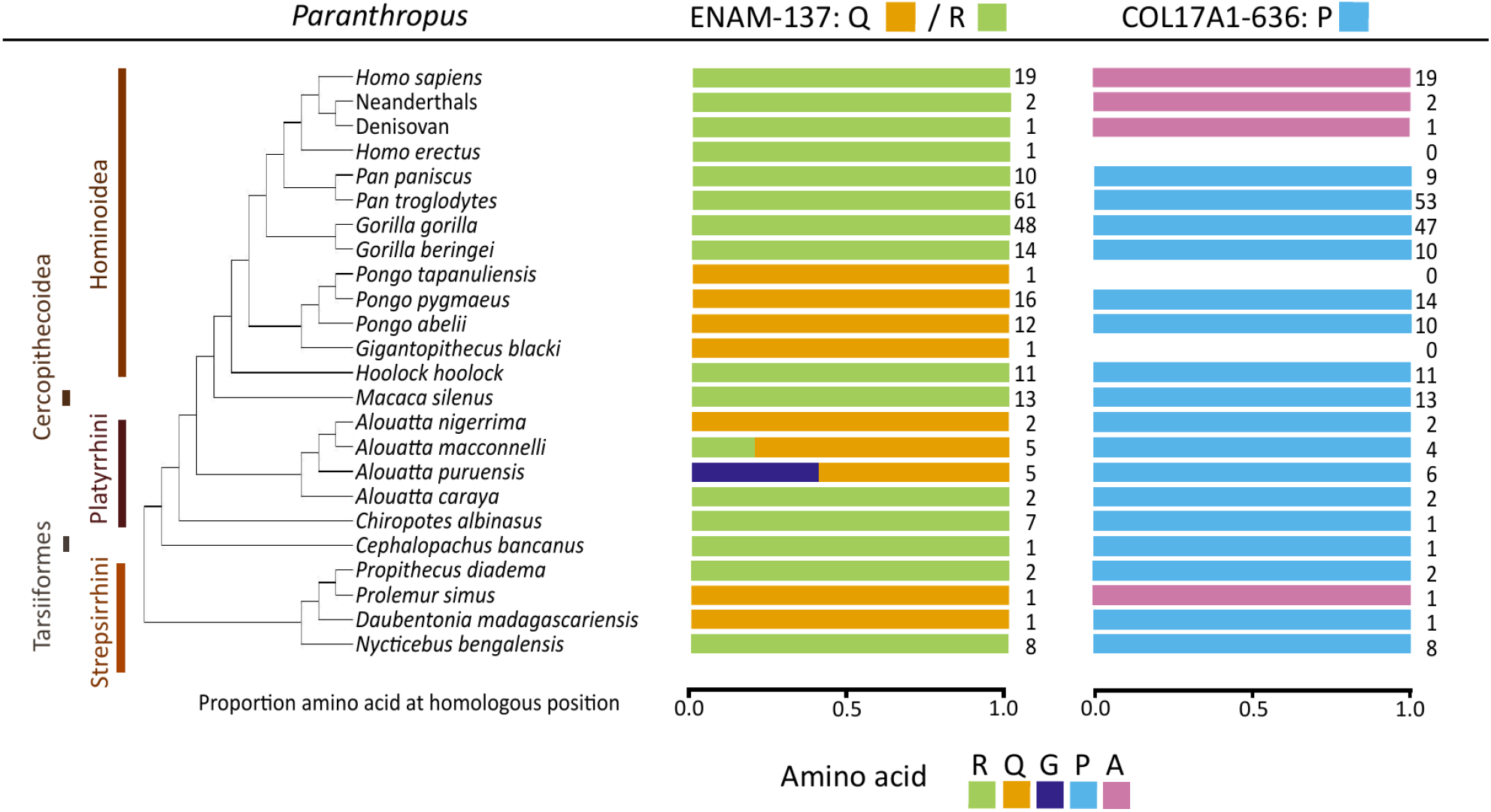
Homologous amino acids across primates. Amino acids at given positions in *Paranthropus* are indicated at top. Amino acids of other primates at homologous positions are shown below the black line and are organised in a cladogram. In ENAM, at positions homologous to position 137, most individuals across primates carry an Arginine (R). Glutamine (Q) is present in Ponginae, in some species of *Alouatta*, and in some species in Lemuroidea. In COL17A1, most samples carry a Proline (P) at positions homologous to position 636. Alanine (A) can only be observed in the genus *Homo* and in one individual of *Prolemur simus*. The number of individuals for which sequence data at the given position is available is indicated to the right of each bar.

**Extended Data Fig. 8.**
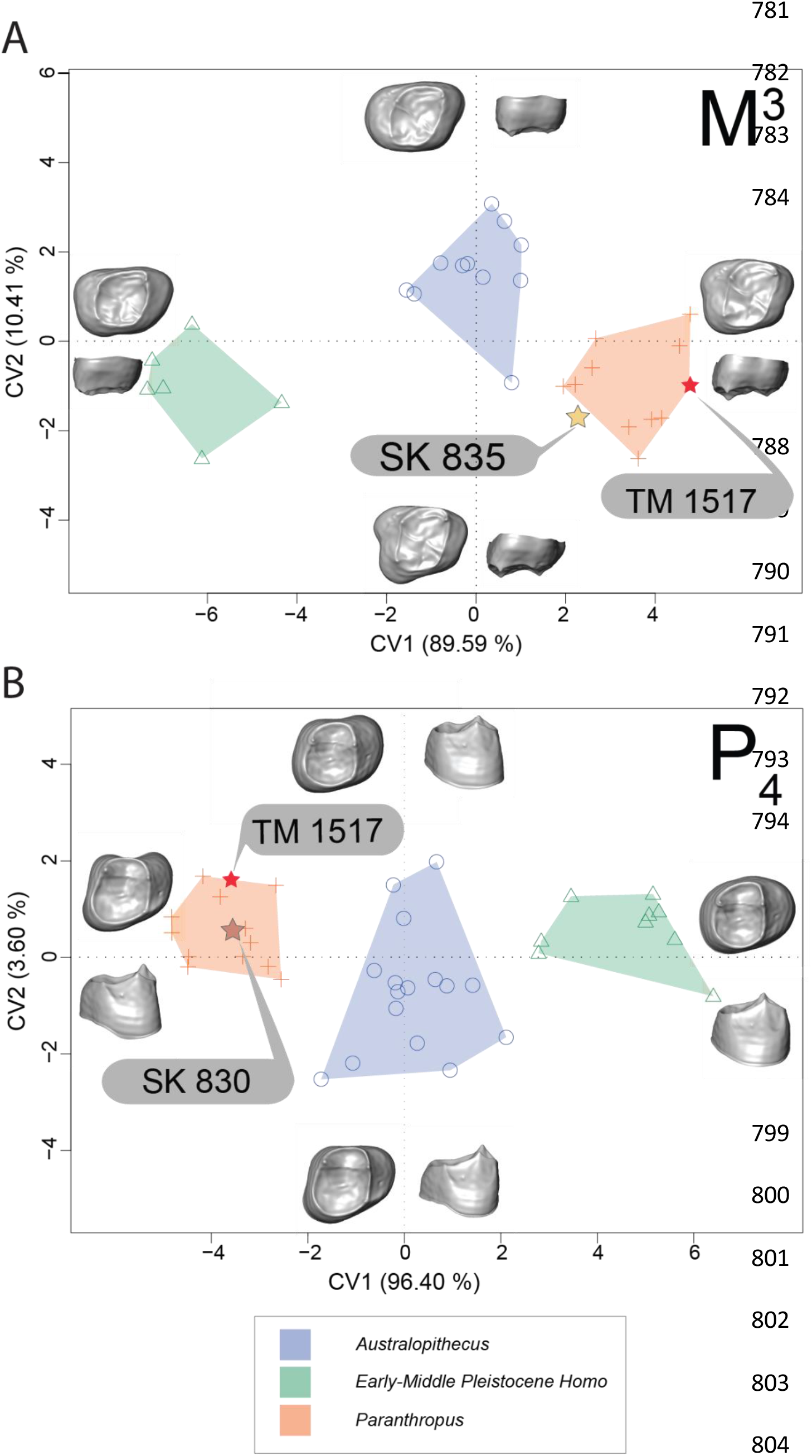
SK 850 and SK 835 fall within the *Paranthropus* morphospace. Bivariate plot of the canonical variates analysis (CVA) scores based on the diffeomorphic surface matching (DSM) deformation fields for the M^3^s (A) and P_4_s (B).

**Extended Data Fig. 9.**
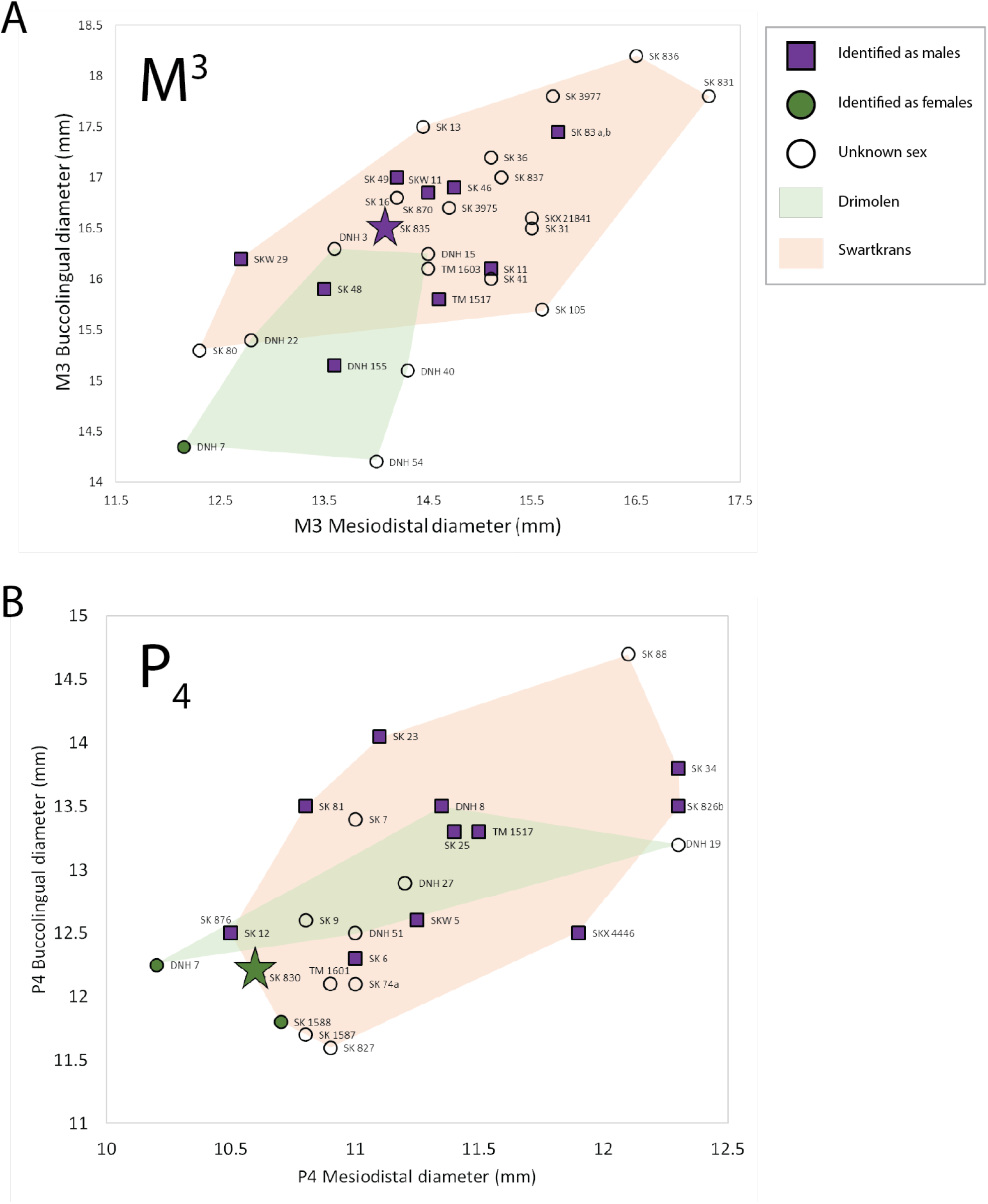
SK 835 crown size falls within Swartkrans but close to the Drimolen grouping. Bivariate plot of upper M3 (A) and lower P4 (B) standard length dimensions. Purple squares represent individuals identified as male in the literature, green circles represent individuals identified as female in the literature, and open circles are individuals of unknown sex. Specimens from Drimolen are grouped in the grey convex hull, and those from Swartkrans are grouped in the beige convex hull. A - Specimen, SK 835, identified as male in this study is denoted with a purple star. B - Specimen, SK 830, identified as female in this study is denoted with a green star.

**Extended Data Fig. 10.**
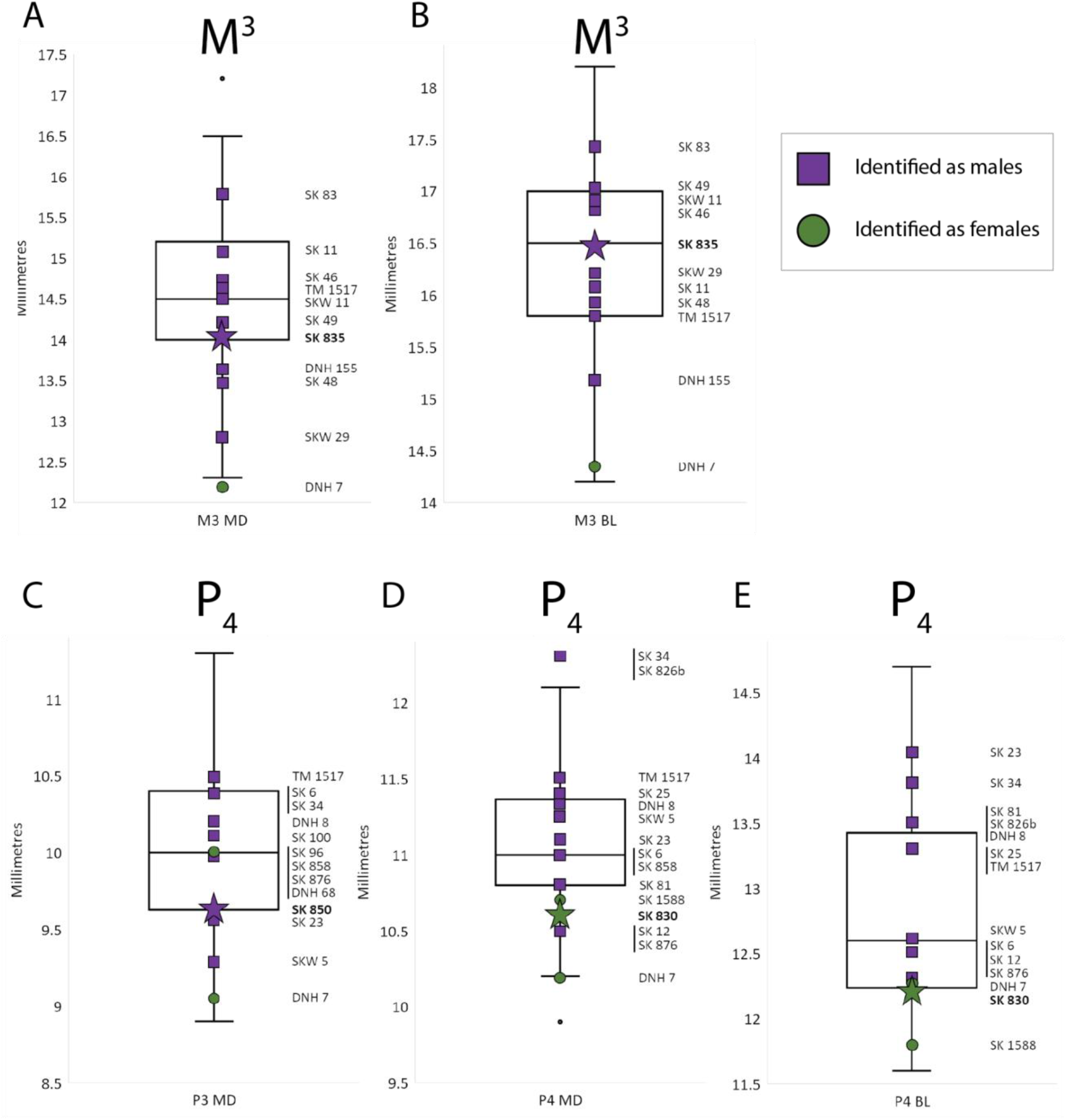
SK 850 MD measurements place it within the lower range of size variability. Boxplots showing the extent of size variation in M3 (A) and lower premolar (B) standard length dimensions. Purple squares represent individuals identified as male in the literature, and green circles represent individuals identified as female in the literature. These specimens are also labelled on the boxplot. A - Specimen SK 835, identified as male in this study is denoted with a purple star. B - Specimen SK 830, identified as female in this study is denoted with a green star, and specimen SK 850, identified as male in this study is denoted with a purple star.

