## Supplementary Information for "Enamel proteins reveal biological sex and genetic variability within southern African *Paranthropus*"

[1]Geogenetics Section, Globe institute, University of Copenhagen, Denmark, [2]Human Evolution Research Institute (HERI), University of Cape Town, Cape Town, South Africa, [3]Novo Nordisk Foundation Center for Protein Research, University of Copenhagen, Denmark, [4]Section for Molecular Ecology and Evolution, Globe Institute, University of Copenhagen, Denmark, [5]Department of Archaeology, University of Cape Town, Cape Town, South Africa, [6]Plio-Pleistocene Palaeontology Section, Ditsong National Museum of Natural History, Pretoria, South Africa, [7]Institute of Evolutionary Biology (UPF-CSIC), PRBB, Barcelona, Spain, [8] Turkana Basin Institute, Nairobi, Kenya, [9]Department of Geological Sciences, University of Cape Town, Cape Town South Africa, [10]Department of Chemistry, University of York, York, United Kingdom, [11]Palaeo-Research Institute, University of Johannesburg, Johannesburg, South Africa, [12]Department of Anthropology, University of Colorado Denver, Colorado, United States of America, [13]Palaeontology Section, National Museums of Kenya, Nairobi, Kenya, [14]D-CYPHR, Centre for Proteomic and Genomic Research, Cape Town, South Africa, [15]CBMN, UMR CNRS 5248, Proteome Platform, University of Bordeaux, Bordeaux, France, [16]Evolutionary Studies Institute, University of the Witwatersrand, Johannesburg, South Africa, [17]Computational Systems Biochemistry, Max Planck Institute of Biochemistry, Martinsried, Germany, [18]Department of Biological and Medical Psychology, University of Bergen, Bergen, Norway, [19]Research Center on Animal Cognition (CRCA), Center of Integrative Biology (CBI), University of Toulouse, CNRS UMR-5169, UPS, Toulouse, France, [20]Institute of Infectious Disease and Molecular Medicine, University of Cape Town, Cape Town, South Africa, [21]Section for Molecular Ecology and Evolution, Globe Institute, University of Copenhagen, Denmark, [22]Univ. Bordeaux, CNRS, MCC, PACEA, UMR 5199, F-33600 Pessac, France, [23]Department of Anthropology, University of Toronto Mississauga, Canada

### SUPPLEMENTARY MATERIALS

#### 1. The palaeoanthropological site of Swartkrans

##### 1.1. History

The UNESCO World Heritage Site, the Cradle of Humankind, in South Africa, known locally as The Cradle, is well known for its abundant hominin remains, ranging in age from just under 3 Ma to around 250 ka. The fossil hominins analysed here derive from the cave site of Swartkrans, located approximately 40 km northwest of Johannesburg, on the Blaauwbank River, in the province of Gauteng. Although Swartkrans cave was mined for lime in the 1930s, the first paleontological explorations of the Swartkrans breccias were only initiated by Robert Broom and John Robinson in 1948. The first hominins recovered from the site were assigned to *Paranthropus crassidens*, now recognised as a junior synonym of *P. robustus*. Remains of *Homo* were also recovered from Swartkrans, the first time these two genera were shown to be contemporaneous. Broom and Robinson's teams excavated at Swartkrans until 1951. Following Broom's death in 1951, Robinson continued to lead excavations for two more years, after which exploration ceased. Excavations restarted under the leadership of Bob Brain from the mid-1960s and continued to the mid-1980s. Collectively the excavations at Swartkrans led to a considerable sample of hominin, faunal and archaeological material, investigation of which continues to this day [1–4]. Brain demonstrated the existence of five members at Swartkrans [5]. The oldest hominin fossils are from Member 1, which dates to between 2.2 and 1.7 Ma (see below), with the youngest coming from Member 3. Other sites in South Africa of comparable age include Drimolen, Gondolin and Kromdraai [6–9].

##### 1.2. Cave geology, stratigraphy and dating

All the caves in the Cradle preserve two basic rock types: sediments, in which the fossils are preserved, and cave carbonates. These carbonates, in this case flowstones, are ubiquitous features at all the cave sites in the Cradle. These deposits provide ages, via U-Pb dating, for the layers of sediments and the fossils preserved in them, and in ideal case, these can be both maximum and minimum ages. A regional summation of all the available U-Pb ages for the Cradle indicates that flowstone growth is not continuous but episodic, with six discrete time windows of formation between 3.2 and 1.3 Ma [9].

At Swartkrans both flowstones and cave sediments (previously known as breccia) are preserved and are divided into a Member system, as follows [10]. The *Paranthropus* teeth analysed here are all from Member 1.

**Member 1** consists of two units, the Hanging Remnant and Lower Bank, both of which have yielded *Paranthropus* and *Homo* remains [10]. Direct ages for the flowstone layers below both sections of Member 1 are indistinguishable from each other at 2.248 ± 0.052 Ma (Hanging Remnant) and 2.249 ± 0.077 Ma (Lower Bank), and mean the fossil bearing sediments over these layers are younger than this [9,11]. These sediments have been dated by cosmogenic nuclide burial dating, which yielded age estimates of 2.19 ±0.08 and 1.80 ± 0.09 Ma respectively [12]. The flowstones capping both sections of Member 1 have U-Pb age estimates of 1.800 ± 0.005 Ma (Hanging Remnant) and 1.706 ± 0.069 (Lower Bank). Although consistent with the estimated age of the sediment, these datings also indicate that the biochronological ages are underestimated [9,13], as are the ESR ages from the Hanging Remnant at 1.6 Ma [14].

**Member 2** falls chronologically between Members 1 and 3, and has yielded fossils, stone tools and bone artifacts [10]. The U-Pb date 1.706 ± 0.069 Ma for one of the flowstones underlying Member 2 provides a maximum age for the deposits [13].
**Member 3** has been dated to 0.96 ± 0.09 Ma [12] and records an increased number of macrovertebrate taxa, with a recovery of 54 taxa.

### 2. *Paranthropus* sample

#### 2.1 History and taxonomy

The taxon *Paranthropus robustus* was established in 1938 based on the specimen TM 1517 to describe a collection of "robust" fossil hominins found at the site[1] of Kromdraai B in the Cradle of Humankind, South Africa [15]. This is the type species of the genus *Paranthropus*, which encompasses two additional hominin species, *P. aethiopicus* and *P. boisei* in eastern Africa. Collectively these taxa display highly derived morphological features, such as large sagittal crests, relatively massive jaws, "anterior pillars" of the face, and postcanine megadontia coupled with thick enamel, possibly related to a functional adaptation to a heavy masticatory load [16,17]. *Paranthropus* spans a large geographic range across Africa, with South African *P. robustus* representing the southernmost known geographical limit, and cranial and mandibular remains of *P. boisei* from Konso, southern Ethiopia [18], representing the northernmost known geographical limit. Since the establishment of *P. robustus*, specimens attributed to this species have been recovered from the South African cave deposits of Kromdraai Member 3, Drimolen, Swartkrans Members 1-3, Coopers and Gondolin, and have been dated to between 2.21 and 1.07 million years ago [19,20].

The phylogenetic relationships within and between the three known species of *Paranthropus* have been the subject of considerable discussion. While most researchers consider *Paranthropus* taxa to be more closely related to each other than to other hominin species (i.e. a monophyletic group [21]), other researchers point to a number of morphological similarities between *P. robustus* and *Au. africanus* in a South African context [22–24], and between *P. aethiopicus* and *Au. afarensis* in an eastern African context [25–27], raising the possibility of paraphyly.

#### 2.2 Sexual dimorphism

There is considerable size variation within *P. robustus*, most of which has previously been attributed to sexual dimorphism, and also, for males, possibly reflecting a gorilla-like pattern of extended growth (bimaturation)[28]. High levels of sexual dimorphism correlate with higher levels of male-male competition in primates, and with male dominance over multiple females [29]. It is assumed, given patterns seen in living apes, and to a lesser extent humans, that bigger individuals are males, and smaller ones' females [30], though sexing on the basis of size alone is problematic [31]. This has implications for our understanding of taxonomy. As one example, a morphological analysis of a *Paranthropus* tooth discovered at the site of Gondolin, the largest of its kind in South Africa, tentatively concluded that this specimen was from a very large *P. robustus* male [32], which would indicate that the extent of sexual dimorphism in this species was previously underestimated. However, other possibilities include that the tooth could represent the first evidence of *P. boisei* in South Africa, or could represent a "robust australopithecine" species that is yet to be described [33].

##### 2.3. Analysed *Paranthropus* teeth

Following successful recovery of palaeoproteins from fauna (Supplementary Information 3), *Paranthropus* material was then sampled. All four sampled *Paranthropus* teeth come from the site of Swartkrans, and are housed at Ditsong Museum, South Africa.

**SK 830** was collected in 1952 from the “Lower breccia”, now recognised as Member 1. It represents an isolated left $P_4$, slightly worn, with a portion of the crown missing and damaged roots.

**SK 850** was collected in 1949, also from the “Lower breccia”, and represents an isolated, significantly worn right $P_3$ crown and root system, both incomplete.

**SK 835** is a left isolated $M^3$, recovered in 1956 from Member 1, and consists of three separate crown fragments and three separate root fragments, reconstructed by [34].

**SK 14132** was collected in later excavations from the Member 1 Hanging Remnant [35], and is an isolated right $M^3$, consisting of the buccal side of the crown with part of the root [35].

#### 3. Faunal samples

Prior to analysis of hominin material, three bovid teeth from the site of Swartkrans, and one bovid tooth from Coopers Cave, South Africa, were sampled for palaeoproteomic analysis, in order to determine the feasibility of protein recovery. Swartkrans is described above. Coopers Cave is located approximately 45 km southwest of Johannesburg between Sterkfontein and Kromdraai fossil sites, and has yielded a partial cranium of *P. robustus* [36]. The site presents three distinct collapsed and deroofed breccia deposits labelled Coopers A, B and D [3]. The faunal material is housed at the University of the Witwatersrand in South Africa. SKX 3730, SKX 4996 a, and SKX 37041 were all found in Swartkrans from Members 1-3, where *Paranthropus* material has also been found. CD.5410 was discovered in Cooper’s Cave Locality D, where a *Paranthropus* RM2 had been retrieved with missing roots with the enamel-dentine junction (EDJ) worn away.

**SKX 4996a**, a left maxilla fragment with M1-M3, attributed to *Connochaetes* sp., derived from Member 1 of the Swartkrans Formation [10].

**SKX 3730**, an isolated upper permanent tooth assigned to *Connochaetes* sp. was recovered from Member 2, where a total of 14 macrovertebrate taxa have been recovered from this infill [10].

**SKX 37041**, an upper permanent tooth is attributed to *Damaliscus* sp. from Member 3. This infill is associated with controlled use of fire though no *Homo* sp. have been recovered from the deposit.

**CD.5420**, a lower left second molar that had originally been attributed to Kudu (*Tragelaphus strepsiceros*), but Hanon et al. [37] argues could belong to eland (*Tragelaphus oryx*). The specimen was recovered from the decalcified sediments of Cooper’s D infill. The flowstone at the base of this sequence has a U-Pb date of 1.37 Ma ± 0.113 [38]. There is no capping flowstone at this site but based on the regional patterns of flowstone/sediment intervals in the Cradle, the fossils are most likely between 1.3 and 1.1 Ma.

### SUPPLEMENTARY METHODS

#### 4. Biomolecular preservation: Intra-crystalline protein degradation analyses (IcPD)

Chiral amino acid analysis was undertaken on the four *Paranthropus* teeth from Swartkrans. Three of the four samples were sent as coarse powders; however, SK 835 contained one chip large enough to view under a microscope. The chip contained two different mineral components: one white, likely to be enamel, and the other creamier, suspected to be dentine (Figure S1). A precision drill was used to isolate the enamel for the analysis.

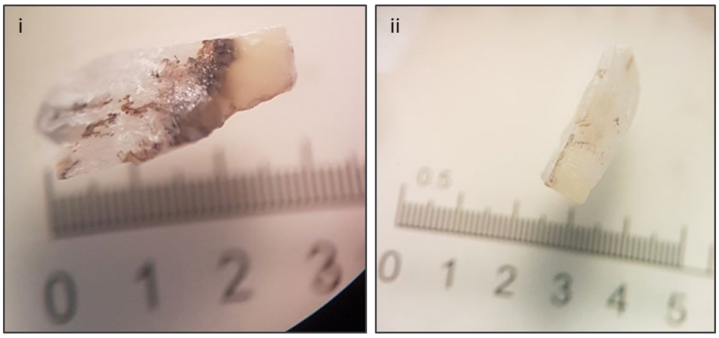


**Figure S1. Microscope images of a small chip of dental material from tooth SK 835 both before (i) and after (ii) cleaning with a precision drill.** Chip (i) shows some signs of black staining, probably iron introduced during taphonomy. Chip (ii) contains a whiter and a creamier component, the creamier component is likely dentine. The suspected dentine as well as the black staining was removed with a precision drill prior to IcPD analysis. Note: scale units in mm.

The enamel samples were powdered with an agate pestle and mortar. All samples were prepared using modified procedures of Penkman et al.[39], but optimised for enamel, using a bleach time of 72 hours to isolate the intra-crystalline protein [40]. Two subsamples were analysed from each portion: one fraction was directly demineralised and the free amino acids analysed (referred to as the 'free' amino acids, FAA, F), and the second was treated to release the peptide-bound amino acids, thus yielding the 'total hydrolysable' amino acid fraction (THAA, H). However, due to the small masses of samples there was not always sufficient material to analyse both fractions. Therefore it was not possible to prepare an FAA subsample for SK 850. After demineralisation, the pH of the solution was raised with potassium hydroxide (KOH) followed by centrifugation for 5 min at 13,000 rpm, whereupon a biphasic solution was formed[40]. The supernatant was extracted and dried. Samples were analysed in duplicate by RP-HPLC, with standards and blanks run alongside samples.

### 5. Proteomics

#### 5.1. Faunal analysis

Enamel proteins were extracted using 10% trifluoroacetic acid (TFA), following the method detailed in Cappellini et al. (2019). Each experiment was injected separately in the mass spectrometer. Laboratory blank extraction was also used to monitor possible contamination. Data was acquired and processed with the same parameters described below for the analysis of the *Paranthropus* samples. In short, enamel peptides were immobilised on C18 Stage-Tips and analysed by nano-liquid chromatography coupled with tandem mass spectrometry (nanoLC-MS/MS). The MS/MS spectra were identified using MaxQuant (V. 1.6.0.17), matching them against a custom built reference database containing all the publicly available sequences for enamel proteins for modern Bovidae species (210 entries). Protein sequences were assembled using the same R based data analysis pipeline as for the *Paranthropus* samples.

#### 5.2. Peptide extraction, clean-up and fractionation:

Enamel was cleaned and separated from dentine with a hand held dremel saw. Peptide extraction was performed following Cappellini et al[41]. In short, 80-250 mg of powdered enamel per tooth (total over all extractions) was demineralized twice overnight at 4 °C in 10% TFA.

In-house stage-tips were prepared using low-bind 200 µL tips and three discs of 3 M Empore $C_{18}$ (Thermo Fisher Scientific). Stage-tips were conditioned with 150 µL methanol, 150 µL AT80 (0.1% TFA in 80% acetonitrile (ACN)/20% pure water) and equilibrated with 150 µL 0.1% TFA. The peptides were loaded on the stage-tip and salts were removed with 150 µL of 0.1% TFA in water[41]. The stage-tips were either directly completely eluted for single-shot LC-MS/MS analysis or eluted into fractions using high-pH reversed-phase offline fractionation.
For the single shot analysis, the peptides were eluted from the stage-tips using 30 µL 40% ACN into a 96 well plate. ACN was evaporated at 45°C to approximately 3 µL and the peptides were resuspended in 10 µL 0.1 % TFA, 5% ACN for the initial faunal samples, and 20 µL for the initial single shot analyses of *Paranthropus* (with 10 µL kept in the freezer for security)..

60-200 mg of enamel were used for offline fractionation. Buffer A was prepared with sterile-filtered 5 mM ammonium bicarbonate (ABC) (pH 7.8) and buffer B was 100 % ACN. Peptides were eluted in fractions of 2 x 20 µL sequentially with an increasing percentage of buffer B at 0 % (pH adjustment fraction), 11 %, 15 %, 20 %, 70 % of buffer B. The fractions were acidified with 4 µL of 10 % TFA and ACN was evaporated at 45°C. For subsequent MS analysis using the nano-LC system, the peptides from samples SK 830 and SK 835 were resuspended in 0.1 % TFA, 5% ACN. For MS analysis using an EvosepOne chromatographic system, the peptides from samples SK 850 and SK 14132 were resuspended in 20 µL 0.1% FA and loaded onto Evotips.

#### 5.3. LC-MS/MS analysis

##### 5.3.1. Copenhagen (CPH)

Peptides were separated on a liquid chromatographic system (EASY-nLC™ 1200 or Evosep One) coupled to an Exploris 480 mass spectrometer (Thermo Scientific, Bremen, Germany). Peptides were separated using an in-house packed column (15 cm x 75 µm, 1.9 µm) packed with $C_{18}$ beads (Reprosil-AQ Pur, Dr. Maisch). The column temperature was maintained at 40°C, using an integrated column oven (PRSO-V1, Sonation).

The nanoLC was operated using a 77 min gradient, ranging linearly from 5 % to 30 % B (80% ACN, 0.1 % formic acid (FA)) in 50 min, 30% to 45 % B in 10 min, 45 % to 80 % B in 2 min, maintained for 5 min before dropping back to the initial conditions in 5 min and equilibrated for 5 more min. Buffer A was milliQ water and the flow rate was set at 250 nL/min. For the initial faunal samples and single shot analyses, 5 µL were injected.

The separation on the Evosep was performed using the predefined commercial gradient 20 SPD, operating at a flow rate of 200 nL/min.

The mass spectrometer was operated in Data Dependent Acquisition (DDA) in positive mode. Spray voltage was set to 2.0 kV, heated capillary temperature at 275°C and funnel RF level at 40. Data was acquired in profile mode. MS1 mass range was set at 350–1400 with an AGC target at 300%. Full MS resolution was set at 120,000 at m/z 200 with a maximum injection time of 25 ms. The HCD fragment spectra resolution was set at 60,000 with a maximum injection time of 118 ms and a Top10 method with a 30 seconds dynamic exclusion. AGC target value was set at 200 % and the intensity threshold at 2e5. The isolation window was set at 1.2 m/z and the normalised collision energy at 30%.

Faunal samples were analysed on a Q Exactive HF-X using the same parameters as described above, except for the normalised collision energy that was set at 28%.

Furthermore, injection blanks were run prior to every sample injection to limit the risk of carry over in the chromatographic column.

###### 5.3.2. South Africa (SA)

Peptides were separated on an EvoSep One system using the 20 SPD gradient described above. The peptides were analysed in a Q-Exactive mass spectrometer, operated in DDA mode. Spray voltage was set to 1.95 kV, heated capillary temperature at 320°C and funnel RF level at 50. Data was acquired in profile mode. MS1 mass range was set at 350–1400 with an AGC target at 3e6. Full MS resolution was set at 70,000 at m/z 200 with a maximum injection time of 240 ms. The HCD fragment spectra resolution was set at 35,000 with a maximum injection time of 110 ms and a Top10 method. AGC target value was set at 2e5. The isolation window was set at 2.0 m/z and the normalised collision energy at 27%.

##### 5.4. Synthetic peptides

KPPEKEPLK, {pGlu}K{Orn}PLKEP and KPPEK{Orn}PLK were purchased from Genscript (purity > 98%). The peptides were solubilized in 50 % ACN, 0.1 % TFA and were dried at 45°C before being stored at -80°C.

Peptides were resuspended in 0.1 % FA. 250 fmol of each peptide was injected for MS analysis. An inclusion list with the 3 peptides at both charge 2 and 3 was generated (Table S1). A survey scan with that inclusion list was run using the same parameters (ie. gradient, collision energy, resolution) as the ones used for the *Paranthropus* samples was carried out. MS2 spectra at the APEX of the chromatographic peaks were extracted. Spectra were compared to the experimental spectra acquired in the *Paranthropus* samples using the Universal Spectrum Explorer[42].

**Table S1: Synthetic peptides inclusion list**. Peptide sequence, charge and mass to charge ratio of the three synthetic peptides covering ENAM-137 used for the survey scan.

| Peptide | Charge | m/z |
|---|---|---|
| KPPEKEPLK | 2 | 533.3188 |
| KPPEKEPLK | 3 | 355.8816 |
| {pGlu}K{Orn}PLKEP | 2 | 468.7793 |
| {pGlu}K{Orn}PLKEP | 3 | 312.8553 |
| KPPEK{Orn}PLK | 2 | 525.8371 |
| KPPEK{Orn}PLK | 3 | 350.8938 |

##### 5.5. Data search strategy

Raw data were searched using MaxQuant (V. 1.6.0.17)[43], pFind (V. 3.1.5)[44] and Peaks Client 7.0 (V 7.5). A custom-built database containing 622 entries was used, and contained enamel specific proteins collected from Uniprot corresponding to human, chimpanzee, gorilla and orangutan. This database was supplemented with the *Gigantopithecus blacki*, *Homo antecessor*, Denisovan enamel proteome and some Neanderthal enamel specific sequences [45,46].

##### 5.5.1 MaxQuant search

The raw files were searched against the custom-built database (**622 entries**), supplemented with a database of contaminants (**110 entries**). Unspecific digestion was selected. Cysteine trioxidation was set as a fixed modification, and Arg->Ornithine, Gln->pyro-Glu, Glu->pyro-Glu, Deamidation (NQ), Phospho (ST) and Oxidation (MPW) were set as variable modifications. The minimum peptide length was set at 6 amino acids for unspecific search and the maximum was set at 20 amino acids. The peptide mass was limited to 3500 Da. The minimum Andromeda score for both modified and unmodified peptides was fixed at 25. Protein, PSM and site FDR were set at 10% and were manually adjusted during the data analysis. The minimum delta score for modified and unmodified peptides was set at 0 and was manually adjusted afterwards (Supplementary Information 9.4). Both iBAQ and Dependent Peptides were enabled.

The *Paranthropus* raw files were searched with the corresponding lab blanks, modern *Homo sapiens, Gorilla* and *Pongo* reference files, as well as raw files corresponding to *Homo antecessor* and *Gigantopithecus blacki* samples[45,46]. This strategy enabled us to directly compare identifications of endogenous peptides from the *Paranthropus* samples with reference spectra from other species and modern material. Moreover, it was also directly possible to compare the modification levels of the *Paranthropus* samples with modern human reference files.

##### 5.5.2. Open Search strategy with pFind

Each *Paranthropus* individual was searched separately using pFind[47]. Asparagine and Glutamine deamidation were set as fixed modifications. Serine phosphorylation, arginine to ornithine conversion, glutamine and glutamic acid conversion to pyroglutamate and methionine, proline and tryptophan oxidation were set as variable modifications. The raw files were searched against the previously described database (**622 entries**), using unspecific digestion. FDR was set at 1%.

##### 5.5.3. PEAKS

We employed the Peaks Clients 7.0 (v.7.5) Spider and Peaks PTM algorithms to precisely identify peptide sequences in all *Paranthropus* individuals by matching them against the mentioned database through an unspecific search. We designated the following modifications as variables: Pyro-glu from E, Pyro-glu from Q, Deamidation (NQ), Oxidation (M), Phosphorylation (STY), Oxidation (HPW), Ornithine from Arginine, and Oxidation to nitro (Y). To guarantee the accuracy of the peptide matches, we set the FDR to 1% and the de novo ALC score to 80%.

#### 5.6. Modification analysis

The raw files were searched again using MaxQuant (V. 1.6.0.17) for low abundant PTM analysis. Besides the variable modifications parameter, the search parameters were kept identical. 5 searches were carried with the most abundant modifications set as variable modifications: deamidation of asparagine (ΔM=+0.984016 Da) and glutamine (ΔM=+0.984016 Da), serine and threonine phosphorylation (ΔM=+79.966331 Da), N-terminal glutamine and glutamic acid conversion to pyroglutamate (ΔM=-17.026549 Da and ΔM=-18.010565 Da respectively) and methionine oxidation (ΔM=+15.9949 Da). Additionally a group of modifications affecting the same amino acid were added to the search (Table S2).

**Table S2: Search iteration for PTM analysis.** Table summing up the different searches carried out for the analysis of PTMs.

| Search | Amino acid | Modification | ΔM (Da) |
|---|---|---|---|
| 5 | R | Arginine to ornithine conversion | -42.021798 |
| 4 | W | Tryptophan oxidation<br>Tryptophan dioxidation<br>Kynurenine<br>Tryptophan oxolactone<br>Tryptophandione | +15.9949<br>+31.990<br>+3.994915<br>+13.979265<br>+29.974178 |
| 3 | F | Phenylalanine oxidation<br>Phenylalanine dioxidation | +15.9949<br>+31.990 |
| 2 | Y | Tyrosine oxidation<br>Tyrosine dioxidation | +15.9949<br>+31.990 |
| 1 | H | Oxohistidine<br>Dioxohistidine<br>Hydroxyglutamate<br>Histidine to aspartic acid | +15.9949<br>+31.990<br>+7.979<br>-22.032 |

The extent of deamidation was assessed using the method described in Mackie et al [48]. Other modifications were investigated by PSM counting. Briefly, the ratio between the number of amino acids that could be affected by a given modification, to the number of amino acids actually modified was calculated. The counts were normalised by the MS count.

#### 5.7. Cross-linking analysis in ancient enamel

Raw data were searched manually and using the following softwares: MaxLynx [49] (MaxQuant, Germany), Mass Spec Studio [50] (CRIMP, version 2.4.0.3545, Canada) and XlinkX [51] (ThermoFisher Scientific). No chemical crosslinkers were used in this study. Cross-link searches were performed using FASTA files containing the sequences for ENAM, ABMN and AMELX including the amino acid substitutions identified from the sequence reconstruction analysis. Search parameters included; non-specific digestion, MS1/MS2 error tolerances at 10 ppm, oxidation of methionine as a variable modification. Cross-links that were searched for include; degradation related: W-W (-2H); Y-Y (-2H); H-K (-2H, +1O); disulfide bonds: C-C (-2H); enzymatic: K-N (-N, -3H). Any cross-links identified from the bioinformatic tools were then verified manually. In addition to this, data were searched manually for highly charged peptides (4+, 5+ and above) that could indicate the presence of cross-linked peptides.

### 6. Phylogenetic and variation analysis of *Paranthropus* specimens

#### 6.1. Protein Reference Datasets

The reconstructed *Paranthropus* sequences were analysed using three protein reference datasets. Details on which samples are contained within each of the datasets are given below.

##### 6.1.1 'Diversity' Dataset

The 'Diversity' dataset is composed of the highest number of individuals (217) and consists mostly of in silico translated protein sequences from the 4 genera of Hominidae. The translated sequences cover the following species: *Pongo abelii, Pongo pygmaeus, Pongo tapanuliensis, Gorilla gorilla, Pan troglodytes, Pan paniscus, Homo sapiens*. Additionally the dataset contains 3 Neanderthal individuals and 1 Denisovan, translated from high coverage ancient genomes. The sequences for these samples were acquired from the 'Hominid Palaeoproteomic Reference Dataset' which is publicly available at Zenodo, under the DOI : 10.5281/zenodo.7728060 (https://zenodo.org/record/7728060#.ZBMOGXbMKbg).

The reference sequences of *Macaca mulatta, Macaca nemestrina* and *Nomascus leucogenys* were also added from Ensembl to be utilised as outgroups for the analysis.

##### 6.1.2 'Representative' Dataset

The 'Representative' dataset contains only a small number of individuals (n=13). These include the following entries:

- a single representative individual for the *Pongo* genus (1 *Pongo abelii*)
- a single individual for the *Gorilla* genus (1 *Gorilla gorilla*)
- 2 individuals for the *Pan* genus (1 *Pan troglodytes* and 1 *Pan paniscus*)
- 3 individuals for the genus *Homo* (1 modern human, 1 Neanderthal and 1 Denisovan)

With the exception of the Neanderthal and Denisovan individuals, the sequences for the rest of the individuals were obtained from the reference proteins available from Ensembl [52]. The exact Ensembl ID codes for each entry are provided below (Table S3). For some species the reference protein sequences were missing. In these cases a random individual of the same species was selected from the in-silico translated samples of the 'Diversity Dataset' and provided the missing proteins. Out of the three high coverage Neanderthal samples, the Altai Neanderthal individual, sequenced in [53] was chosen as the Neanderthal representative. Similarly to the 'Diversity' dataset, the reference sequences of *Macaca mulatta, Macaca nemestrina* and *Nomascus leucogenys* were added from Ensembl and functioned as outgroups in the analysis.

**Table S3: Construction of the `Representative' dataset.** Accession numbers of reference sequences obtained from Ensembl, used for the 'Representative' dataset

| Species name | Accession numbers for the sequences used |
|---|---|
| *Homo sapiens* | ENST00000411641.7,ENST00000295897.9,ENST00000322937.10,ENST00000651267.2,ENST00000380712.7,ENST00000339336.9,ENST00000225964.10,ENST00000648076.2,ENST00000396073.4,ENST00000260228.3, ENST00000683306.1 |
| *Pan troglodytes* | ENSPTRT00000029292.6,ENSPTRT00000064657.3,ENSPTRT00000045270.2,ENSPTRT00000065610.3,ENSPTRT00000084033.1,ENSPTRT00000030049.4,ENSPTRT00000017231.5,ENSPTRT00000005563.4,ENSPTRT00000091821.1,ENSPTRT00000007863.3,ENSPTRT00000030041.4 |
| *Pan paniscus* | ENSPPAT00000038165.1,ENSPPAT00000063516.1,ENSPPAT00000065436.1,ENSPPAT00000032117.1, ENSPPAT00000059429.1, ENSPPAT00000002118.1, ENSPPAT00000044929.1, ENSPPAT00000008319.1, ENSPPAT00000045147.1 |
| *Gorilla gorilla* | ENSGGOT00000047574.1 ,ENSGGOT00000024652.2 ,ENSGGOT00000013709.3, ENSGGOT00000023433.2, ENSGGOT00000016458.3, ENSGGOT00000013270.3, ENSGGOT00000000517.3, ENSGGOT00000017106.3, ENSGGOT00000003330.3, ENSGGOT00000007947.3 |
| *Pongo abelii* | ENSPPYT00000016730.3,ENSPPYT00000055783.1, ENSPPYT00000017210.2, ENSPPYT00000023465.2, ENSPPYT00000017209.2, ENSPPYT00000010431.3, ENSPPYT00000003164.2, ENSPPYT00000017211.2, ENSPPYT00000004535.2, ENSPPYT00000017201.2 |
| *Nomascus leucogenys* | ENSNLET00000008418.2, ENSNLET00000034704.1, ENSNLET00000009824.2, ENSNLET00000011565.2, ENSNLET00000009808.2, ENSNLET00000011502.2, ENSNLET00000019531.2, ENSNLET00000009826.2, ENSNLET00000009150.3, ENSNLET00000009767.2 |
| *Macaca mulatta* | ENSMMUT00000024712.4, ENSMMUT00000005417.4, ENSMMUT00000042101.3, ENSMMUT00000011539.4, ENSMMUT00000066901.1, ENSMMUT00000004282.4, ENSMMUT00000002071.4, ENSMMUT00000021490.4, ENSMMUT00000015396.4, ENSMMUT00000072769.2 |
| *Macaca nemestrina* | ENSMNET00000006791.1, ENSMNET00000068311.1, ENSMNET00000040507.1, ENSMNET00000037668.1, ENSMNET00000058966.1, ENSMNET00000054703.1, ENSMNET00000035689.1, ENSMNET00000055162.1, ENSMNET00000022295.1, ENSMNET00000054841.1 |

##### 6.1.3 ‘Independent’ Dataset

Finally, a dataset labelled ‘Independent’ was utilised to reaffirm the phylogenetic results, using an independently created reference dataset. The ‘Independent’ dataset contains multiple translated samples from the genomes of different modern hominids. This dataset also includes the archaic protein sequences from previously published palaeoproteomic studies.

All translated protein sequences, except Neanderthals and the Denisovan individual were predicted from translating DNA data from VCF files and the alongside published annotation [54–59] or an annotation obtained from hg38 using liftover (https://genome-store.ucsc.edu/). For the prediction, beginning (“start_co”) and ending codons (“stop_co”) of each gene’s exon were extracted from the GTF annotation file. Then a consensus sequence of the assembly incorporating the variants has been made using samtools v1.9 [60] and bcftools v1.9 [61] with the command:

```
samtools faidx assembly.fasta ${<scaffold>}:${<start_co>}-${<stop_co>} | bcftools consensus --sample samplename -m mask.bed -H 1 file.vcf
```

At heterozygous positions, bcftools was randomly run either with the option -H1 or -H2 for random allele selection. The resulting open reading frame was translated with phython3, using standard genetic code. Low quality regions that are represented by the character “N” at DNA level were translated into masked amino acids represented by the character “X”.
The proteins were aligned using MAFFT v7.490 [62]:

```
mafft --maxiterate 1000 --globalpair protein.fasta > aligned_protein.fasta
```

A first round of less stringent trimming followed using TrimAl v1.2rev59 [63]:

```
trimal -in aligned_protein.fasta  -out trimmed_protein.fasta  -gt 0.9 -cons 60
```

For some proteins the stringency at this first step of trimming was lowered, leaving out the -cons option (minimum percentage of the positions in the original alignment to conserve after trimming) and lowering gt (the minimum amount of non-gap characters in a column): COL1A1 and COL17A1, -gt = 0.4; COL1A2 and COL2A1, gt = 0.8; ENAM, gt = 0.9. The resulting alignments were manually checked for possible misalignments, which were then masked representing the masked amino acid by the character “X”.

Alongside the ancient peptide sequences of *Paranthropus*, other published Hominoid ancient peptide sequences of *Homo antecessor*, *Homo erectus* [46], and *Gigantopithecus blacki* [45] were also analysed. All ancient sequences were added subsequently to the alignment. To prevent any misalignment because of the fragmented nature of the ancient sequences, a curated alignment of the ancient sequences to a human reference was added to the previously created reference alignments with the command:

```
mafft --maxiterate 1000 --add trimmed_protein.fasta ancient_protein_with_humanref.fasta > aln_prot.fasta
```

The human reference sequence that came with the ancient sequence was then removed from the resulting alignment.

Enamel proteins sequences of Neanderthals and the Denisovan individual were predicted as described above, using publicly available VCF files for the individuals "Altai" (http://cdna.eva.mpg.de/neandertal/altai/AltaiNeandertal/VCF/), "Vindija33.19" (http://ftp.eva.mpg.de/neandertal/Vindija/VCF/Vindija33.19/), and "Denisovan" (http://cdna.eva.mpg.de/neandertal/altai/Denisovan/).

#### 6.2. Protein sequence processing (‘Representative’ & ‘Diversity’ datasets)

Both datasets were processed in the same exact manner, using PaleoProPhyler’s [64] 3rd module (Extended Data Fig 5). Below you will find a small description of each step of this process.

##### 6.2.1 Alignment

The reference dataset was merged with the reconstructed proteins of the 4 *Paranthropus* samples. Each protein was then separated in its own multiple sequence alignment (MSA). An alignment was run on each of the MSAs utilising Mafft [62] as well as a 2-step alignment process:

- Modern samples were separated from the ancient samples and aligned by themselves using mafft’s:

```
mafft --ep 0 --op 0.5 --lop -0.5 --genafpair --maxiterate 20000 --thread {threads} --bl 80 --fmodel modern_samples.fa > modern_samples_aligned.fa
```

- The aligned modern sequences were then also aligned onto the ancient samples using maff’s:

```
mafft-einsi  --addlong ancient_samples.fa modern_samples_aligned.fa > final_alignment.fa
```

This 2-step process minimised the misalignments caused by the high missingness and presence of gaps in the ancient data. The individual protein MSAs were also manually inspected and corrected for any possible misalignments.

##### 6.2.2 Isobaric sites

The MSAs were automatically corrected for the isobaric amino acids of Isoleucine (I) and Leucine (L). The same process as described in Welker et al [46] was followed: positions in the alignment where either an I or an L was present in the recovered ancient proteins were marked. If *all* modern samples present in the dataset were bearing either an I or an L, the amino acids of the ancient samples were switched to match that. If some of the modern samples were bearing an I and some were bearing an L, all Is were switched to Ls for that particular site and for all sequences in the MSA. The final aligned and I/L corrected MSA for both reference datasets are available as Fasta files at https://zenodo.org/record/7801259:

##### 6.2.3 Concatenation and formatting

The aligned and I/L corrected protein datasets were concatenated and converted to the appropriate format for each phylogenetic software. If a sample in the alignment was missing a specific protein (e.g. female samples missing the AMELY protein), it had the entire length of the missing protein replaced by the missing symbol ‘?’ in the concatenation. Both the ‘Representative’ and ‘Diversity’ concatenated datasets are also available in all the various formats at: https://zenodo.org/record/7801259

#### 6.3 Phylogenetic inference ('Representative' & 'Diversity' datasets)

The aligned and I/L corrected sequences were used to generate multiple phylogenetic trees using the different datasets, genes, software and sample compositions. With the exception of IQtree Beast2 and StarBeast3, all other trees were generated as part of the PaleoProPhyler workflow. All trees presented in the supplementary were plotted using Figtree (http://tree.bio.ed.ac.uk/software/figtree/). The commands for each tree generated through this process are presented below.

##### 6.3.1 MrBayes, PhyML, IQ tree

- Command for MrBayes [65]:

```
mb MrBatch.txt > log.txt;
```

Where 'MrBatch.txt' is a txt file that contains the input below:

```
set autoclose=yes
execute CONCATINATED_o.nex
charset AMELY = 1-206;
charset ENAM = 207-1348;
charset COL17A1 = 1349-2861;
charset AMBN = 2862-3308;
charset AMTN = 3309-3517;
charset AHSG = 3518-3885;
charset ALB = 3886-4494;
charset ODAM = 4495-4773;
charset COL1A1 = 4774-6278;
charset AMELX = 6279-6483;
charset MMP20 = 6484-6966;
partition BY_PROTEIN = 11: AMELY, ENAM, COL17A1, AMBN, AMTN, AHSG, ALB, ODAM, COL1A1, AMELX, MMP20;
set partition=BY_PROTEIN;
prset aamodelpr = mixed;
mcmc nchains = 4 nruns=4 ngen = 5000000 samplefreq=100 printfreq=100 diagnfreq=1000;
sumt relburnin = yes burninfrac = 0.25;
sump;
Quit;
```

- Command for PhyML [66]:

```
phyml-mpi -i CONCATINATED_aln_e.phy -d aa -b 100 -m JTT -a e -s BEST -v e -o tlr -f m --rand_start --n_rand_starts 4
--r_seed $RAND --print_site_lnl --print_trace --no_memory_check
```

- Command for IQ tree [67]:

```
path_to_iqtree -s CONCATINATED_aln_e.phy -st AA -m Blosum62+F -bb 1000 -alrt 1000
```

##### 6.3.2 BEAST2, StarBeast3

BEAST2 Analysis [68]:

In addition to the PaleoProPhyler workflow, a Beast2 tip dated analysis was also performed. A set of 2 individuals per subspecies was randomly selected from the concatenated 'Diversity' dataset. Following [46], the gamma site model category was set to 4 and JJT was chosen as the substitution model through BEAUti v2.1. The ages of each ancient individual were included in this analysis and are as follows :

- Altai Neanderthal - 112 kya [53]
- Chagyrskaya Neanderthal - 80 kya [69]
- Altai Denisovan - 82 kya [70,71].
- All 4 *Paranthropus* individuals - 1.8 mya (this study)

The XML file generated through BEAUti and used in this analysis is available at https://zenodo.org/record/7801259

StarBeast 3 [72]

StarBEASTv3 was also employed to generate a multispecies coalescent phylogenetic tree. BEAUti v2.1 was used to generate the input XML file for the analysis and BEAST2, with the Starbeast3 extension, was used to run it. The phylogenetic reconstruction was carried out, using the 'Diversity' dataset. All samples of the dataset were assigned to a taxon. Humans, Neanderthals and Denisovans were assigned to different taxons. For the samples belonging to the species *Pan paniscus*, *Pan troglodytes* and *Gorilla gorilla*, 3 taxons were considered, disregarding subspecies. For the 3 species of the genus *Pongo,* 3 taxons were also considered. All four *Paranthropus* samples were assigned to the same taxon. We applied the Blosum62 amino acid substitution model and strict molecular clock. The MCMC chains were run for more than 40.000.000 states and sampled every 50.000 states. The MCMC runs were inspected for ESS values above the recommended value of 100 using Tracer (http://tree.bio.ed.ac.uk/software/tracer/) and the run was terminated soon after they were sufficiently above that number. The XML file and the txt file linking each sample to a taxon is available at https://zenodo.org/record/7801259.

#### 6.4 ‘Independent’ dataset phylogenetic analysis

The protein alignments of the independently generated reference dataset (see section 6.1.3 ‘Independent’ Dataset) were downsampled to great apes with a maximum of three individuals per (sub-)species and one individual for the outgroup (*Papio papio*). The downsampled alignments were then concatenated using catfasta2phyml.pl (https://github.com/nylander/catfasta2phyml) with the command:

```
catfasta2phyml.pl --fasta --concatenate --verbose *.fasta
```

All isobaric sites in the ancient sequences (isobaric amino acids of Isoleucine (I) and Leucine (L)) were changed to Leucine. If any reference protein sequence had an Isoleucine at the homologous position, it was changed to Leucine, as well. The resulting concatenated alignment, excluding lines with ancient proteins, underwent one more round of trimming first on the samples (rows) using TrimAl [63]:

```
trimal -in concatenated_alignment.fasta -out concatenated_alignment_trimmed.fasta -resoverlap 0.50 -seqoverlap 75
```

Each protein in the concatenated alignment forms a partition. The evolutionary model for each protein, i.e. partition was estimated using the ModelFinder [73] option in IQtree v1.6.12:

```
iqtree  -s alignment_partition.fasta -nt {{threads}  -m MF  -msub nuclear
```

The resulting models were stored in a nexus file. This file then informed the creation of a Maximum Likelihood phylogeny in IQtree:

```
iqtree -s alignment.fasta  -alrt 5000  -bb 5000  -nstop 500  -nm 10000  -wsplits  -nt {threads}
```

With the same alignment and definition of partitions, a bayesian phylogeny was created using MrBayes v3.2.7a. In the concatenated alignment, each gene formed a partition for which various parameters were estimated (lset app=(1,2,3,4,5,6,7,8,9,10) rates=invgamma nucmodel=Protein; unlink statefreq=(all) revmat=(all) shape=(all) pinvar=(all); prset applyto=(all) ratepr=variable aamodel=mixed). The MCMC algorithm ran through 3 000 000 iterations in 2 runs on 4 chains. The temperature parameter was set to 0.2, the starttree was random. The convergence of the algorithm was assessed using Tracer v.1.7.2 after discarding 25% of the iterations as burn-in.

Both Maximum Likelihood and Bayesian analysis were additionally performed as above on an alignment that only retains columns, if an amino acid from at least one ancient sample is present. For these shortened alignments no partitions were defined and all regarding parameters left at default.

All phylogenies were visualised in figtree v1.4.4.

#### 6.5 Genetic Variation Metrics

A test was designed in order to assess the probability of sampling 4 individuals from the population of a species and identifying variation on the protein level. Due to the huge availability of genetic data, modern humans (*Homo sapiens*) were chosen as our species of comparison. Our test was created in the form of a snakemake workflow [74] and is available online at Github: (https://github.com/johnpatramanis/Code_for_Genetic_Diversity_Sampling). The github repository includes a small tutorial on how to reproduce the results, as well as a Conda environment (https://docs.conda.io/en/latest/) with all the necessary prerequisite software. The full workflow is described below.

Gnomad's [75] merging of the 1000 Genomes Project [76] and the Simons Genome Diversity Project [57] callset (https://gnomad.broadinstitute.org/downloads#v3-hgdp-1kg) was used as an (imperfect) sample of *Homo sapiens*'s genetic variability. Since all of the proteins of interest were located on one of the following chromosomes: **3,4,10,11,17,X,Y**, only the relevant chromosome files were downloaded from Gnomad. These chromosome files were then merged into a single VCF [77] file. The VCF file constituted the input to our workflow, however any alternative VCF file can also be utilised for the exact same analysis, as long as it contains data mapped onto the GRCh38 human reference genome [78] .

The proteins and exact amino acid positions that were recovered for all 4 *Paranthropus* individuals were first identified. A total of **424** shared amino acids across **6** proteins were detected. The exact positions of these shared amino acids were outputted and are available in the 'Protein_Coverage.txt' file. Note that this file can also be reproduced by running the 'Get_Protein_Coverage.py' python3 [79] script but requires the *Paranthropus* fasta alignments to be placed in the same folder. The *Paranthropus* fasta alignments are also available in the Github repository for convenience.

Variants that were within the genes that code the 6 proteins of interest were isolated from the input VCF file using bcftools [60]. The workflow requires that the genetic coordinates of those genes are provided through a file named 'Gene_locs_pure.txt'. This file should have one genome coordinate per line (e.g. **chr4 70628744 70646824**), each corresponding to one gene of interest.

The isolated variants were then used as input for Ensembl's Variant Effect Predictor (VEP) [80]. Version 108 of VEP, and the GRCh38 cache (https://ftp.ensembl.org/pub/release-109/variation/vep/homo_sapiens_vep_109_GRCh38.tar.gz) were chosen for this analysis. Variants that had a missense effect on the amino acid sequence were then identified from the result of VEP and outputted as a list of genetic coordinates. Only variants that had an effect on the regions of the protein that were recovered for all of our 4 *Paranthropus* individuals ('Protein_Coverage.txt') were kept. The outputted genetic coordinates were then used by bcftools to isolate these variants from the VCF file. The genotypes of all individuals for these variants were outputted in a ‘genotypes’ .txt file, with one variant per line and columns equal to the number of individuals of the original VCF file.

This ‘genotypes’ file was then loaded into python3 where all the metrics were calculated. Initially 3 metrics to assess the population genetic diversity of our dataset were obtained: a) Expected heterozygosity (2*p*q), b) Observed heterozygosity (proportion of heterozygous sites) and c) Watterson’s estimator [81] for the population genetic parameter theta. For the final step 4 individuals from this dataset were randomly sampled and their genotypes (8 alleles) were checked for all of the potential variant positions. Afterwards, 4 measurements at the amino acid level were recorded in order to compare to what was observed in Paranthropus: 1) If any of the sites have at least one alternative amino acid allele. 2) How many of the sites have at least one alternative amino acid allele. 3) If any of the sites have at least 2 copies of the alternative amino acid allele. 4) If any of the sites have at least one individual homozygous for the alternative amino acid allele. This process was carried out on 4-individual samples, and repeated 1000 times, without replacement of the sampled present-day human individuals.

### 7. Geometric morphometric analysis

The enamel-dentine junction (EDJ) of the $P_4$ SK 830 and of the $M^3$ SK 835 [34] were analysed and compared with those of *Homo*, *Australopithecus* and *Paranthropus* (including TM 1517, the holotype of *Paranthropus robustus*; Table S4).

Both SK 830 and SK 835 were scanned by X-ray microtomography with the General Electric V|Tome|x s industrial microCT system at the PLACAMAT platform (University of Bordeaux, France), and with the EasyTom XL Duo instrument at the PLATINA platform of the IC2MP (University of Poitiers, France) [34]. The scans were done according to the following parameters: 70-110 kV voltage; 340-350 mA current; 0.1 mm Cu and 1.2 mm Al filters. The final volumes were reconstructed with an isotropic voxel size of 27.5 µm and 25 µm for SK 830 and SK 835, respectively.

The comparative specimens were scanned using either X-ray or neutron microtomography and the final volumes were reconstructed with an isotropic voxel size of 10-30 µm for isolated teeth and 40-60 µm for jaw fragments [82]. For all specimens, image stacks were imported into Avizo v.8.0 (FEI Visualization Sciences Group), and the images were segmented using semiautomatic procedures and an adaptation of the half-maximum height method [83–85]**.** All the EDJ surfaces were generated using the 'constrained smoothing' option.

We used a diffeomorphic surface matching (DSM) approach to analyse the EDJ conformation. This landmark-free, mesh-based approach relies on the construction of average surface models, and the difference between surfaces is interpreted as the amount of deformation needed to align them by using diffeomorphic shape matching [86,87]. The metric of currents used in DSM analyses takes all data points into account and does not assume a point-to-point correspondence between samples which allows direct comparison of surfaces that have different numbers of sample points [88]. Moreover, this metric takes into account the local orientation of a surface (i.e., the normals) to strengthen the measure of shape dissimilarities (this metric does not only measure how distant two surfaces are, but also how their respective local orientations differ) [88]. The deformations between surfaces are mathematically modelled as smooth and invertible functions (i.e., diffeomorphisms). From a set of surfaces, an atlas of surfaces is created. The method estimates an average object configuration or mean shape from a collection of object sets (here the EDJ surfaces) and computes the deformations from the mean shape to each specimen. In addition, a set of initial control points located near its most variable parts, and a set of momenta parameterizing the deformations of the mean shape to each individual are estimated [82,88–93]. When comparing a group of closely related specimens (as it is the case here), it is assumed that the inaccuracy (noise) introduced by comparing whole surfaces is small compared to the information that can be extracted (signal). Compared with landmarks, correspondence between surfaces is not strictly homologous. The assumption about signal to noise ratio has been evaluated in previous studies, showing that DSM can distinguish the EDJ of various hominin groups, even better than with landmark-based analyses, suggesting that DSM can offer reliable accuracy and precision, of a similar order than landmark-based analyses [82,90,92]. For each dental position, the EDJ surfaces decimated to 50000 polygons were manually oriented, then superimposed using the rigid and uniform scale option (corresponding to a shape alignment, removing size) of the 'Align Surfaces' module in Avizo. This was done by minimizing the root mean square distance between the points of each specimen to corresponding points on the reference surface using an iterative closest point algorithm. We used the Deformetrica v. 4.3 software (https://www.deformetrica.org) [87,88] to generate a global mean shape (GMS) with a set of diffeomorphisms relating the GMS to each individual and the output (control points and deformation momenta) used to perform the statistical analyses to explore the EDJ shape variation and to classify the data. The output data were imported in R with the package RToolsForDeformetrica v.0.1 [94]. Using the packages ade4 v.1.7-6 [95] and Morpho v.2.8 [96] for R v.4.0.4

[97], we started by computing principal component analyses (PCA). We then performed cross-validated canonical variates analysis (CVA) based first on three groups assigned with equal prior probabilities (*Early-Middle Pleistocene Homo*, *Australopithecus* and *Paranthropus*) using the R package Morpho v.2.8 [98]. Since CVA computation requires the number of variables to be much smaller than the number of specimens, we computed the CVA based on a subset of the first PC scores [87–90] showing the highest degree of correct classification (screening the correct classification results and selecting the minimum number of PC scores enabling to reach the optimum of correct classification) [99]. This choice of the PC scores subset is a compromise between including a sufficient proportion of overall shape variation and limiting the number of variables to avoid unrealistic and unstable levels of discrimination [100].

Correct classification for each group and overall classification resulting from the cross-validated CVA are reported in Table S5. The specimens SK 830 and SK 835 were projected *a posteriori* into the CVA morphospace. Typical probabilities confirmed that SK 830 and SK 835 very likely belonged to *Paranthropus,* and not to *Australopithecus* or *Homo* (Table S6). We ran DSM analyses restricted to the *Paranthropus* subsample and PCAs. We computed cross-validated CVAs based on 8 PCs for both $M^3$s and $P_4$s using two groups (distinguishing the specimens from Swartkrans and Kromdraai [SK] on the one hand and those from Drimolen [D] on the other). Only for one specimen from Swartkrans, SKW 11, the morphology looked more similar to that of the Drimolen specimens. If considered a priori as part of the group SK, it was misclassified as belonging to group D and decreased substantially the overall classification rate (75%). Further inspection of the EDJ morphology revealed closer similarities between SKW 11 and the Drimolen specimen than with those from Swartkrans and Kromdraai (Figure S1). Some specimens from Drimolen (DNH 39, DNH 62, DNH 67, and DNH 70) and Swartkrans (SK 15) recently attributed to *Paranthropus* show a more simple EDJ shape (without accessory crests or cusps, and without Carabelli trait or protostylid) than TM 1517 and the major part of the *P. robustus* hypodigm [82], comparable to the morphology of SKW 11. Consequently, the specimen SKW 11 was considered as representing the same group as the Drimolen specimens and included in group D. The specimens SK 830 and SK 835 were then projected a posteriori and their typicality probabilities were computed. Results of these analyses are reported in Tables S7 and S8.

**Table S4: List of the comparative microtomographic (microCT) data of dental remains attributed to *Australopithecus* (AUS), Early and Middle Pleistocene *Homo* (HOM), and *Paranthropus* (PAR)**

| specimen | tooth | site | taxonomic attribution | Chronology | ref. of microCT data |
|---|---|---|---|---|---|
| StW 2 | $M^3$ | Sterkfontein (South Africa) | AUS | ~2.8-2.2 Ma | [82] |
| StW 43 | $M^3$ | Sterkfontein (South Africa) | AUS | ~2.8-2.2 Ma | [82] |
| StW 92 | $M^3$ | Sterkfontein (South Africa) | AUS | ~2.8-2.2 Ma | [82] |
| StW 128 | $M^3$ | Sterkfontein (South Africa) | AUS | ~2.8-2.2 Ma | [82] |
| StW 189 | $M^3$ | Sterkfontein (South Africa) | AUS | ~2.8-2.2 Ma | [82] |
| StW 252 L | $M^3$ | Sterkfontein (South Africa) | AUS | ~2.8-2.2 Ma | [82] |
| StW 252 R | $M^3$ | Sterkfontein (South Africa) | AUS | ~2.8-2.2 Ma | [82] |
| StW 277 | $M^3$ | Sterkfontein (South Africa) | AUS | ~2.8-2.2 Ma | [82] |
| StW 483 | $M^3$ | Sterkfontein (South Africa) | AUS | ~2.8-2.2 Ma | [82] |
| StW 498 | $M^3$ | Sterkfontein (South Africa) | AUS | ~2.8-2.2 Ma | [82] |
| StW 524 | $M^3$ | Sterkfontein (South Africa) | AUS | ~2.8-2.2 Ma | [82] |
| CA 772 | $M^3$ | China | HOM | Middle Pleistocene | ESRF heritage database for palaeontology, evolutionary biology and archaeology (2021). http://paleo.esrf.eu. |
| M3550 | $M^3$ | Zhoukoudian (China) | HOM | ~0.8-0.7 Ma | [82,90,101,102]) |
| NG0802.1 | $M^3$ | Ngebung (Indonesia) | HOM | Middle Pleistocene | [102] |
| Sangiran 4 | $M^3$ | Sangiran (Indonesia) | HOM | Early Pleistocene | [82,90,101,102]) |
| Sangiran 7-3d | $M^3$ | Sangiran (Indonesia) | HOM | Early Pleistocene | [82] |
| Sangiran 7-17 | $M^3$ | Sangiran (Indonesia) | HOM | Early Pleistocene | [82] |
| DNH 3 | $M^3$ | Drimolen (South Africa) | PAR | ~2.0 Ma | this study |
| DNH 22 | $M^3$ | Drimolen (South Africa) | PAR | ~2.0 Ma | this study |
| DNH 54 | $M^3$ | Drimolen (South Africa) | PAR | ~2.0 Ma | this study |
| SK 31 | $M^3$ | Swartkrans (South Africa) | PAR | ~2.2-1.8 Ma | [82] |
| SK 49 | $M^3$ | Swartkrans (South Africa) | PAR | ~2.2-1.8 Ma | [82] |
| SK 52 | $M^3$ | Swartkrans (South Africa) | PAR | ~2.2-1.8 Ma | [82] |
| SK 105 | $M^3$ | Swartkrans (South Africa) | PAR | ~2.2-1.8 Ma | [82] |
| SK 831a | $M^3$ | Swartkrans (South Africa) | PAR | ~2.2-1.8 Ma | [82] |
| SK 836 | $M^3$ | Swartkrans (South Africa) | PAR | ~2.2-1.8 Ma | [82] |
| SKW 11 | $M^3$ | Swartkrans (South Africa) | PAR | ~2.2-1.8 Ma | [82] |
| TM 1517 | $M^3$ | Kromdraai (South Africa) | PAR | initial Early Pleistocene | [82] |
| MLD 2 L | $P_4$ | Makapansgat (South Africa) | AUS | Late Pliocene | [82] |
| MLD 2 R | $P_4$ | Makapansgat (South Africa) | AUS | Late Pliocene | [82] |
| Sts 52 | $P_4$ | Sterkfontein (South Africa) | AUS | ~2.8-2.2 Ma | [82] |
| StW 14 | $P_4$ | Sterkfontein (South Africa) | AUS | ~2.8-2.2 Ma | [82] |
| StW 56 | $P_4$ | Sterkfontein (South Africa) | AUS | ~2.8-2.2 Ma | [82] |
| StW 104 | $P_4$ | Sterkfontein (South Africa) | AUS | ~2.8-2.2 Ma | [82] |
| StW 131 | $P_4$ | Sterkfontein (South Africa) | AUS | ~2.8-2.2 Ma | [82] |
| StW 140 | $P_4$ | Sterkfontein (South Africa) | AUS | ~2.8-2.2 Ma | [82] |
| StW 142 | $P_4$ | Sterkfontein (South Africa) | AUS | ~2.8-2.2 Ma | [82] |

| | | | | | |
|---|---|---|---|---|---|
| StW 194 | $P_4$ | Sterkfontein (South Africa) | AUS | ~2.8-2.2 Ma | [82] |
| StW 232 | $P_4$ | Sterkfontein (South Africa) | AUS | ~2.8-2.2 Ma | [82] |
| StW 327 | $P_4$ | Sterkfontein (South Africa) | AUS | ~2.8-2.2 Ma | [82] |
| StW 404 | $P_4$ | Sterkfontein (South Africa) | AUS | ~2.8-2.2 Ma | [82] |
| StW 413 | $P_4$ | Sterkfontein (South Africa) | AUS | ~2.8-2.2 Ma | [82] |
| StW 498 L | $P_4$ | Sterkfontein (South Africa) | AUS | ~2.8-2.2 Ma | [82] |
| StW 498 R | $P_4$ | Sterkfontein (South Africa) | AUS | ~2.8-2.2 Ma | [82] |
| KNM-ER 992B L | $P_4$ | Koobi Fora (Kenya) | HOM | ~1.5 Ma | [82] |
| KNM-ER 992B R | $P_4$ | Koobi Fora (Kenya) | HOM | ~1.5 Ma | [82] |
| M3887 | $P_4$ | Zhoukoudian (China) | HOM | ~0.8-0.7 Ma | [82,90,101,102]) |
| PA81 | $P_4$ | Changyang (China) | HOM | ~0.2 Ma | [93,103,104] |
| PA525 | $P_4$ | Xichuan (China) | HOM | Middle Pleistocene | [93,103,104] |
| PA528 | $P_4$ | Xichuan (China) | HOM | Middle Pleistocene | [93] |
| Sangiran 1b | $P_4$ | Sangiran (Indonesia) | HOM | Early Pleistocene | [101,105] |
| Sangiran 7-26 | $P_4$ | Sangiran (Indonesia) | HOM | Early Pleistocene | [82] |
| Tighenif 2 | $P_4$ | Tighenif (Algeria) | HOM | ~1.0 Ma | [106] |
| DNH 27a | $P_4$ | Drimolen (South Africa) | PAR | ~2.0 Ma | this study |
| DNH 51 | $P_4$ | Drimolen (South Africa) | PAR | ~2.0 Ma | this study |
| DNH 68 | $P_4$ | Drimolen (South Africa) | PAR | ~2.0 Ma | this study |
| SK 23 | $P_4$ | Swartkrans (South Africa) | PAR | ~2.2-1.8 Ma | [82] |
| SK 25 L | $P_4$ | Swartkrans (South Africa) | PAR | ~2.2-1.8 Ma | [82] |
| SK 25 R | $P_4$ | Swartkrans (South Africa) | PAR | ~2.2-1.8 Ma | [82] |
| SK 34 L | $P_4$ | Swartkrans (South Africa) | PAR | ~2.2-1.8 Ma | [82] |
| SK 55 | $P_4$ | Swartkrans (South Africa) | PAR | ~2.2-1.8 Ma | [82] |
| SK 828 | $P_4$ | Swartkrans (South Africa) | PAR | ~2.2-1.8 Ma | [82] |
| SK 1587a | $P_4$ | Swartkrans (South Africa) | PAR | ~2.2-1.8 Ma | [82] |
| SK 1588 | $P_4$ | Swartkrans (South Africa) | PAR | ~2.2-1.8 Ma | [82] |
| SKW 5 | $P_4$ | Swartkrans (South Africa) | PAR | ~2.2-1.8 Ma | [82] |
| TM 1517 | $P_4$ | Kromdraai (South Africa) | PAR | initial Early Pleistocene | [82] |

### SUPPLEMENTARY RESULTS

#### 8. Biomolecular preservation

##### 8.1. Intra-crystalline protein degradation analyses

Dental enamel subsamples from the four *Paranthropus* specimens studied using palaeoproteomics were analysed using reverse phase high-performance liquid chromatography (RP-HPLC) to measure amino acid concentration, composition and racemisation. This approach validated the endogenous origin of the enamel proteins we identified, by comparing the amino acid composition and racemisation profiles of the four *Paranthropus* samples with those from hominin samples previously analysed.

In a closed system, the ratios of the two amino acid (AA) enantiomers (D and L) from the free (FAA) and the total hydrolysable (THAA) fractions are expected to be highly correlated, enabling the identification of compromised samples[107,108]. In such a system, as long as no (or negligible) mineral diagenesis has occurred, the extent of intra-crystalline protein degradation is solely time and temperature dependent, with no leaching of degradation products or introgression of exogenous contaminants[109]. The D/L values of all the amino acids will increase with time, but each amino acid racemises at a different rate, therefore providing different resolution over different timescales. To date, the only other Early Pleistocene hominin dental enamel sample analysed to measure amino acid racemisation originates from an isolated *H. erectus* dental specimen (D4163) from the ~1.8 Ma archaeological site of Dmanisi (Georgia)[46], here used for comparison.

###### 8.1.1. Amino acid composition

The amino acid composition profiles of the four analysed dental enamel samples from the Swartkrans cave are consistent with each other and are coherent with the expected composition of other hominin enamel samples dated approximately to the early Pleistocene [40,107] (Figure S2 A). The amino acids used to calculate the summed concentrations deriving from the THAA fraction are: Asx, Glx, (aspartic acid/asparagine and glutamic acid/glutamine respectively), Ser, L-Thr, L-His, Gly, Arg, Ala, Tyr, Val, Phe, Leu and Ile. Compared to collagen-based biominerals, such as bone and dentine, in general the proportion of Gly in the THAA fraction of the enamel from Swartkrans is slightly lower (about 20-35 % compared to about > 40 %) and the proportion of Glx and Leu are higher. The proportion of Gly in the Swartkrans samples is greater than that from the Dmanisi *H. erectus* enamel, but it is still lower than the expected levels found in the dentine from the same specimen [46]. Additionally, the proportion of Glx and Leu are also more similar to the composition of enamel than for dentine. This indicates the amino acids analysed for all four samples are most likely derived from enamel proteins, as opposed to dentine collagen or other exogenous sources.

The total summed concentration from the THAA portion for samples SK 830, SK 835 and SK 14132 are fairly consistent (1350-1530 pmol $mg^{-1}$) and are slightly lower then for the *H. erectus* dental enamel from Dmanisi (1670 pmol $mg^{-1}$), as expected from specimens of comparable age, but originating from lower latitudes. For sample SK 850 this value is lower (~1030 pmol $mg^{-1}$) than those for the other teeth from Swartkrans, indicating lower levels of total amino acid concentrations in this tooth (Figure S2 B). For specimens where both the FAA and THAA portions were analysed, the percentage of FAA amino acids were generally high (~50-85 %), consistent with closed system amino acids of Early Pleistocene age.

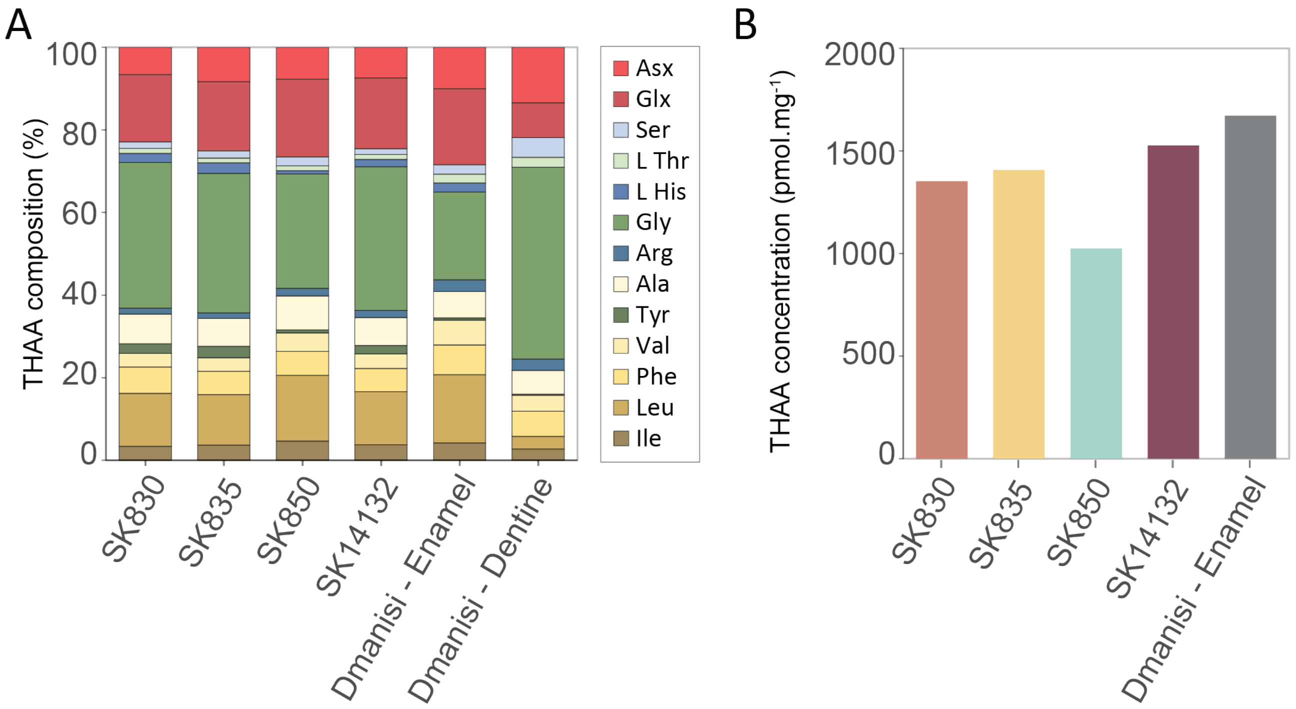


**Figure S2: Amino acid composition and concentration of the dental enamel.** A - Summed amino acid concentration from the THAA fraction [46]. B - THAA composition of the four *Paranthropus* teeth from Swartkrans, and the enamel and dentine fractions from a *H. erectus* tooth from Dmanisi (D4163) for comparison [41].

##### 8.1.2. Amino acid racemisation

In a closed system, all of the products of protein degradation should be retained, so the racemisation of the FAA and THAA fractions should be highly correlated over geological time. More labile amino acids, such as those that are unbound (FAA) are more likely to leach out of an open system, so open system behaviour tends to result in data points falling away from the general closed system trend[39]. Whilst taxonomic effects influence the rates of racemisation, it is likely that the relative relationship between the racemisation of FAA and THAA is similar enough between taxa to identify whether significant leaching of the endogenous protein, and/or introgression of exogenous proteinaceous material has occurred. The D/L enantiomer values of aspartic acid/asparagine, glutamic acid/glutamine, phenylalanine and alanine (D/L Asx, Glx, Phe and Ala, respectively) were measured.

The FAA and THAA racemisation values from the *Paranthropus* samples plot very close to the trajectory defined by previously analysed enamel samples belonging to multiple taxa from various geographic regions and chronologies (Figure S3). These results show that the *Paranthropus* enamel we analysed exhibits a closed system behaviour, strongly supporting the endogenous origin of the intra-crystalline enamel amino acids we analysed. While the Phe D/L values plot within the general trend defined by the reference samples (Figure S3A), the Glx D/L values plot slightly further away from it (Figure S3B); this minor discrepancy is likely due to the greater challenge in accurate measurement of Glx FAA D/L [110], due to its spontaneous lactam formation. Furthermore, the Swartkrans racemisation values we report would most likely fall within the general variation if a richer reference database, including more measures from Early Pleistocene specimens, was available. It was not possible to retrieve data for both the Glx and Phe FAA fractions from tooth SK 850 because of its low sample mass (Supplementary methods 4).

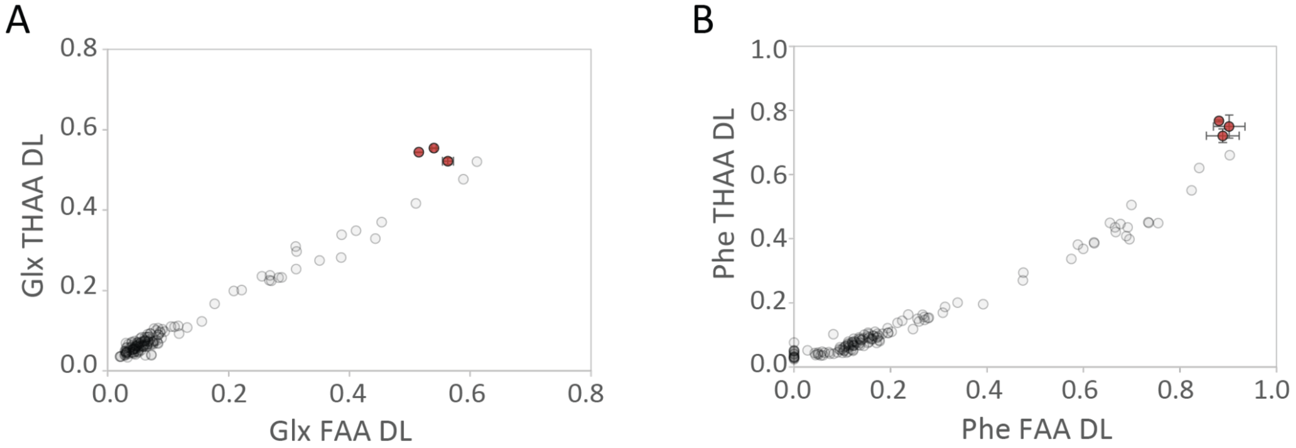


**Figure S3: Glx and Phe intra-crystalline FAA and THAA racemisation**. Comparison of the *Paranthropus* enamel intra-crystalline FAA vs THAA racemisation for two different amino acids, Glx (A) and Phe (B), to a larger database of Pleistocene enamel D/L values from a variety of different taxa (e.g. Elephantidae, Bison, Equids) and geographical regions. Note: only three of the four Swartkrans samples are plotted, due to the lack of data for the FAA fraction from enamel specimen SK 850.

The extent of racemisation in the four teeth from Swartkrans is relatively consistent for all of the amino acids studied, for both the FAA and THAA fractions, indicating they are similar in age (Figure S4, Figure S5 and Extended Data Fig 3). The extent of racemisation is close to equilibrium for Asx, Ala and Phe, consistent with the antiquity of the samples. Additionally, the extent of racemisation in the dental enamel specimens from the Swartkrans cave is greater than the *H. erectus* enamel specimen from Dmanisi. The rate of enamel intra-crystalline protein degradation is controlled by the integrated temperature history of the sample, which is different between Dmanisi (current Mixed Air Temperature (MAT) = 9 °C) and Swartkrans (current MAT = 16 °C), with less seasonality in this region of South Africa and therefore overall warmer temperatures. Although this parameter will have changed over Pleistocene timescales, the greater level of racemisation observed in the Swartkrans material is consistent with an Early Pleistocene age for this region.

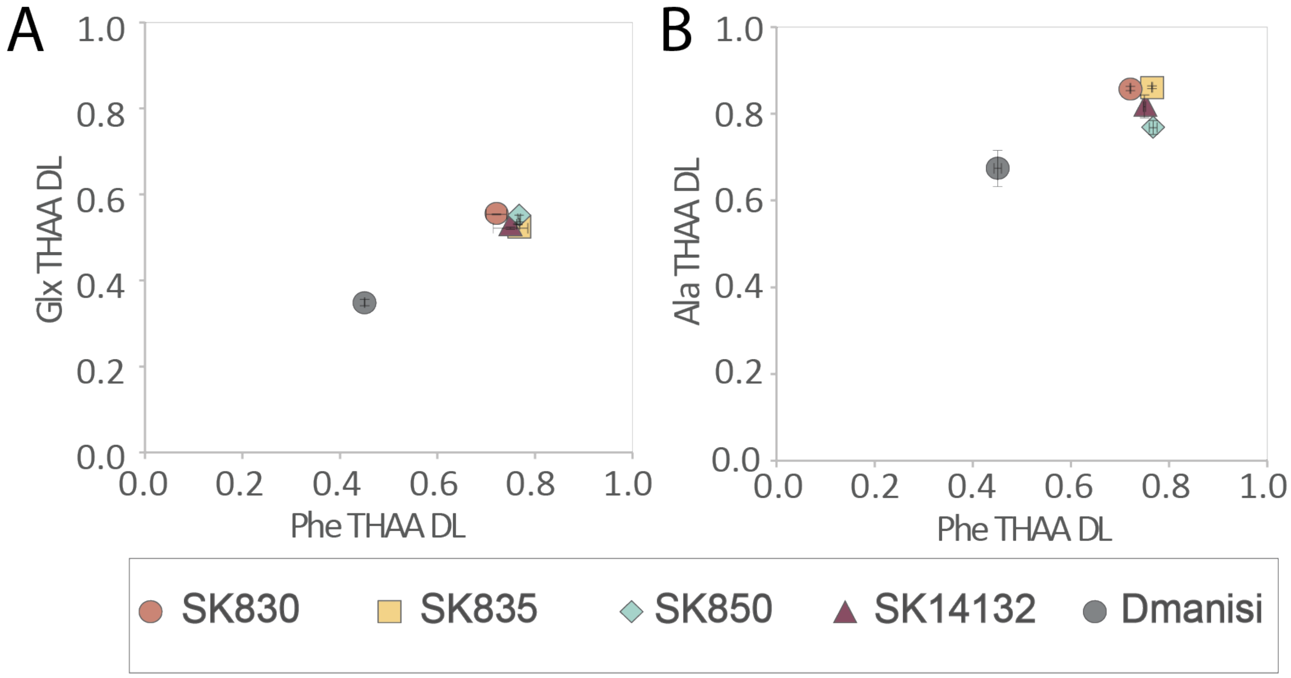

**Figure S4: Comparison of intra-crystalline THAA racemisation.** Comparison of the intra-crystalline THAA racemisation vs THAA racemisation covariance of Phe to Glx (A) and Ala (B), between the *Paranthropus* and the *H. erectus* enamel from Dmanisi. Error bars represent 1σ about the mean.

In conclusion, the amino acids analysed in the four *Paranthropus* teeth from the Swartkrans cave are consistent with being derived from enamel proteins preserved in a biomineral environment that exhibit closed system behaviour, with no identification of leaching or contamination. The extent of racemisation in the amino acids for the four teeth are comparable and consistent with the expectations of samples dating to ~2 Ma from this region. The extent of racemisation in the dental enamel from the Swartkrans cave is greater than in the enamel from Dmanisi, indicating a greater extent of protein degradation than the previously oldest hominin palaeoproteomic sample.

#### 8.2. Analysis of faunal remains

The analysis of the four faunal remains led to the successful recovery of endogenous peptide sequences. It was possible to recover on average 2000 PSM per sample analysed (Figure S5A), corresponding to an average of 340 amino acid positions covered using mass spectrometry (Figure S5B). Moreover, the identified peptide sequences showed evidence of age degradation with an average length of 9.8 amino acids and high modification rate. On average, more than 95% of the detected asparagine or glutamine were deamidated (Figure S5C). Additionally, close to a third of the detected arginine were converted to ornithine (Figure S5D).

All four samples showed consistent proteomics coverage. It was thus possible to confirm the possibility of recovering biomolecular material from samples originating from the same site and era as the *Paranthropus* samples.

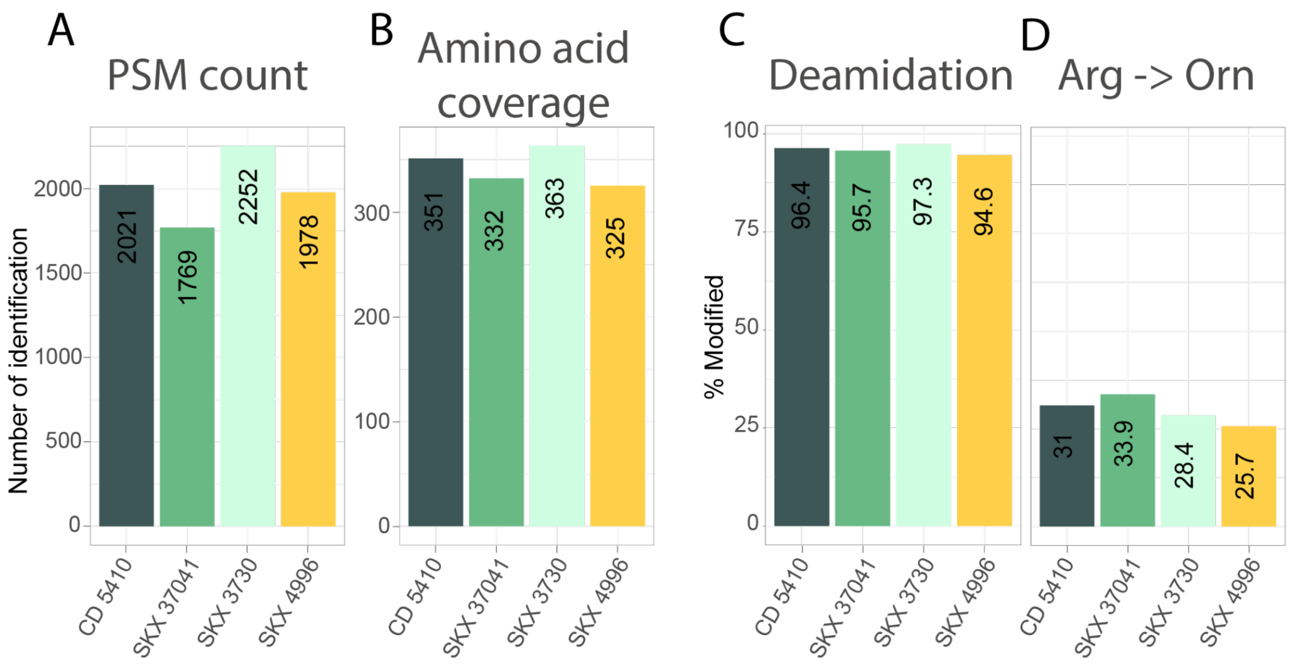


**Figure S5: Analysis of faunal remains.** A-B- Barplot showing the number of identifications in the four faunal specimens studies with A- the PSM count and B- the total amino acid coverage. C-D Barplots showing the percentage of modification per specimen studied with C- asparagine and glutamine deamidation and D- conversion or arginine to ornithine. The extent of modification (C and D) was calculated based on PSM counts.

### 9. Palaeoproteomics results

#### 9.1. Protein sequence database construction

Different databases were considered for reaching optimal depth in the sequence reconstruction. The choice of the database is crucial in palaeoproteomics since sequences absent from the database will not be identified [111,112]. Thus, the database should cover all sequences present in the samples. On the other hand, a large database will considerably increase the search space by providing more matching possibilities to the algorithm. This is especially critical while performing unspecific searches where the increase of the search space with the addition of sequences in the database is far greater than in a specific search [113]. Therefore, a balance had to be found between having a complete and large database and having a reduced one that is specific to the sample.

Three different databases were tested on three of the *Paranthropus* samples: SK 830, SK 850 and SK 14132 (Figure S6). Databases were constructed only with sequences previously identified in enamel samples, including the following genes: AHSG, ALB, AMBN, AMELX, AMELY, AMTN, COL17A1, COL1A1, COL1A2, COL2A1, ENAM, MMP20 and ODAM.

A first database was constructed with Uniprot sequences from Human, chimpanzee, gorilla and orangutan, supplemented with manually collected sequences from *Gigantopithecus blacki*, *Homo antecessor*, *Denisovan* and *Neanderthal*. This constitutes the "Core Database" and contains **622** protein sequences. A second broader database ("All Monkeys Database") was built adding seven more primate species enamelome from Uniprot and NCBI to the "Core Database" (Table S9) and regrouped **1090** protein sequences. This represents the main evolutionary branches of non-human primates. The third database ("Ape Genome Database") was built by adding enamel specific sequences for more than 250 species coming from a large genome collection with a total of **3957** protein sequences [59].

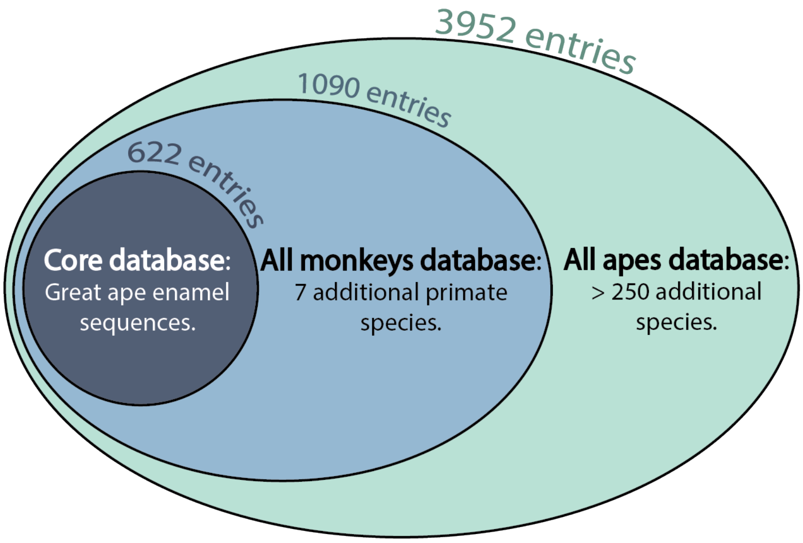


**Figure S6: Database selection**. Schematic representation of the construction of the three different databases tested: the core database, the "All Monkeys" database and the "All apes" database.

**Table S9: Construction of the database.** List of species included in the "All Monkeys" database.

| **Group** | **Species** | **Common name** |
|---|---|---|
| Human | *Homo sapiens* | Human |
| Extinct humans | *Homo antecessor* | *Homo antecessor* |

| | | |
|---|---|---|
| | *Homo erectus* | *Homo erectus* |
| Panins | *Pan troglodytes* | Chimpanzee |
| | *Pan paniscus* | Bonobo |
| Gorilla | *Gorilla gorilla* | Western gorilla |
| | *Gorilla beringei* | Eastern gorilla |
| Orangutans | *Pongo abelii* | Sumatran orangutan |
| | *Pongo pygmaeus* | Bornean orangutan |
| Gibbon | *Nomascus leucogenys* | Northern white-cheeked gibbon |
| Theropithecus gelada | *Theropithecus gelada* | Gelada |
| Macaque | *Macaca mulatta* | Rhesus macaque |
| Baboon | *Papio anubis* | Olive baboon |
| Vervet | *Chlorocebus sabaeus* | Green monkey |
| Colobine | *Colobus angolensis* | Angola colobus |
| Langur | *Rhinopithecus roxellana* | Golden snub-nosed monkey |

Total amino acid coverage per sample for the searches with the different databases were calculated and are displayed Table S10. Noticeably, there is a direct correlation between the size of the database and the sequence coverage. It has been demonstrated that the bigger the database, the less coverage can be obtained, likely due to the exponential increase of the search space. Thus, it was decided to use the reduced but specific “Core Database” for sequence reconstruction.

**Table S10: The choice of the database impacts the search outputs.** Total amino acid coverage per sample per search and the coverage variation compared to the core database.

| **Sample** | **Core Database** | **All Monkeys Database** | **Loss/Core Database** | **All Apes Database** | **Loss/Core Database** |
|---|---|---|---|---|---|
| **SK 830** | 511 | 453 | - 13 % | 430 | - 19 % |
| **SK 850** | 672 | 628 | - 7 % | 478 | - 41 % |
| **SK 14132** | 751 | 720 | - 4 % | 666 | - 13 % |

#### 9.2. Recovery of ancient proteins

The reconstructed sequences correspond to the most abundant proteins in the enamel proteome (eg. AMELX/Y, ENAM, AMBN). The proportion of those proteins were preserved from the modern reference samples to the ancient *Paranthropus* samples (Extended Data Fig. 1 B). We observed the proportion of collagen reduced in the ancient samples, indicating the sampling of dentine in the modern reference or the poor conservation in ancient samples of collagen proteins not localised in the enamel mineral matrix. Moreover, the proportion of enamelin was higher in ancient samples, possibly due to the hydrophilic character of the recovered section of the protein, favouring its preservation [114].

Noticeably, most of the recovered amino acid positions (**425**) were covered in all four *Paranthropus* samples (Extended Data Fig. 1 C). This observation demonstrates the consistency in the recovered sequences and the sequence reconstruction workflow.

Additionally, the length of the recovered peptides was observed to be significantly shorter than in the modern *Homo sapiens* samples (Extended Data Fig. 1 D). This is likely due to diagenetic alteration of the biological material. This observation adds confidence that endogenous peptides were detected.

#### 9.3. Fractionation for increased coverage

High pH reversed-phase fractionation has been demonstrated to be an efficient strategy to increase the depth of coverage of the analysis [115,116] even when sample input amount is limited [117]. The fractionation was manually performed on stage-tips [118] to obtain five fractions total. Manual stage-tip fractionation, in opposition to the use of a HPLC system was used to avoid potential losses in both the autosampler and the fraction collector. Moreover, scaling the amount of stationary phase to the peptide input amount is necessary for optimal sample recovery. Therefore, for these sensitive samples with limited input amount, using less stationary phase (ie. a small StageTip) was preferred. The analysis of the pH adjustment fraction was attempted and led to a blockage of the chromatographic column. Thus, only four fractions could be run using the nano-LC system. The analysis of the pH adjustment fraction was, instead, successful using an Evosep LC system, enabling the acquisition of five fractions for samples SK 850 and SK 14132.

The offline high pH separation of the peptides was orthogonal to the later one performed online at low pH using either the nano-LC or the Evosep system. The fractionation decomplexified the samples prior to MS analysis. That decomplexification led to more spectra being identified, especially in a DDA scheme, where (i) low abundant peptides had more chances of being selected for fragmentation and sequencing and (ii) different ions were less likely to simultaneously be selected for fragmentation leading to chimeric spectra. Moreover, the consecutive analysis of the fractions on the MS increased the total MS acquisition time and provided more chances of acquiring high quality MS2 spectra, leading to an increased number of PSMs, noticeable in all four *Paranthropus* samples. This was directly observed with a gain of 207% of PSMs in sample SK 850 compared to the single shot analysis (Extended Data Fig. 2). The gain was also present with the total amino acid coverage where 17% more positions were covered using high pH stage-tip fractionation. The highest gain at the PSM level compared to that of sequence coverage indicates that the biggest advantage with fractionation is that more PSMs were recorded that could support amino acid identifications, giving more confidence in the reconstructed sequence. However, the increase in sequence coverage is still valuable for analysis, as it maximises the amount of information from a limited sample resource. Off-line fractionation has, therefore, proven to be a suitable approach to increase the depth of enamel palaeoproteomic analysis. However, it does require a higher sample input amount compared to a single analysis.

#### 9.4. Sequence reconstruction

Proteomics sequence reconstruction and its subsequent phyloproteomic analysis is a tedious and time consuming process. Therefore, efforts have been focused on making it systematic and reproducible. For that purpose, a computational pipeline has been developed for sequence assembly and has been used on the *Paranthropus* sampleset.

##### 9.4.1. Computational pipeline for sequence assembly

A computational pipeline for generating a consensus sequence was developed (Extended Data Fig. 4). Reconstructing sequences from output tables from search engines is extensive work that remains subjective and prone to human errors. Therefore, reproducibility and consistency in the process is reduced. A systematic approach can be used with the in-house built R-script for the generation of consensus sequences from MaxQuant´s output tables and an aligned version of the database used for the search in FASTA format as inputs. This strategy enabled a faster and more efficient data analysis process. Moreover, this strategy was also reproducible and transparent with the use of an exclusion list to reference the rejected peptide sequences.

The R script has been built as an R-project. For a simplified use, a folder should be created where the full R-project folder should be saved, along with the MaxQuant txt folder and the aligned database.

The R-project folder contains 3 distinct scripts: Function.R, Prepare.R and Reshape.R. Function.R is used for loading functions that will be used in the other files. Prepare.R loads the relevant MaxQuant tables for further analysis as well as the database, which is then indexed with gene level and protein level positions. Reshape.R processes the tables and generates the output files.

Input: evidence.txt, msms.txt, summary.txt, aligned database (as FASTA file), exclusion list.csv

1. The aligned FASTA file is loaded and expanded from a protein level table to a site level table where each row corresponds to an amino acid position in a protein sequence. The amino acids are thus referenced using the “aligned” and “gapless” positions. Therefore it is easily possible to go from the position referenced in MaxQuant (“gapless”) to a gene position “aligned”.
2. Experimental annotations are extracted from the summary table and added to the evidence and msms.txt files. It is recommended to set up the MaxQuant search using the experiment and fractions annotations. That way, the script would not have to be modified to suit the experimental design.
3. Unique peptide sequences are extracted from the msms.txt table and mapped to all possible protein matches in the database. The table is expanded and each peptide sequence-protein pair is referenced with the start and end position of the peptide at the protein (gapeless) and gene (aligned) levels. The razor principle is then applied to peptide sequences matching multiple genes, where the referred peptides are assigned to the gene with the most peptide identifications.

   Output: *PepSeqs.csv* → gathers unique peptide sequences with unique gene position.
4. The delta score is recalculated from the msms.txt table for the PSMs where the second best identification differs from the first one by a I/L, G/E or N/D. In those cases the delta score is reprocessed as the difference between the score of the first best identification with the score of the third best identification.
5. Facultative step: Identification of the optimal delta score cut off. The algorithm will loop with different delta score cutoffs and apply the subsequent FDR at 1%. A plot representing the number of PSMs as a function of the delta score will be displayed. The optimal delta score

cutoff has to be chosen at the maximum of the curve to maximise the identifications (Supplementary 9.4.2).

6. The chosen delta score cutoff is applied to the msms.txt table, and the identifications are filtered to reach a FDR of 1% per sample. The delta score has been optimised for the *Paranthropus* samples at 15, that value can be modified in the script.
7. The evidence table is filtered based on the re-calculated delta score and the FDR at 1%. The reverse sequences, contaminants, and sequences found in the exclusion list are filtered out. Identifications (peptide sequences, modified peptide sequences and PSMs) are calculated per experiment and per sample.

   Output: *IDbyExp.csv* → table with the identifications per experiment. *IDbySample.csv* → table with the identifications per sample.
8. Entries without genes are filtered out. Entries from the evidence file are collapsed per charge state to obtain unique modified peptides per row. The table is then expanded to the site level. Modified and unmodified amino acids are collapsed to obtain unique unmodified amino acids. Information from overlapping peptides is collapsed on the site level (Figure S7). Absolute amino acid coverage per sample and per experiment can be calculated.

   Output: *AACoverageRawFile.csv* → table with the total amino acid coverage per experiment. *AACoverageSample.csv* → table with the total amino acid coverage per sample. *sites.csv* → table gathering all the covered positions per sample with the median and maximum score, the average mass error, the maximum log10 intensity of all peptides covering that site in that sample, the PSM count and the list of peptide sequences covering that site.
9. For a given sample, more than one amino acid can be identified covering the same position on the gene sequence. Those positions have to be detected to generate a consensus by only keeping the amino acid with the highest coverage. The relative coverage (PSM count) and the relative intensity of the conflicting sites are generated through the ratio between the site count/intensity of the given amino acid and the total count/intensity of the site for one sample. The consensus is based on the most common variant.

   Output: *VarSites.csv* → Displays the list of conflicting sites with their relative coverage.
10. An empty data table is generated where the gaps are filled with dashes. The sites are then pasted together into a consensus sequence by sample.
11. Reference sequences from the database are added to the consensus. The sequences can be exported as FASTA files.

   Output: *Hominid-Consensus_Ref_MQ.fasta* → Consensus sequences for all the samples aligned with the reference sequences. *Paranthropus-Consensus_Ref_MQ.fasta* → Consensus sequences of given samples aligned with the reference sequences. *Paranthropus-Consensus_MQ.fasta* → Consensus sequences of given samples only.

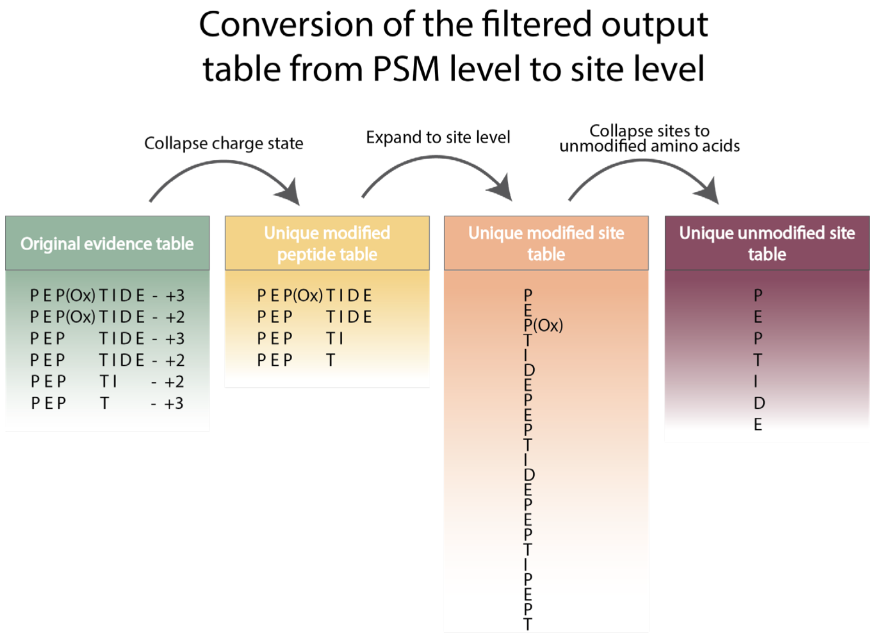


**Figure S7: Tables are converted for sequence assembly**. Schematic representation of the conversion of the filtered evidence table to a site level table using the developed automated sequence assembly script.

##### 9.4.2. Delta score recalculation and optimization

In MaxQuant, the delta score represents the difference between the score of the best matching PSM against the second best matching PSM with a different sequence [119]. This filtering step is applied by default in MaxQuant and is used for the removal of doubtful hits by assessing how different the top identification is compared to a different peptide. Yet, in the case of sequence reconstruction, peptides bearing isobaric residues where both options are present in the database would lead to a delta score of zero. This is the case for I/L but also deamidation products Q/E and N/D. By default, the delta score cutoff in MaxQuant is set at 6, leading to the exclusion of those peptides from the outputs.

The delta score was recalculated using the Andromeda score of the first and third best hits and not the first and second anymore. This approach, while saving identifications, was also blind towards other types of isobaric residues and peptides containing several deamidated sites or a combination of a deamidated site and I/L. After delta score filtering, the dataset was filtered to bring down the FDR to 1% per sample. The PSMs were ordered by increasing score and assigned a p-value. PSMs with p-values < 0.01 were removed [120–122].

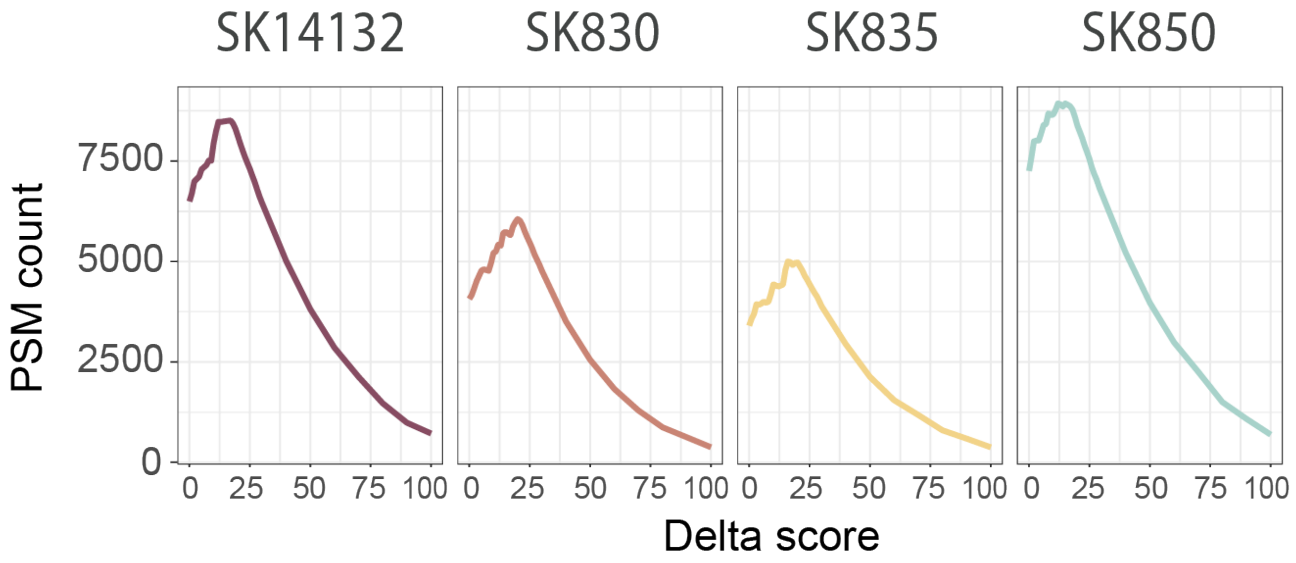


**Figure S8: The optimal delta score should be assessed based on the dataset.** PSM counts as a function of the delta score cutoff applied on the *Paranthropus* samples.

The optimal delta score cutoff was assessed to guarantee the highest PSM coverage (figure S8). The optimal delta score cutoff was set at the inflexion point of the curve, here 15. Noticeably, a low delta score would not increase the number of identified PSMs by enabling more low quality hits among the outputs leading to less PSMs at 1% FDR. This demonstrates the orthogonality of both filtering techniques for statistical purposes. The optimal delta score was identified as 12 for the faunal remains and 6 for the *Paranthropus* samples analysed in South Africa**.**

##### 9.4.3. Generation of a consensus sequence

Different amino acids covering one specific position can be identified among one sample. In those cases, the algorithm will generate the consensus sequence based on the highest relative coverage. It is possible to export the output table containing the conflicting sites for manual examination. Those sites can reveal being covered by falsely identified peptides (artefacts), contaminants, chemical modifications (ie. W→ D) or show heterozygotes among the population. It is important to manually assess the plausibility of the peptide identified to discriminate artefact from heterozygous sites. The list of the identified conflicting positions can be found in the VarSites.csv table.

##### 9.4.4. Alternative search strategies

The pFind [47] search outputs were analysed for the potential identification of new SAPs (single amino acid polymorphisms) that were not included in the database. No new substitutions were detected using this approach. The depth obtained with pFind was on par with the one from MaxQuant (Table S11). No additional manual filtering was performed on the pFind outputs, justifying the higher number of identifications.

**Table S11: Summary of the outputs from the searches using pFind**. The table gathers for each sample the number of PSMs and the identification rate, corresponding to the percentage of identified MS2 spectra.

| Sample | PSM count | Identification rate (%) |
|---|---|---|
| SK 14132 | 6446 | 7.3 |
| SK 830 | 5846 | 5.5 |
| SK 835 | 5018 | 4.6 |
| SK 850 | 8560 | 8.6 |

**Table S12: Output summary from Peaks spider algorithm.** The table shows each sample's MS2 PSM count and the number of identified peptide sequences.

| Sample | PSM count | Peptide Sequences |
|---|---|---|
| SK 14132 | 1621 | 395 |
| SK 830 | 1315 | 305 |
| SK 835 | 411 | 137 |
| SK 850 | 526 | 187 |

#### 9.5. SAP validation

SAP containing regions were thoroughly examined and validated. The following criteria were considered to assess the legitimacy of a polymorphism (i) PSM coverage, (ii) peptide specificity, (iii) ion coverage around the mutated amino acid, (iv) correlation with spectral predictions, (v) potential chemical artefacts, (vi) number nucleotide change necessary.

##### 9.5.1. Individual SAP validation

17 phylogenetically relevant SAPs were identified in the *Paranthropus* samples (Table S13). The detection of those SAPs were manually validated and compared to spectral predictions using Prosit [123] with the HLA model [124] (Supplementary Document 1). Not every SAP was identified in every sample, making the study of several individuals beneficial for a more comprehensive reconstruction.

**Table S13: Summary of the detected site variations across the four *Paranthropus* samples.** This table groups the varying sites detected in the *Paranthropus* samples by gene and position in the gene. The name of the samples in which that site was covered is indicated in the Samples column as well as the number of PSMs covering that site in the PSMs column and the maximum score of a peptide covering that site in any sample is displayed in the Max Score column. Finally the last column indicates the phylogenetic relevance of the site.

| Gene | Position | Samples | PSMs | Max Score | Phylogenetic Signal Importance |
|---|---|---|---|---|---|
| AHSG | 117 | SK 14132 - SK 830 - SK 850 | 4 - 1 - 4 | 63.6 | Differentiates from *Pongo* and non-hominid primates, matches *Gorilla*, *Pan*, *Homo* |
| ALB | 574 | SK 830-SK 14132-SK 835 - SK 850 | 3 - 3 - 9 - 5 | 103.1 | Differentiates from *Gorilla*, matches *Pongo*, *Pan*, *Homo* |
| ALB | 202 | SK 14132 | 4 | 119.6 | Differentiates between *Pan*, *Gorilla*, *Pongo*, matches only with *Homo* |
| AMBN | 270 | SK 830 - SK 14132 - SK 835 - SK 850 | 15 - 9 -5 - 12 | 122.5 | *Differentiates from Pongo and some non-hominid primates, matches Gorilla, Pan,Homo* |
| AMELX | 127 | SK 830 - SK 835 - SK 850 - SK 14132 | 3 - 1 - 2 - 2 | 83.2 | Differentiates from *Pongo*, matches *Gorilla*, *Pan*, *Homo* |
| AMELX | 135 | SK 835 - SK 850 - SK 14132 | 2 - 16 -24 | 83.3 | Differentiates from *Pongo*, matches *Gorilla*, *Pan*, *Homo* |
| AMELY | 101 | SK 835 | 1 | 68.8 | Differentiates from *Pongo*, matches *Gorilla*, *Pan*, *Homo* |
| AMELY | 109 | SK 835 - SK 850 | 1 - 2 | 68.8 | Differentiates from *Pongo*, matches *Gorilla*, *Pan*, *Homo* |
| AMELY | 184 | SK 835 - SK 850 | 8 - 8 | 111.0 | Differentiates from *Pongo*, matches *Gorilla*, *Pan*, *Homo* |
| AMTN | 99 | SK 835 | 1 | 52.6 | Differentiates from *Pongo* and *Gorilla*, matches *Pan*, *Homo* |
| COL17A1 | 636 | SK 835 - SK 14132 | 1-1 | 64.3 | Differentiates from *Homo*, matches everything else in hominids + other primates |
| ENAM | 137 | SK 830 - SK 835 - SK 850 - SK 14132 | 48 - 53 - 28 - 22 | 121.5 | Differentiates from *Homo*, *Pan*, *Gorilla*, matches *Pongo*, differentiates *Paranthropus* individuals |
| ENAM | 147 | SK 830 - SK 835 - SK 850 - SK 14132 | 9 -4 - 30 - 23 | 138.7 | Differentiates from *Pongo* and *Gorilla*, matches *Pan*, *Homo* |
| ENAM | 190 | SK 830 - SK 835 - SK 850 - SK 14132 | 239 - 86 - 249 - 240 | 157.2 | Differentiates from *Pan*, matches *Gorilla*, *Homo* and *Pongo* |
| ENAM | 212 | SK 835 - SK 850 - SK 14132 | 2 - 91 - 109 | 169.8 | Differentiates from *Pongo*, matches *Gorilla*, *Pan*, *Homo* |

| | | | | | |
|---|---|---|---|---|---|
| MMP20 | 292 | SK 830 - SK 850 | 2 - 2 | 50.9 | Differentiates from *Pongo*, matches *Gorilla*, *Pan*, *Homo* |
| ODAM | 271 | SK 830 - SK 835 | 5 - 2 | 123.9 | Differentiates from *Pongo*, matches *Gorilla*, *Pan*, *Homo* |

##### 9.5.2. ENAM-137 validation with synthetic peptides

Synthetic peptides covering the ENAM-137 site were purchased to experimentally retrieve the retention time and fragment intensity information from “true positive” peptides as described by Cappellini et al[41]. The peptide sequence KPPQK**Q**PLK could be identified in the three individuals having a glutamine at ENAM-137: SK 830, SK 850 and SK 14132. Its homologous sequence: KPPQK**R**PLK was identified in SK 835, yet could not be identified in SK 14132. To validate the heterozygous character of SK 14132, a third synthetic peptide covering ENAM-137 with the R version was purchased: QK**R**PLKQP.

The experimental spectra acquired from the *Paranthropus* samples strongly correlated with the spectra obtained from the synthetic peptides (Figure S9, Fig. 3).

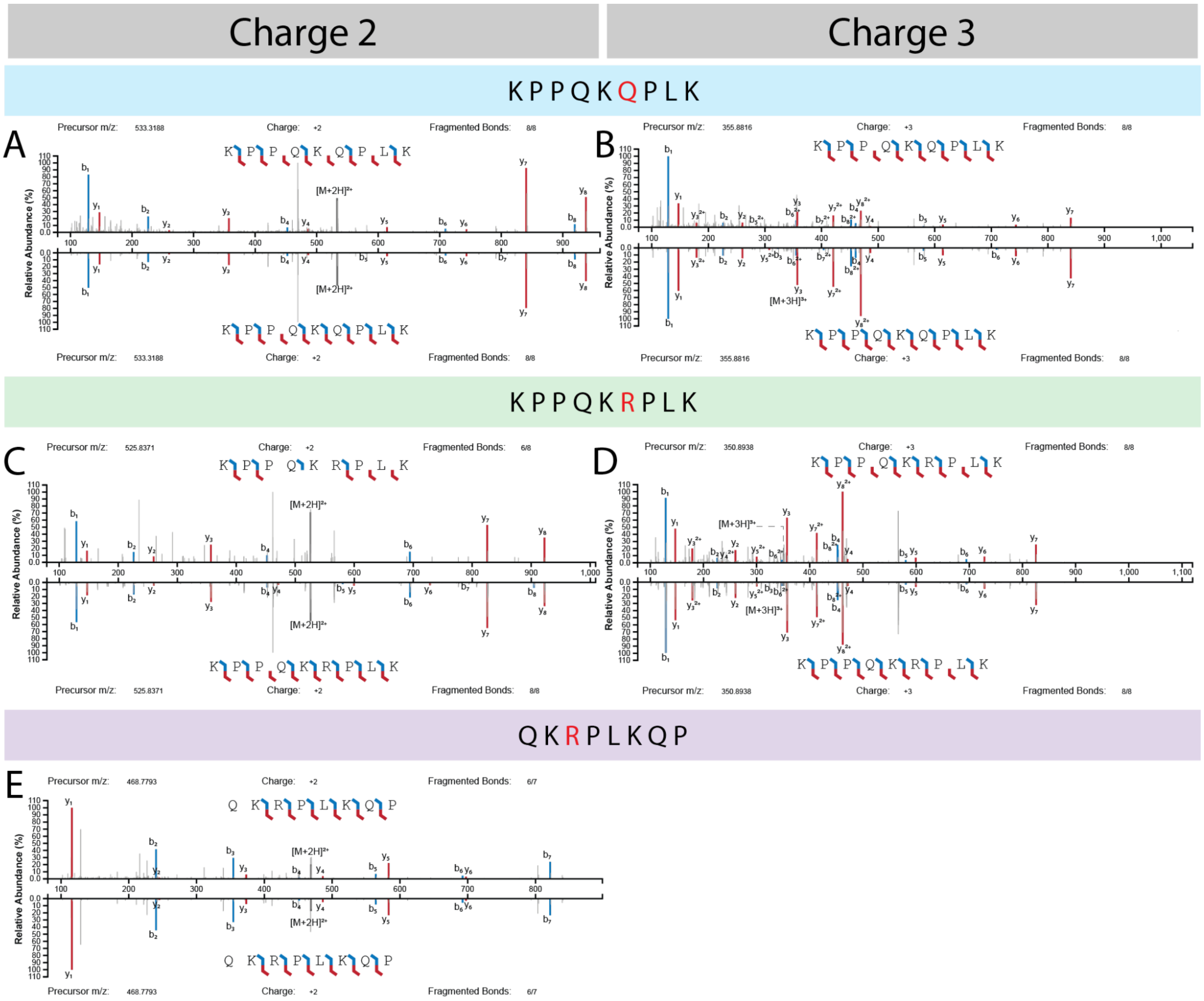


**Figure S9: Validation of ENAM-137 detection using synthetic peptides**. Mirror plot of the identified spectra from the *Paranthropus* dataset (top) and spectra acquired from synthetic peptides (bottom) representing the true fragmentation pattern. A- KPPQ(De)KQ(De)PLK, charge 2. B - KPPQ(De)KQ(De)PLK, charge 3. C - KPPQ(De)KR(Orn)PLK, charge 2. D - KPPQ(De)KR(Orn)PLK, charge 3. E - Q(Glu)KR(OrnPLKQ(De)P, charge 2.

Moreover, the retention times of the synthetic and endogenous peptides coincided with each other (Table S14). The hydrophilic nature of the peptides make them elute early in the gradient. In extreme parts of the gradients, the variability in retention times is higher, explaining the increased variability for the peptides 1 and 3 eluting the earliest. Moreover, the peptides identified in the *Paranthropus* samples were in a different matrix, making an accurate retention time comparison challenging [125,126].

Additionally, those peptides were identified bearing age related modifications (ie. Deamidation and arginine to ornithine conversion), adding confidence in their identification and the endogenous character of the sequences.

**Table S14: Retention time comparison with the synthetic peptides.** Information about the chosen peptides identified in the dataset (i.e., *Paranthropus*) along with the retention time of the synthetic peptides and the corresponding deviation.

| Peptide | Charge | m/z | PSM | Mean RT (min) | RT synthetic peptide (min) | Deviation (%) | Max Score | Sample |
|---|---|---|---|---|---|---|---|---|
| KPPQ(De)KQ(De)PLK | 2 | 533.3188 | 6 | 7.17 | 8.54 | 16 | 98.629 | SK 830 - SK 850 - SK 14132 |
| KPPQ(De)KQ(De)PLK | 3 | 355.8816 | 8 | 7.73 | 8.54 | 9 | 84.213 | SK 830 - SK 850 - SK 14132 |
| (gl)QKR(ar)PLKQ(de)P | 2 | 468.7793 | 4 | 10.72 | 10.70 | 0.1 | 69.998 | SK 835 - SK 14132 |
| KPPQKR(ar)PLK | 3 | 350.8938 | 5 | 6.57 | 5.84 | 12.5 | 97.635 | SK 835 |
| KPPQKR(ar)PLK | 2 | 525.8371 | 1 | 6.15 | 5.84 | 16 | 47.082 | SK 835 |

##### 9.5.3. Specificity of the peptides

Peptides extracted from ancient samples have been altered through time. The main alterations are the modification of amino acid residues (PTMs) and hydrolysis. Endogenous enzymes are present in the enamel[41]. As a consequence, the enamel proteome will consist mostly of peptides. Yet, those peptides will be further hydrolyzed and thus shortened over time. This can be seen from the peptide length distribution of the *Paranthropus* samples that were shorter than the modern samples (Figure S10, Table S15).

**Table S15: Peptide length decreases with age**. PSM count, mean peptide length and approximate age of the available samples.

| Sample | *H. antecessor* | *H. erectus* | *Gigantopithecus* | *Gorilla* | *H. sapiens* | SK 14132 | SK 830 | SK 835 | SK 850 | *Pongo* |
|---|---|---|---|---|---|---|---|---|---|---|
| Count | 6774 | 1868 | 1229 | 9418 | 6738 | 4194 | 3307 | 2808 | 4409 | 11297 |
| Mean | 11.58 | 9.54 | 9.01 | 12.8 | 12.55 | 9.3 | 9.29 | 9.16 | 9.25 | 12.6 |
| Age (Ma) | 0.8 | 1.77 | 1.9 | 0 | 0 | 2 | 2 | 2 | 2 | 0 |

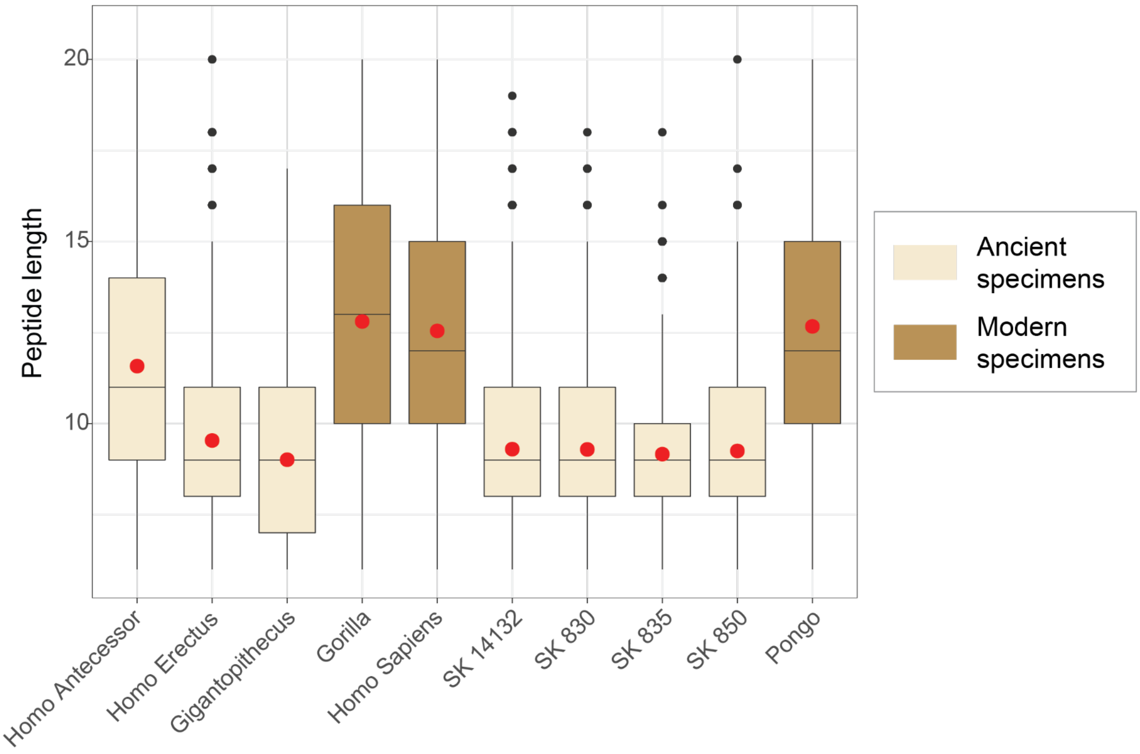


**Figure S10: The peptide length is lower in ancient samples compared to modern references.** Peptide length distribution of the different specimens studied, including ancient specimens as well as modern reference samples. The red dot represents the average peptide length.

The reduction of the size of the detected peptides introduces an additional challenge and hinders the specificity of the peptides. Sites were sometimes covered by very short peptides, and those peptides were unspecific. Yet, as a guide, it is discouraged to keep those peptides as evidence since it could be exogenous contamination.

Peptides bearing age-related modifications and matching with few bacteria or other non enamel-specific sequences were likely to be endogenous and to actually support the reconstructed protein sequence. An additional search was carried out, including in the database the bacterial proteins whose peptides (Table S16) could potentially match the enamel proteins. After protein filtering with a FDR at 1 %, the additional sequences were not identified. As a consequence, the peptides (Table S16) were considered part of the enamel proteins and kept for sequence reconstruction.

**Table S16: List of potential contaminants matching with the identified peptide sequences**. The proteins were added to the extended database.

| **Gene** | **Position** | **Potential matches** |
|---|---|---|
| **AHSG** | **117** | WP_233222100.1 sigma-70 family RNA polymerase sigma factor [Sphingomonas deserti] |
| | | MCQ2286586.1 MAG: hypothetical protein MJZ76_06905 [Bacteroidales bacterium] |
| | | RMD88154.1 MAG: DUF4440 domain-containing protein [Alphaproteobacteria bacterium] |
| | | PSJ42099.1 RNA polymerase subunit sigma-24 [Sphingomonas deserti] |
| | | UTC91503.1 TetR/AcrR family transcriptional regulator [Treponema denticola] |
| | | WP_255818665.1 TetR family transcriptional regulator [Treponema putidum] |

| | | |
|---|---|---|
| | | WP_044978826.1 TetR/AcrR family transcriptional regulator [Treponema putidum] |
| | | MBV5314908.1 MAG: HAMP domain-containing histidine kinase [Prolixibacteraceae bacterium] |
| | | MBL7848187.1 MAG: hypothetical protein JNL40_12005 [Cyclobacteriaceae bacterium] |
| **ALB** | **574** | GHJ61412.1 hypothetical protein NOK12_39300 [Nocardioides sp. OK12] |
| | | WP_261821250.1 hypothetical protein [Nocardioides sp. OK12] |
| | | WP_227630751.1 hypothetical protein, partial [Klebsiella pneumoniae] |
| **ALB** | **202** | WP_261821250.1 hypothetical protein [Nocardioides sp. OK12] |
| | | WP_084287698.1 RNA-directed DNA polymerase [Desulfovermiculus halophilus] |
| | | GHJ61412.1 hypothetical protein NOK12_39300 [Nocardioides sp. OK12] |
| | | MCL2627072.1 MAG: hypothetical protein FWD44_00030 [Oscillospiraceae bacterium] |
| | | WP_135114708.1 amino acid ABC transporter permease [Microbacterium paludicola] |
| | | MCJ7777353.1 MAG: ATP-grasp domain-containing protein [Sedimentisphaerales bacterium] |
| | | WP_055993351.1 amino acid ABC transporter permease [Microbacterium sp. Root53] |
| | | WP_134745790.1 amino acid ABC transporter permease [Microbacterium sp. MEC084] |
| | | OYV86766.1 MAG: hypothetical protein B7Z63_03485 [Ignavibacteriae bacterium 37-53-5] |
| **AMTN** | **99** | CAH2059869.1 unnamed protein product [Thlaspi arvense] |
| **COL17A1** | **636** | MCI9320541.1 MAG: collagen-like protein [Lachnospiraceae bacterium] |
| | | MCI9124705.1 MAG: collagen-like protein [Eubacterium sp.] |
| | | SOY28350.1 Collagen triple helix repeat (20 copies) [Acetatifactor muris] |
| | | RKJ18300.1 collagen-like protein [bacterium D16-50] |
| **MMP20** | **292** | WP_255037882.1 PIN domain-containing protein [Lacihabitans soyangensis] |
| | | WP_255077629.1 PIN domain-containing protein [Lacihabitans sp. CCS-44] |
| | | KAI9065413.1 hypothetical protein FKP32DRAFT_1685007 [Trametes sanguinea] |
| | | OSD08473.1 hypothetical protein PYCCODRAFT_1473077 [Trametes coccinea BRFM310] |
| | | RKO90250.1 hypothetical protein BDK51DRAFT_48489 [Blyttiomyces helicus] |

##### 9.5.4. Fragment intensity predictions

Peptide sequences can be retrieved on MS2 spectra by identifying the mass difference between successive fragment ions[127,128]. Yet, different fragment ions can exhibit different stability and lead to unique fragmentation patterns [127,129–131]. Therefore, based on the chemistry of the peptides, some fragment ions can be absent from MS2 spectra due to low stability or poor fragmentation properties. A complete ion series can thus not be retrievable for every peptide. Yet, for sequence reconstruction, it is advised to have consecutive ions covering the SAP for unambiguous identification. We took advantage of newly developed deep learning approaches to add fragment intensity predictions as an additional layer of confidence in the SAP identifications [123,124,132].

Using synthetic peptides enabled an unbiased validation of the identified SAP, yet it remains expensive, both with the purchase of the peptides and the additional mass spectrometric time required. Furthermore, this strategy was time consuming and did not enable a rapid validation or refutation of the identified peptide. It was thus challenging to implement such an approach for all possible SAPs in a dataset.

The empirical method using synthetic peptides was compared to newly developed deep learning strategies for fragmentation pattern predictions with models trained on non-tryptic peptides[123,124]. Currently, Prosit is able to accurately predict fragmentation spectra from tryptic and non-tryptic peptides bearing carbamidomethylation of cysteine, methionine oxidation and TMT (Tandem Mass Tag) labels. Yet, the current models were unable to support other types of modifications, like phosphorylation.

To benchmark the approach, we used reference peptides used by Cappellini et al.[41] where they acquired MS2 spectra for 21 synthetic peptides. 3 of those peptides contain phosphorylation making the fragmentation prediction impossible with the currently available models. 5 peptides were tryptic whereas the 13 remaining were unspecific (Table S17).

**Table S17: Spectral predictions benchmarked using synthetic peptides.** List of synthetic peptides sequenced by Cappellini et al. and here used to benchmark the Prosit predictions on enamel peptides.

| Peptide sequence | Mass | Charge |
|---|---|---|
| PPPN(De)TRR | 419.7301 | 2 |
| DKPVLQ(De)K | 414.7449 | 2 |
| GPRKTFPG | 430.2426 | 2 |
| PLKAEPDD | 442.7216 | 2 |
| VEESSFKE | 477.7244 | 2 |
| GVEESSFKE | 506.3251 | 2 |
| PQ(De)Q(De)PGQ(De)KPF | 515.248 | 2 |

| | | |
|---|---|---|
| LKAEPDDL | 450.7373 | 2 |
| DELKPLVD | 464.7529 | 2 |
| KELLLFR | 459.7922 | 2 |
| PPPAQ(De)Q(De)PFQ(De)PQ(De) | 619.7824 | 2 |
| DDSPN(De)FPPLK | 565.7718 | 2 |
| LFPYHQ(De)PL | 508.266 | 2 |
| AVLEDFA | 382.6949 | 2 |
| FQ(De)DAKDVFL | 452.2715 | 2 |
| FPPPAQ(De)Q(De)PFQ(De) | 580.2689 | 2 |
| VLEDFAAFLN(De)K | 634.3321 | 2 |
| FDAVTM(Ox)LGKELLL | 733.4022 | 2 |

The prediction of the 18 studied peptides showed excellent correlation with the experimental spectra acquired from the synthetic peptides (Supplementary Document 2). For the generation of the prediction, a deamidated glutamine residue was substituted by glutamic acid and, likewise, deamidated asparagine was substituted by aspartic acid. The accurate predictions generated by the deep learning strategies demonstrated their applicability in palaeoproteomics for peptide validation. Accordingly, prediction tools were used as additional evidence for peptide validation.

#### 9.6. Sex identification and confidence in the female attribution

The biological sex of the *Paranthropus* individuals were primarily inferred by the presence or absence of AMELY specific peptides. The samples SK 850 and SK 835 were assigned as male individuals due to the unambiguous presence of AMELY-specific peptides (Figure 2). This presence was confirmed by the identification of multiple overlapping peptides. All overlapping peptides identified are accessible from the output files of the sequence assembly script in the 'Site table. csv' file, available on Pride (PXD040221) and on GitHub (https://github.com/ClaireKoenig/ProteinSequenceAssembly). Yet, the absence of those AMELY-specific peptides is weak evidence for the assignment of the other individuals as females. Therefore, to strengthen this assignment, we looked at the site intensity of AMELY-59 (male) versus AMELX-60 (male and female) for the four *Paranthropus* samples.

The site intensity represents the sum of the intensities of all PSMs covering that site. The site intensities of AMELY-59 as a function of AMELX-60 is displayed in (Figure 2). The horizontal cut-off represents the minimum intensity of an AMELY-specific peptide covering the AMELY-59 site in a male individual. The vertical cut-off represents the minimum intensity of a peptide covering the AMELX-60 site in a male individual. Here we are assuming that the AMELX to AMELY ratio remains constant across individuals. Therefore, if a sample has an AMELX-60 site intensity above the vertical cutoff line, we would expect the detection of AMELY-specific peptides to be possible from a technical point of view. Both SK 830 and SK 14132 demonstrate AMELX-60 site intensities above the vertical cutoff. Thus we can strongly assume that the absence of AMELY-specific peptides from those individuals is not due to the mass spectrometer's limit of detection. We can then strongly assume that the observed variation is biological and thus, SK 830 and SK 14132 can be assigned as probable females.

#### 9.7. Modification analysis

Successive searches were carried out using MaxQuant to estimate the modification rate of different types of molecular damage in the *Paranthropus* samples. The extent of the modification was estimated using PSM counting [41].

The conversion of arginine residues to ornithine has been observed in high numbers. The modification rate in the modern control was at 1% and this increased to at least 39% in the *Paranthropus* samples (Extended Data Fig. 1). Moreover the observed rate among the four individuals was similar, with a standard deviation of 2.2 %. The conversion of arginine to ornithine and the loss of the reactive guanidino group is coherent with the age-induced degradation process (Extended Data Fig. 1 ).

Oxidation has also been demonstrated to be diagenetically induced [41]. Therefore oxidative products of the aromatic amino acids (histidine (H), phenylalanine (F), tyrosine (Y) and tryptophan (W)) have been investigated.

- **Histidine** oxidation, di-oxidation as well as histidine conversion to aspartic acid and to hydroxyglutamate could be observed in both the modern reference and in the *Paranthropus* samples (Figure S11). In comparison to the modern reference, the level of histidine di-oxidation was not significantly increased. On the other hand, aspartic acid and hydroxyglutamate products exhibited a higher rate of conversion compared to the modern reference. Yet, histidine modifications remained extremely non-abundant, making an accurate estimation challenging.

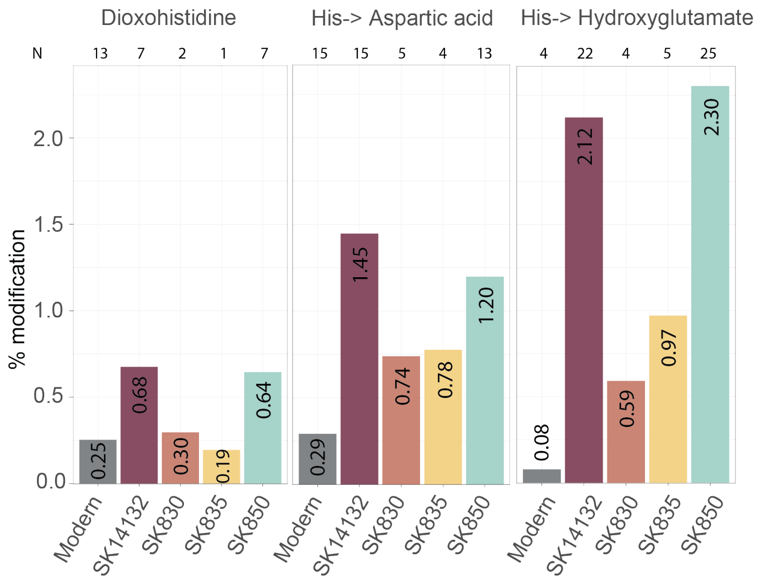


**Figure S11: Histidine modifications.** A - Extent of histidine di-oxidation. B - Extent of histidine conversion to aspartic acid. C - Extent of histidine conversion to hydroxyglutamate. N represents the number of modified sites detected per sample.

- **Phenylalanine** mono and di-oxidation has been observed in the *Paranthropus* samples (Figure S12 A). The extent of phenylalanine oxidation was higher in the ancient samples compared to the modern reference, even though the low number of identifications cannot lead to an accurate estimation.

- **Tyrosine** oxidation and di-oxidation has also been observed in the *Paranthropus* samples (Figure S12 B). Both the rates of mono and di-oxidation were higher than in the modern control. It is assumed that tyrosine oxidative products were generated through diagenetic processes[41]. This observation was consistent with the recovery of ancient enamel protein sequences.

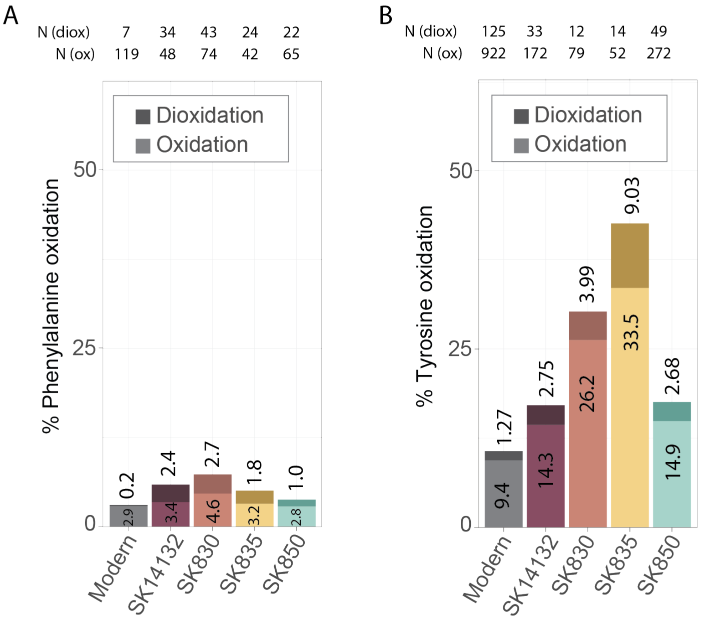


**Figure S12: Phenylalanine and tyrosine modifications.** A - Extent of mono and di-oxidation of phenylalanine. B - Extent of mono- and di- oxidation of tyrosine. N represents the number of modified sites detected per sample.

- **Tryptophan** is observed to be heavily modified through time [41]. In fact, most of the tryptophan residues in the *Paranthropus* samples were modified (Figure S13). Oxidation and di-oxidation were the most represented modifications. In three out of four samples, the rate of oxidation was higher than the rate of di-oxidation. Yet, in comparison with the modern reference, the oxidation rate was up to 1.3 fold higher in the *Paranthropus* samples. With di-oxidation, the lowest sample is already at least 3 fold higher than the modern control. This observation suggests that most of the observed tryptophan mono-oxidation could originate from sample preparation. The higher di-oxidation level in the ancient material would thus likely originate from sample degradation. The most abundant advanced oxidative product was oxolactone, followed by kynurenine, which both exhibit higher modification rates in the ancient material. On the other hand, no conclusion can be drawn for tryptophandione.

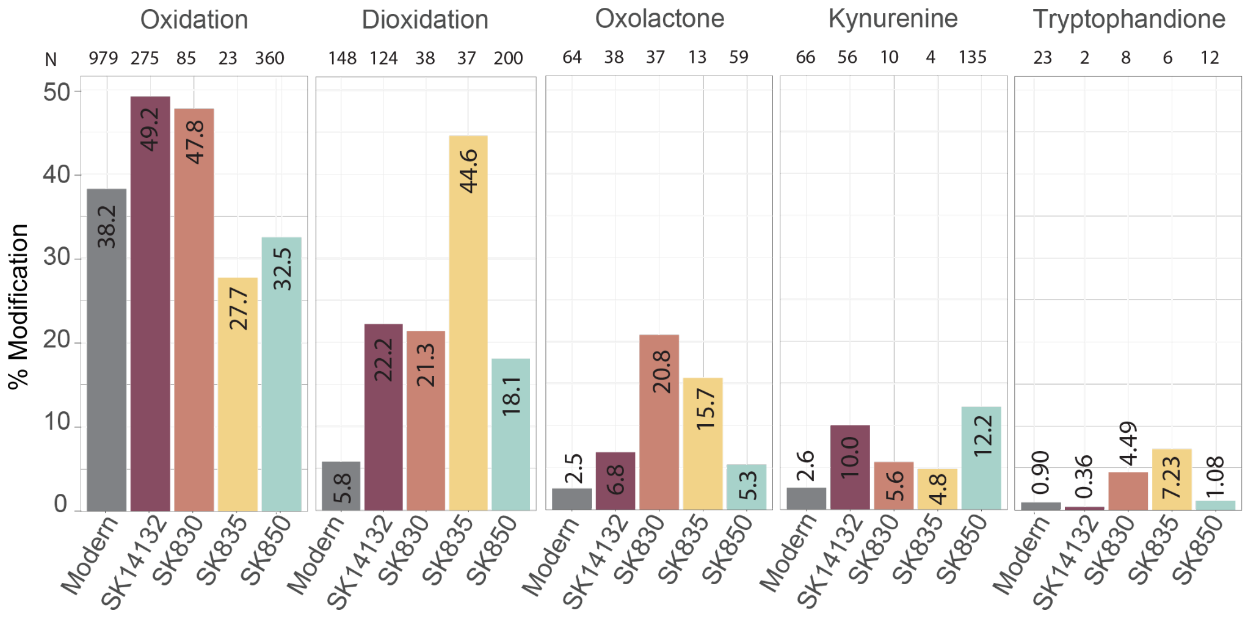


**Figure S13: Tryptophan modifications**. A - Extent of mono-oxidation of tryptophan. B - Extent of di-oxidation of tryptophan. C - Extent of tryptophan conversion to oxolactone. D - Extent of tryptophan conversion to kynurenine. E - Extent of tryptophan conversion to tryptophandione. N represents the number of modified sites detected per sample.

All samples showed similar modification patterns and modification rates. Variability between samples was increased when the depth was reduced, noticeable with a lower number of PSMs, making an accurate estimation more challenging. Yet, interestingly, the individual SK 835 showed some higher tyrosine oxidation and di-oxidation levels than the other samples and exhibited a lower rate of tryptophan mono-oxidation along with a higher rate of tryptophan di-oxidation. Higher di-oxidation in this sample could suggest that the already oxidised residues could have been further oxidised. Altogether, this variation in the oxidation level of SK 835 could indicate a different conservation state (ie. different storage environment)[48]. Yet, that sample has already been extensively studied and this variation could also potentially originate from the multiple scanning techniques employed.

#### 9.8. Cross-linking analysis in ancient enamel

The data acquired from the ancient enamel samples were also searched for cross-linked peptides. The detection and identification of such peptides would be significant, as it could increase the protein sequence coverage through the identification of additional peptide sequences undetectable using standard bottom-up proteomics analysis workflows. This thereby increases the opportunities for the identification of phylogenetically informative amino acid substitutions. The cross-linking searches were targeted to the three most abundant proteins identified in the data, ENAM, ABMN, and AMELX. Specific protein–protein interactions of these proteins has been suggested in the literature, for example, during the secretory stage of enamel formation [133], but how these enamel matrix components interact with one another to form an assembled matrix remains unclear. Amelogenin-ameloblastin complexes have been described as potential functional entities at the early stage of enamel mineralization [133], and stepwise hierarchical self-assembly for AMEL have also been described [134]. However, there is very little information in the literature about the occurrence and type of cross-linking that may occur in enamel proteins. In this work, a variety of cross-linked structures were searched, including naturally occurring cross-links such as disulfide bonds, the enzymatic cross-link between glutamine and lysine reported in modern enamel samples using enzyme activity assays and immunohistochemistry [135], and protein cross-links induced by oxidative reactions observed in other mineralized tissues [136]. Despite the extensive search through the cross-linking search tools and manual searches, no cross-linked peptides were identified. There are potentially many reasons for this, but the main reason may be related to the S/N for cross-linked peptides compared to that of non-cross-linked peptides. Further investigations into this topic, such as an enrichment of cross-linked proteins during the sample preparation, would greatly advance the identification of cross-linked peptides in ancient samples and hopefully increase sequence coverage.

#### 9.9. Samples analysed in South Africa

To replicate the palaeoproteomics workflow in a different lab and foster a collaboration with local skills, three samples (SK 14132, SK 850 and SK 830) were re-analysed at the D-CYPHR regional hub in Cape Town, South Africa. Peptides were separated on an Evosep One system using the commercial 20SPD gradient and sequenced using a Q-Exactive mass spectrometer. The raw files were processed in the same way using MaxQuant and the data was analysed using the in-house developed R-script.

Here, we compared the total amino acid coverage of the three single shots processed in South Africa with the three single shots analysed in Copenhagen for the corresponding samples (Figure S14). Noticeably, the vast majority of amino acid positions recovered from the samples analysed in South Africa overlap those recovered from in Copenhagen. This attests to the consistency of the results and demonstrates that the analysis is reproducible. Yet, the coverage obtained in Copenhagen from the single shot analysis of the 3 samples was higher than the one obtained from the South African raw files with 358 additional positions covered. From this experiment, we can conclude that the use of a more sensitive mass spectrometer, combined with a higher resolution acquisition method leads to an increase in coverage, which is valuable for palaeoproteomic applications. This experiment demonstrated the possibility of recovering endogenous peptide sequences from another laboratory with a different instrumental set-up. Emphasis was placed on the engagement and skill development of the local scientific community. Proteomics analysis is well implemented in South Africa, however not in a palaeoproteomics context.

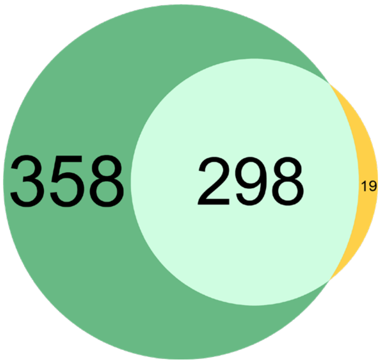


**Figure S14: *Paranthropus* samples were analysed both in Denmark and in South Africa.** Venn diagram of the amino acid position covered in the reconstructed protein sequences. In green: CPH only - in yellow: SA only - in light green: identified in both.

### 10. Phylogenetic reconstruction and variation analysis of *Paranthropus* specimens

#### 10.1. Protein Reference Datasets

All reference datasets created for the phylogenetic analysis are available at: https://zenodo.org/record/7801259 , under the folder 'Reference_Datasets_Unaligned'.

#### 10.2 Alignment Results

##### 10.2.1 Alignment statistics

The aligned and corrected datasets were scanned for informative sites through Paleoprophyler. A table was generated (Tables S18,S19) for each sample describing the number of sites in comparison with the reference dataset. Each sample covers roughly 10% of all available informative sites (not including singletons) of the proteins recovered for it. Although the number of polymorphic/informative sites greatly increases in the 'Diversity' dataset, the coverage of those sites by the ancient samples remains roughly the same. Most of these 'added' polymorphic sites of the 'Diversity' dataset express within-species diversity and do not provide additional information for phylogenetic placement. Note that the numbers for both datasets were calculated by including the outgroups of *Hylobates* and *Macaca* and could change if only members of Hominidae were included.

**Table S18: Statistics obtained from the protein alignments using the 'Representative' reference dataset.** The following numbers were calculated for each sample and for each protein (from left to right): a) Number of total amino acids sites of that protein, b) Number of amino acids covered in the ancient sample, c) Number of polymorphic sites in the whole dataset, d) Number of polymorphic sites covered by the ancient sample, e) Number of polymorphic sites in the whole dataset present in more than one individual, f) Number of polymorphic sites in the whole dataset present in more than one individual and covered by the ancient sample, g) Polymorphic sites where the ancient sample presents a unique polymorphism.

| Sample | Protein | Total sites | Sites present in Ancient Sample | Polymorphic Sites | Polymorphic Sites Covered by Ancient Sample | Polymorphic sites (Non singletons) | Polymorphic sites covered by ancient Sample (non singletons) | Unique Sites for ancient Sample |
|---|---|---|---|---|---|---|---|---|
| SK 830 | ENAM | 1142 | 134 | 141 | 11 | 31 | 4 | 0 |
| | COL17A1 | 1513 | 32 | 111 | 2 | 24 | 0 | 0 |
| | AMBN | 447 | 99 | 65 | 13 | 16 | 4 | 0 |
| | AHSG | 368 | 21 | 67 | 3 | 29 | 1 | 0 |
| | ALB | 609 | 53 | 53 | 7 | 16 | 2 | 0 |
| | ODAM | 279 | 15 | 45 | 6 | 14 | 1 | 0 |
| | AMELX | 205 | 158 | 5 | 4 | 1 | 1 | 0 |
| | MMP20 | 483 | 34 | 34 | 6 | 8 | 1 | 0 |
| | **Total** | **5046** | **546** | **521** | **52** | **139** | **14** | **0** |
| SK 835 | AMELY | 206 | 39 | 26 | 7 | 10 | 1 | 0 |
| | ENAM | 1142 | 135 | 141 | 11 | 31 | 4 | 0 |
| | COL17A1 | 1513 | 34 | 111 | 3 | 24 | 1 | 0 |
| | AMBN | 447 | 99 | 65 | 10 | 16 | 3 | 0 |
| | AMTN | 209 | 21 | 27 | 1 | 11 | 1 | 0 |
| | ALB | 609 | 67 | 53 | 9 | 16 | 2 | 0 |
| | AMELX | 205 | 148 | 5 | 5 | 1 | 1 | 0 |
| | MMP20 | 483 | 43 | 34 | 5 | 8 | 0 | 0 |
| | **Total** | **4814** | **586** | **462** | **51** | **117** | **13** | **0** |
| SK 850 | AMELY | 206 | 47 | 26 | 5 | 10 | 1 | 0 |
| | ENAM | 1142 | 162 | 141 | 14 | 31 | 4 | 0 |
| | COL17A1 | 1513 | 33 | 111 | 2 | 24 | 0 | 0 |
| | AMBN | 447 | 132 | 65 | 18 | 16 | 4 | 0 |
| | AMTN | 209 | 26 | 27 | 0 | 11 | 0 | 0 |
| | AHSG | 368 | 23 | 67 | 4 | 29 | 2 | 0 |
| | ALB | 609 | 60 | 53 | 11 | 16 | 4 | 0 |
| | COL1A1 | 1505 | 17 | 61 | 0 | 2 | 0 | 0 |
| | AMELX | 205 | 182 | 5 | 5 | 1 | 1 | 0 |
| | MMP20 | 483 | 48 | 34 | 10 | 8 | 2 | 0 |
| | **Total** | **6687** | **730** | **590** | **69** | **148** | **18** | **0** |
| SK 14132 | ENAM | 1142 | 171 | 141 | 15 | 31 | 4 | 0 |
| | COL17A1 | 1513 | 51 | 111 | 4 | 24 | 1 | 0 |
| | AMBN | 447 | 133 | 65 | 15 | 16 | 4 | 0 |
| | AMTN | 209 | 26 | 27 | 0 | 11 | 0 | 0 |
| | AHSG | 368 | 25 | 67 | 3 | 29 | 1 | 0 |
| | ALB | 609 | 89 | 53 | 12 | 16 | 5 | 0 |
| | COL1A1 | 1505 | 73 | 61 | 0 | 2 | 0 | 0 |
| | AMELX | 205 | 184 | 5 | 5 | 1 | 1 | 0 |
| | MMP20 | 483 | 29 | 34 | 2 | 8 | 0 | 0 |
| | **Total** | **6481** | **781** | **564** | **56** | **138** | **16** | **0** |

**Table S19: Statistics obtained from the protein alignments using the ‘Diversity’ reference dataset.** The following numbers were calculated for each sample and for each protein (from left to right): a) Number of total amino acids sites of that protein, b) Number of amino acids covered in the ancient sample, c) Number of polymorphic sites in the whole dataset, d) Number of polymorphic sites covered by the ancient sample, e) Number of polymorphic sites in the whole dataset present in more than one individual, f) Number of polymorphic sites in the whole dataset present in more than one individual and covered by the ancient sample, g) Polymorphic sites where the ancient sample presents a unique polymorphism.

| Sample | Protein | Total sites | Non-missing sites in Ancient Sample | Polymorphic Sites | Polymorphic Sites Covered by Ancient Sample | Polymorphic sites Non singletons | Polymorphic sites covered by ancient Sample (not singletons) | Unique Sites for ancient Sample |
|---|---|---|---|---|---|---|---|---|
| SK 830 | ENAM | 1142 | 134 | 165 | 13 | 145 | 11 | 0 |
| | AMBN | 447 | 99 | 74 | 16 | 58 | 11 | 0 |
| | COL17A1 | 1526 | 32 | 183 | 2 | 94 | 2 | 0 |
| | AMELX | 205 | 158 | 9 | 7 | 5 | 4 | 0 |
| | ODAM | 279 | 15 | 59 | 7 | 47 | 6 | 0 |
| | MMP20 | 483 | 34 | 42 | 6 | 29 | 5 | 0 |
| | ALB | 609 | 53 | 60 | 9 | 48 | 8 | 0 |
| | AHSG | 371 | 21 | 79 | 3 | 62 | 3 | 0 |
| | **Total** | **5062** | **546** | **671** | **63** | **488** | **50** | **0** |
| SK 835 | ENAM | 1142 | 135 | 165 | 12 | 145 | 11 | 0 |
| | AMBN | 447 | 99 | 74 | 13 | 58 | 9 | 0 |
| | COL17A1 | 1526 | 34 | 183 | 3 | 94 | 3 | 0 |
| | AMELX | 205 | 148 | 9 | 8 | 5 | 4 | 0 |
| | AMELY | 206 | 39 | 28 | 7 | 17 | 4 | 0 |
| | MMP20 | 483 | 43 | 42 | 5 | 29 | 3 | 0 |
| | ALB | 609 | 67 | 60 | 11 | 48 | 9 | 0 |
| | AMTN | 209 | 21 | 41 | 1 | 33 | 1 | 0 |
| | **Total** | **4827** | **586** | **602** | **60** | **429** | **44** | **0** |
| SK 850 | ENAM | 1142 | 162 | 165 | 16 | 145 | 14 | 0 |
| | AMBN | 447 | 132 | 74 | 22 | 58 | 17 | 0 |
| | COL1A1 | 1510 | 17 | 94 | 0 | 13 | 0 | 0 |
| | COL17A1 | 1526 | 33 | 183 | 2 | 94 | 2 | 0 |
| | AMELX | 205 | 182 | 9 | 9 | 5 | 5 | 0 |
| | AMELY | 206 | 47 | 28 | 5 | 17 | 2 | 0 |
| | MMP20 | 483 | 48 | 42 | 11 | 29 | 9 | 0 |
| | ALB | 609 | 60 | 60 | 13 | 48 | 11 | 0 |
| | AHSG | 371 | 23 | 79 | 4 | 62 | 4 | 0 |
| | AMTN | 209 | 26 | 41 | 1 | 33 | 1 | 0 |
| | **Total** | **6708** | **730** | **775** | **83** | **504** | **65** | **0** |
| SK 14132 | ENAM | 1142 | 171 | 165 | 17 | 145 | 15 | 0 |
| | AMBN | 447 | 133 | 74 | 18 | 58 | 13 | 0 |
| | COL1A1 | 1510 | 73 | 94 | 1 | 13 | 0 | 0 |
| | COL17A1 | 1526 | 51 | 183 | 4 | 94 | 3 | 0 |
| | AMELX | 205 | 184 | 9 | 9 | 5 | 5 | 0 |
| | MMP20 | 483 | 29 | 42 | 2 | 29 | 1 | 0 |
| | ALB | 609 | 89 | 60 | 14 | 48 | 12 | 0 |
| | AHSG | 371 | 25 | 79 | 3 | 62 | 3 | 0 |
| | AMTN | 209 | 26 | 41 | 1 | 33 | 1 | 0 |
| | **Total** | **6502** | **781** | **747** | **69** | **487** | **53** | **0** |

##### 11.2.2 Informative sites of the ancient samples

- **Novel amino acid substitutions:**

When inspecting all four *Paranthropus* individuals, no amino acid substitutions identified for any of the samples were unique or 'novel' (ie. not present in the known biological variation represented in the enamel proteins databases) .

- **Comparisons with Modern Humans:**

In any of the samples, a maximum of two substitutions were identified that differentiate between one of the *Paranthropus* individuals and a member of the *Homo* clade (Modern humans, Neanderthals, Denisovans). These two substitutions are COL17A1-636 and ENAM-137 (protein name - amino acid position in alignment).

COL17A1-636 was covered in two of the four individuals (SK835, SK14132) and both of them share the ancestral SAP with all other Great Apes (Figure S15 A). This position corresponds to the *Homo sapiens* Ensembl reference transcript ID: ENST00000648076.2, which is labelled as the canonical isoform of COL17A1.

The detection of glutamine in ENAM at position 137 using mass spectrometry does not exclude the possibility of having a glutamic acid (E) residue at that position given that a deamidated glutamine bears the same mass as a glutamic acid residue. In the *Paranthropus* samples, all the PSMs contained deamidated glutamine. Yet, deamidation is correlated with degradation and deamidated residues will more likely be detected in ancient samples compared to non deamidated ones (Extended Data Fig.1). Moreover, the *Pongo* lineage bears a glutamine at ENAM-137, making it evolutionary feasible,  while no other primate bears an E at that position. On the other hand, all other African great apes bear an R in that position and a change from R to E would require at least two substitutions on the DNA level, which makes it less likely than the single substitution required for a Q. We thus consider glutamine more likely to be present at ENAM-137 compared to glutamic acid and proceed with the analysis considering glutamine at ENAM-137 in the *Paranthropus* samples.

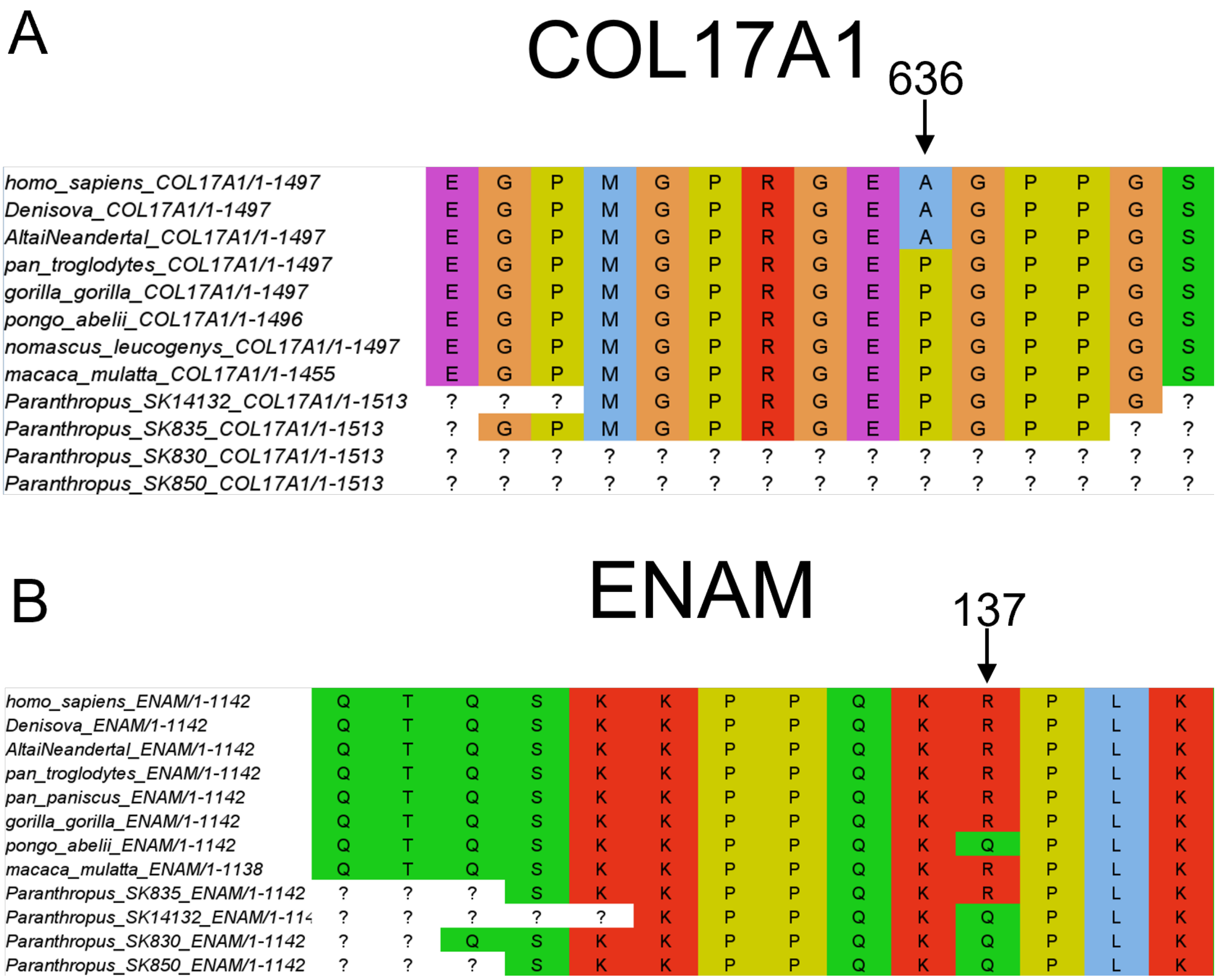


**Figure S15: Multiple sequence alignment of COL17A1 and ENAM.** (A) Multiple sequence alignment of COL17A1 using the 'Representative' dataset. The position of COL17A1-636 is displayed, covered in two out of the four samples, where *Paranthropus* shares the ancestral SAP for Great Apes. (B) Multiple sequence alignment of ENAM using the 'Representative' dataset. The position of ENAM-137 is displayed, covered in all four samples, where three *Paranthropus* have evidence of the Q allele and two for the R allele. Both alleles were represented in the spectra from sample SK14132 (possibly due to heterozygosity in that position) but this individual was represented just with a Q, due to the constraints of the fasta format and the higher number of recovered peptides for the Q allele.

ENAM-137 was covered in all four *Paranthropus* individuals. This position corresponds to the *Homo sapiens* Ensembl reference transcript ID: ENST00000396073.4, which is labelled as the canonical isoform of ENAM. As mentioned in the main text, that position was identified as variable between the four individuals:

In two individuals (SK830, SK850) only a glutamine (Q) was detected in that position, which differentiates from all African great apes, but not Pongo. The 'Diversity' dataset indicates that this particular site is fixed within modern Pongo species (sample of 27 individuals). This is further supported by the 'Independent' dataset, which also includes a *Gigantopithecus* sample. The *Gigantopithecus* sample also bears the Q SAP for that position. We conclude that this Q SAP on ENAM-137 is fixed in modern Pongos and likely in other related extinct Pongidae. In one individual (SK835), only an arginine (R) was identified in that position, which is shared with all other African great apes and not with

Pongo (Figure S15 B). Finally, one individual (SK14132) showed a strong signal for the presence of both SAPs, which was interpreted as a sign of heterozygosity for that site. This individual was represented with the Q SAP for the purpose of the phylogenetic analysis, since the fasta format does not allow for a diploid representation. Although the Q SAP matches that of the genus Pongo, our assessment is this is probably a result of the same mutation occurring twice in the Hominidae tree rather than incomplete lineage sorting or admixture (Extended Data Fig. 7).

Furthermore, a search of all reported present day human variation on those two sites (ENAM-137, COL17A1-636) was conducted. The search was able to identify a single SNP reported in present day humans, **rs773896528**, which can cause an R to Q substitution on ENAM-137. According to GnomAD however, this SNP only exists in East Asian populations with an extremely low frequency of 5.442e-05 (a single individual in the entire dataset). It is thus possible this reported mutation is caused by a statistical error. Even if this reported mutation is true, it is quite unlikely that the SNP is related to the *Paranthropus* variant, but rather due to a *de novo* mutation in contemporary humans. No SNP in present day humans that can cause an A to P substitution in COL17A1 was identified.

Finally a single substitution that is exclusively shared between one *Paranthropus* and all members of the *Homo* clade was also observed. This site, ALB-183, is particularly interesting since there are four possible alternative SAPs within present day hominids (Figure S16). Each different substitution on that site is unique to a hominid genus, making this site extremely informative. Unfortunately this position is covered in only one out of the four *Paranthropus* samples.

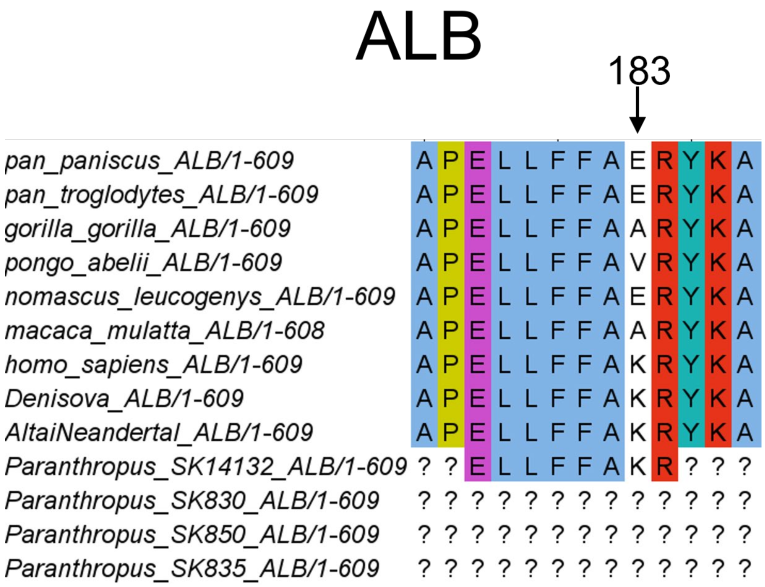


**Figure 16: Multiple sequence alignment of ALB using the 'Representative' dataset.** The position of ALB-183 is displayed, covered in only one *Paranthropus* individual. The individual shares the same SAP with modern humans, Neanderthals and Denisovans, to the exclusion of all other hominids.

- **Comparisons with other Great Apes:**

A number of SAPs that differentiate the *Paranthropus* samples from other Great Apes (Hominidae) were identified. From a total of 17 informative sites, most of the SAPs only differentiate from the *Pongo* lineage. A small number of SAPs differentiate from the other three genera of

Hominidae. A full list of all Hominidae informative SAPs as well as which species they match/differentiate from is provided in section 9.5.1 - 'SAP validations' (Table S13).

- **Comparisons with other ancient samples:**

The *Paranthropus* samples were also compared with the previously published palaeoproteomic data of Welker et al. 2020[46]. The *Homo erectus* individual from that study, also displays the R SAP on ENAM-137. Comparison with *Homo antecessor* was not possible, due to that site not being covered. Comparisons for the site of COL17A1-636 or ALB-183 were also not possible due to the same reason. Future comparisons with other palaeoproteomic samples will be limited to not only proteins, but the sites that are covered in these proteins. Although we cannot draw any conclusions, this is a hint that resolving the genetic relations of these archaic Hominidae will be more challenging than previously believed.

##### 10.2.3 Pairwise distances between *Paranthropus* and modern samples.

###### 10.2.3.1 **Pairwise distance matrix**

The pairwise distance matrix (Figure S20) shows relatively small values for all members of Hominidae but especially small for the African branch of the great apes. The lowest values between the *Paranthropus* individuals and any other great apes are between them and *Homo sapiens* or the Altai Neanderthal (equal distance), followed closely by the Denisovan. This is due to the Denisovan individual having additional unique SAPs compared to humans and Neanderthals, leading to slightly higher distance values.

| | **Altai Neanderthal** | **Denisova** | **SK14132** | **SK830** | **SK835** | **SK850** | ***G.gorilla*** | ***H.sapiens*** | ***P.paniscus*** | ***P.troglodytes*** | ***P.abelii*** |
|---|---|---|---|---|---|---|---|---|---|---|---|
| **Altai Neanderthal** | 0 | 0.000909 | 0.002763 | 0.001931 | 0.001794 | 0.001454 | 0.011111 | 0.000757 | 0.01031 | 0.010343 | 0.02282 |
| **Denisova** | 0.000909 | 0 | 0.004149 | 0.003868 | 0.003592 | 0.002911 | 0.011266 | 0.00106 | 0.010701 | 0.010649 | 0.02298 |
| **SK14132** | 0.002763 | 0.004149 | 0 | 0.00001 | 0.002062 | 0.00001 | 0.005539 | 0.002763 | 0.008803 | 0.009705 | 0.012511 |
| **SK830** | 0.001931 | 0.003868 | 0.00001 | 0 | 0.002372 | 0.00001 | 0.007741 | 0.001931 | 0.010195 | 0.009688 | 0.0136 |
| **SK835** | 0.001794 | 0.003592 | 0.002062 | 0.002372 | 0 | 0.00207 | 0.005392 | 0.001794 | 0.007554 | 0.009002 | 0.019949 |
| **SK850** | 0.001454 | 0.002911 | 0.00001 | 0.00001 | 0.00207 | 0 | 0.004366 | 0.001454 | 0.00758 | 0.008755 | 0.016119 |
| ***G.gorilla*** | 0.011111 | 0.011266 | 0.005539 | 0.007741 | 0.005392 | 0.004366 | 0 | 0.011265 | 0.01188 | 0.011574 | 0.021895 |
| ***H.sapiens*** | 0.000757 | 0.00106 | 0.002763 | 0.001931 | 0.001794 | 0.001454 | 0.011265 | 0 | 0.010506 | 0.010495 | 0.022977 |
| ***P.paniscus*** | 0.01031 | 0.010701 | 0.008803 | 0.010195 | 0.007554 | 0.00758 | 0.01188 | 0.010506 | 0 | 0.004076 | 0.028057 |
| ***P.troglodytes*** | 0.010343 | 0.010649 | 0.009705 | 0.009688 | 0.009002 | 0.008755 | 0.011574 | 0.010495 | 0.004076 | 0 | 0.024203 |
| ***P.abelii*** | 0.02282 | 0.02298 | 0.012511 | 0.0136 | 0.019949 | 0.016119 | 0.021895 | 0.022977 | 0.028057 | 0.024203 | 0 |

**Figure S20: The Pairwise Distance Matrix of the 'Representative' dataset.** The matrix was generated through PHYLIP's protdist and using the 'Representative' dataset as input.

#### 10.3. Phylogenetic inference Results

The aligned and corrected sequences were used to generate multiple phylogenetic trees using the different datasets, genes, software and sample compositions.

##### 10.3.1 ‘Representative’ Dataset

The first set of trees are the ‘concatenated‘ trees for the ‘Representative’ dataset (Figures S18, S19, S20). These 3 trees were generated using 3 different software and approaches but the same input dataset. The trees agree on the topological placement of the 4 *Paranthropus robustus* individuals but vary in their support. The MrBayes tree presents high supports for the topology, while the 2 maximum likelihood trees from PhyML and IQTree showcase much lower supports.

The Beast2 - tree (Figure S21) also supports a similar topology but in a bifurcating format. The placement of 3 *Paranthropus* (SK14132,SK830,SK850) as a single clade that is an outgroup to the genus *Homo* is confident, but the placement of SK835 as separate to either the other 3 *Paranthropus* individuals or the *Homo* clade is not.

###### 10.3.1.1 Bayesian Species Tree - MrBayes

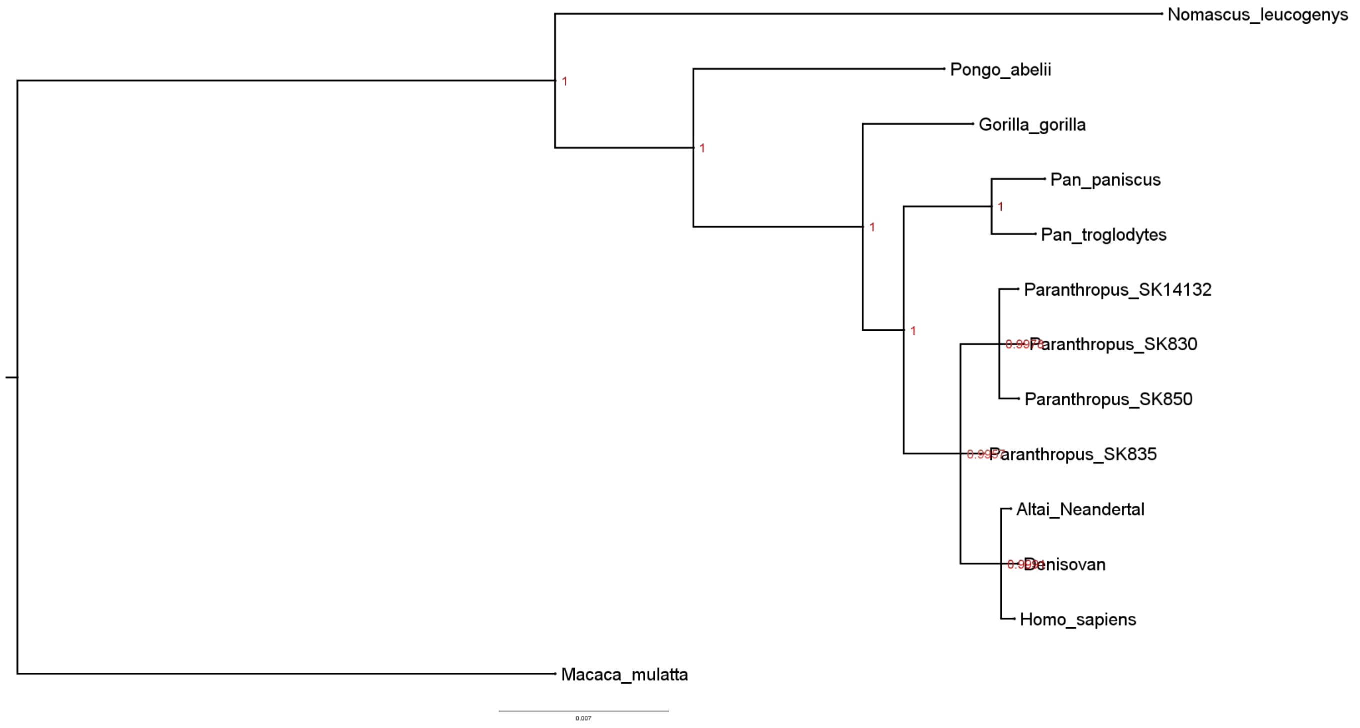


**Figure S18: Bayesian species tree generated using MrBayes and the ‘Representative’ dataset.**

##### 10.3.1.2 Maximum Likelihood Species Tree - PhyML

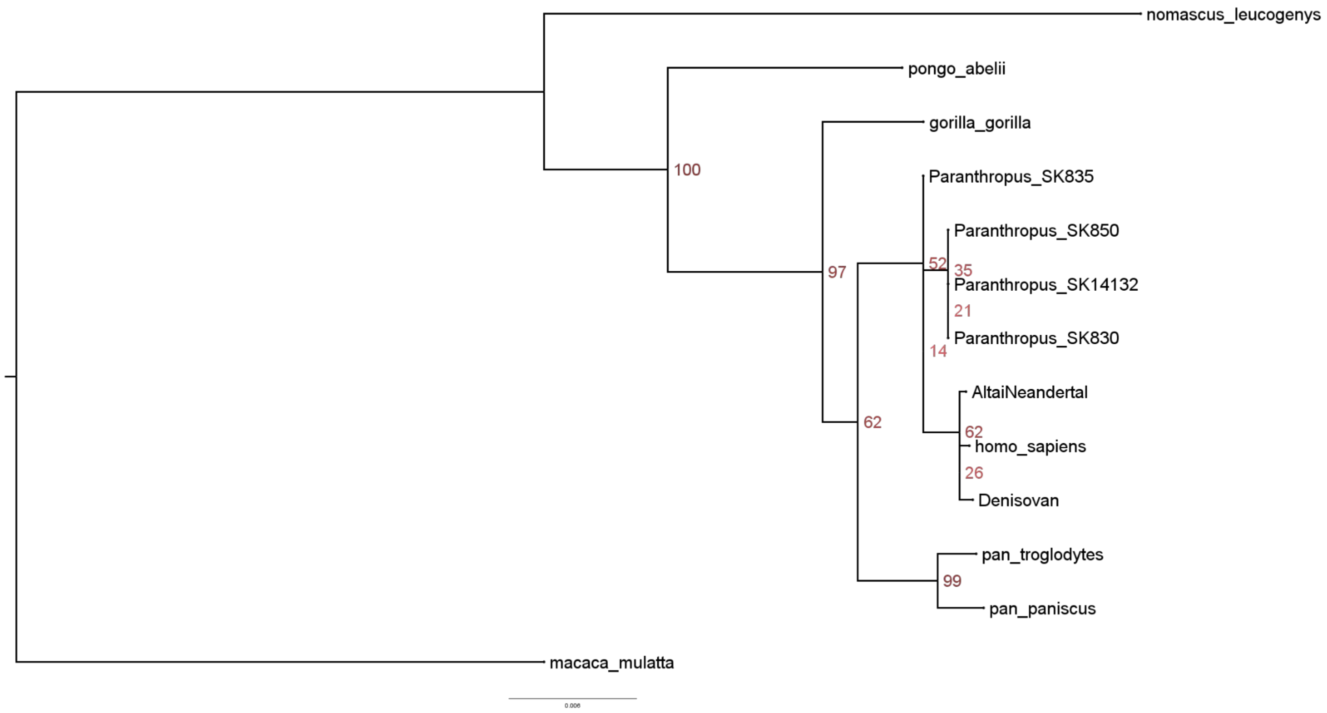


**Figure S19: Maximum Likelihood species tree generated using PhyML and the 'Representative' dataset .**

10.3.1.3 Maximum Likelihood Species Tree - IQ tree

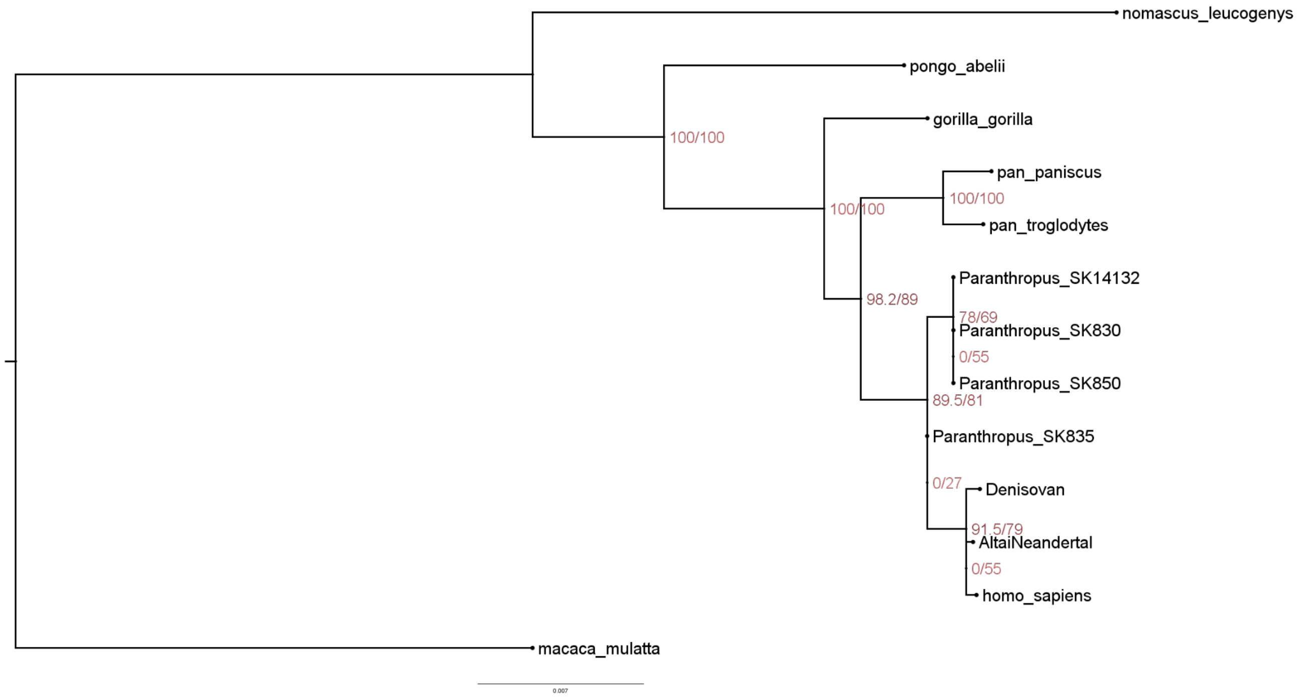


**Figure S20: Maximum likelihood species tree generated using IQTree and the ‘Representative dataset’.**

#### 10.3.1.4 Beast2 - species tree

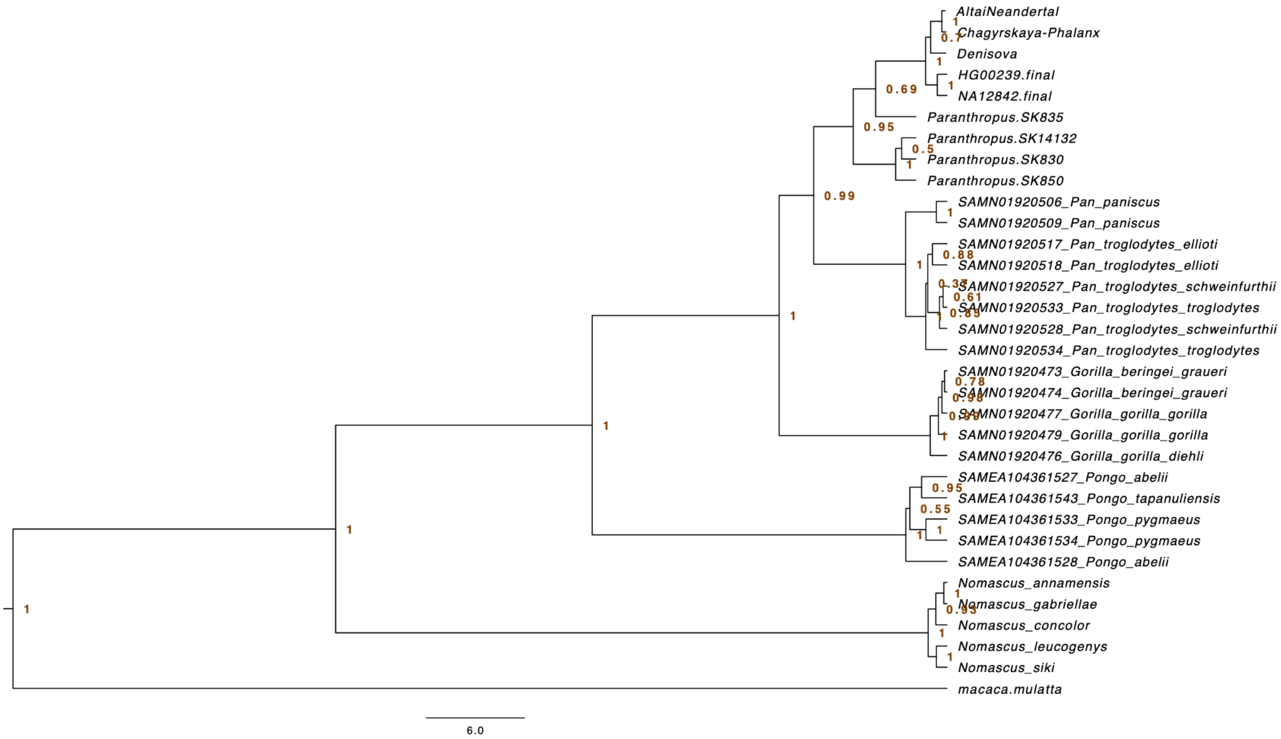


**Figure S21: Beast2 species tree generated from the 'Representative' dataset with a few additional individuals.**

##### 10.3.2 'Diversity' Dataset

The second set of trees are the concatenated trees for the 'Diversity' dataset (Figures S22,S23,S24). Neither the bayesian (Figure S22) nor the maximum likelihood trees (Figure S23) could place the *Paranthropus* samples within a clade with high confidence. The bayesian tree places all *Paranthropus* samples within the same clade along with modern humans, Neanderthals and Denisovans. It's possible that in the presence of a large dataset with relative within-species diversity, exemplified here with the large number of modern human samples, the few SAPs differentiating between *Paranthropus* and other species lose their importance.

The maximum likelihood tree places *Paranthropus* in the same manner as the smaller 'Representative' dataset trees, but again with low bootstrap values. The small number of informative sites that was recovered for the *Paranthropus* samples, along with inconsistencies between the gene trees, are likely the source of the low confidence bootstrap values.

When the samples were all assigned to a specific 'taxon' through Starbeast3's options, the multispecies coalescent placed the *Paranthropus* samples as a sister group to the *Homo* clade (Figure S24). Although the tree only shows a small number of tips (a single tip per taxon), compared to the 'Representative' trees, these *taxon*-tips are composed of genetic information from dozens (e.g. *Pan paniscus* tip) or even hundreds of individuals (e.g. *Homo sapiens* tip). Although these results agree with the 'Representative' dataset trees and show high support, they are generated using strong priors (manually assigning each sample to a taxon).

##### 10.3.2.1 Bayesian Tree - MrBayes

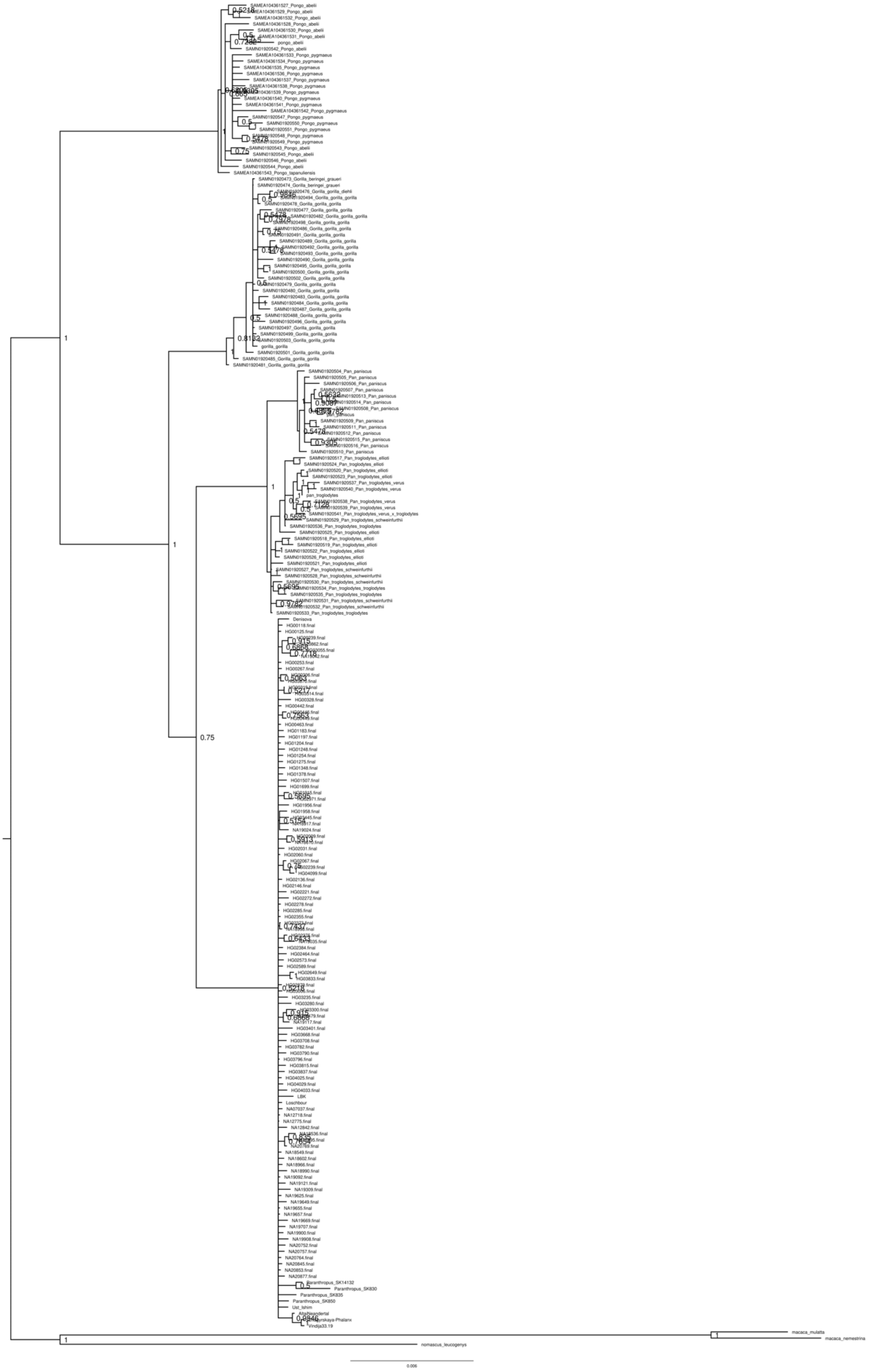

**Figure S22: Bayesian species tree generated using MrBayes and the 'Diversity' dataset.**

10.3.2.2 Maximum Likelihood Tree - PhyML

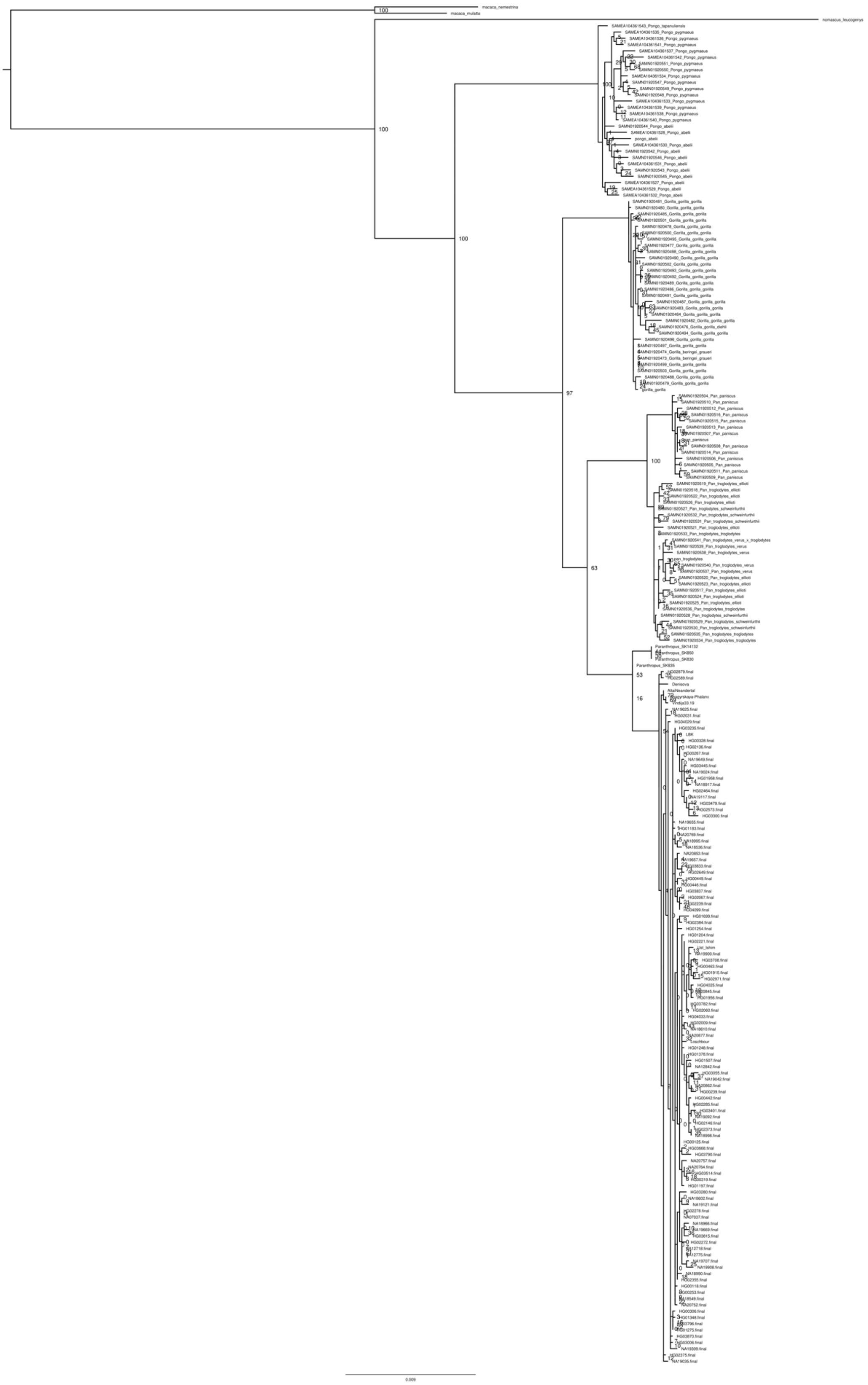

**Figure S23: Maximum Likelihood species tree generated using PhyML and the 'Diversity' dataset.**

##### 10.3.2.3 StarBeast3 - Multispecies Coalescent Tree

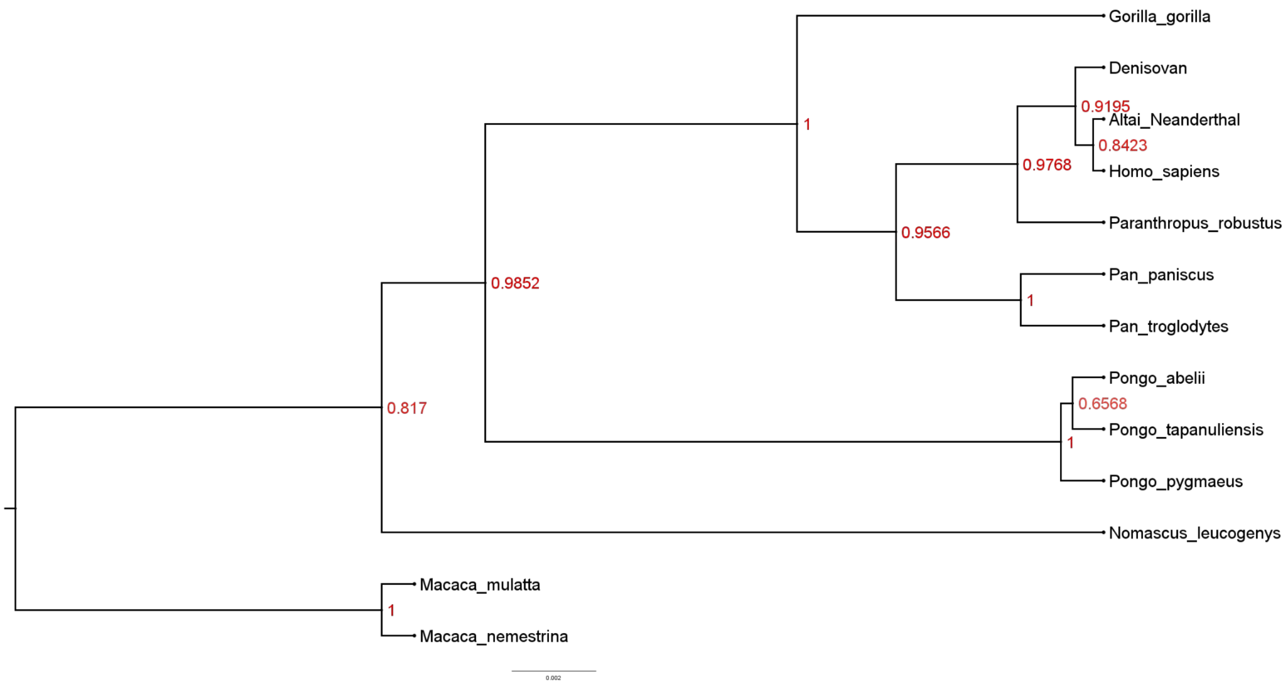


**Figure S24: Multispecies Coalescent Tree generated using Starbeast3 and the 'Diversity' dataset.** All samples were all assigned to one of the 13 taxons which are visible as tips in the tree.

##### 10.3.3 'Independent' Dataset

The third set of trees are the results using the 'Independent' dataset. For the maximum likelihood tree, separation of the genera is in accordance with the cannon in *Hominidae* (Figures S25-28). Both the maximum likelihood (Figure S25) and the Bayesian tree (Figure S27) support the same placement of the 4 *Paranthropus* individuals, as the trees generated by the 'Representative' dataset. Three of the *Paranthropus* individuals (SK830, SK850, SK14132) group together, to the exclusion of the 4th individual (SK835). With the exception of *Pan*, individuals of different species within the same genus do not necessarily form a monophyletic clade, including the species of *Homo*.

A reduction of the whole reference dataset still yields a similar topology with correct separation of the genera as in the dataset with the full length sequences (Figure S26, Figure S28). Both for the maximum likelihood and the Bayesian reduced tree, the results resemble the trees of the non-reduced dataset. The reduction of the dataset still yields a similar topology with correct separation of the genera as in the dataset with the full length sequences, with the exception of the Bayesian-reduced tree no longer separating the two species of *Pan*.

Note that in particular if data is fragmentary, such as in the case of *Homo erectus* from Dmanisi, the individual is placed with low confidence both in the maximum likelihood and the Bayesian tree. In cases like these, the placement of the sample might also be in unexpected positions on the tree, as demonstrated by that *Homo erectus* sample in the Bayesian tree (Figure S27). This is almost certainly due to the low quality and coverage of the sample, as pointed out by authors that study [46] rather than a real phylogenetic signal. We advise caution when interpreting trees that include samples from different palaeoproteomic studies, as the same informative sites need to be recovered for meaningful comparisons between them.

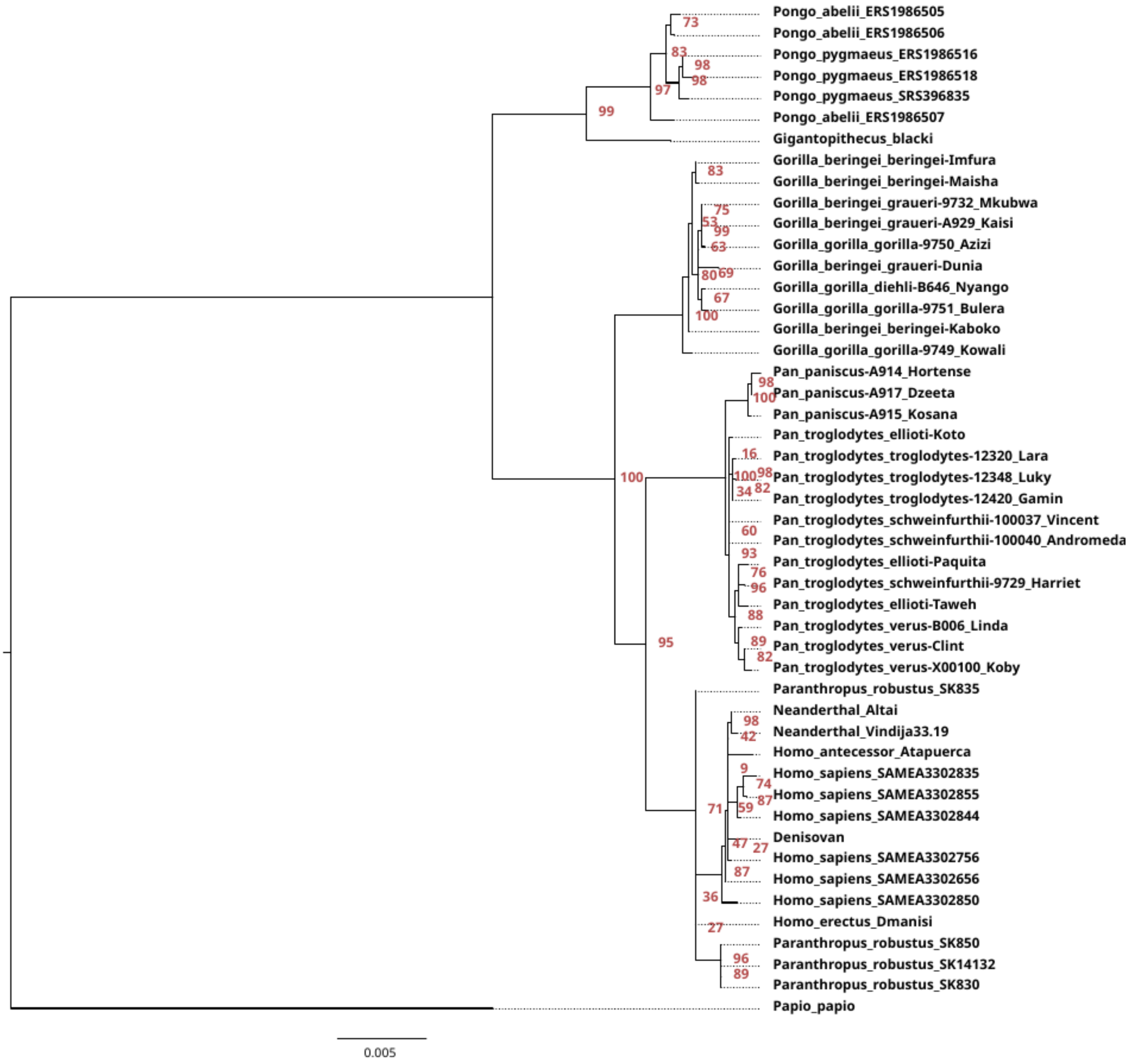


**Figure S25 : Maximum Likelihood tree generated using IQtree v1.6.12 and the 'Independent' dataset.**

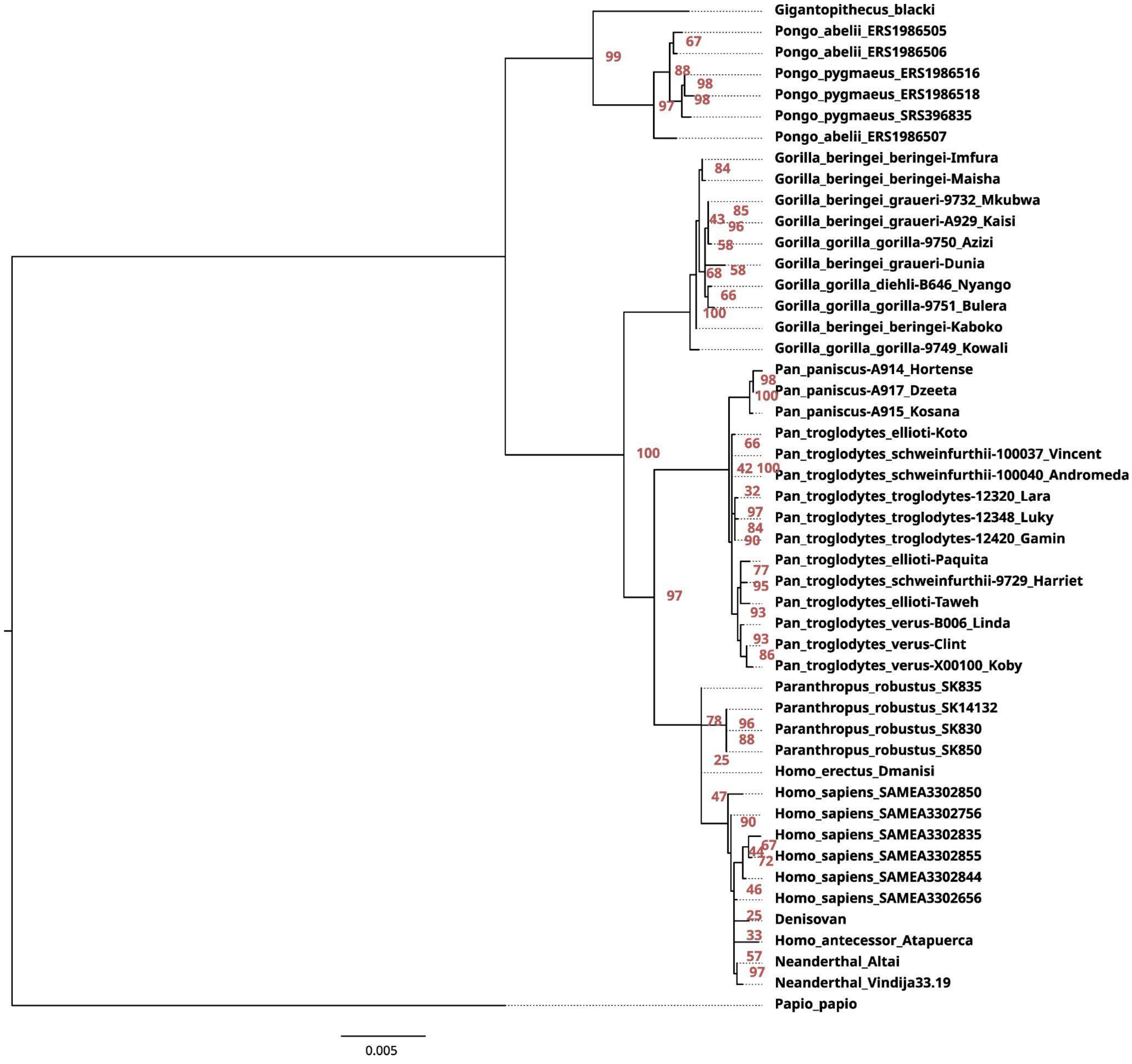


**Figure S26: Maximum Likelihood tree generated using IQtree v1.6.12 and the 'Independent' dataset, reduced to amino acid positions for which at least one ancient sample was available.**

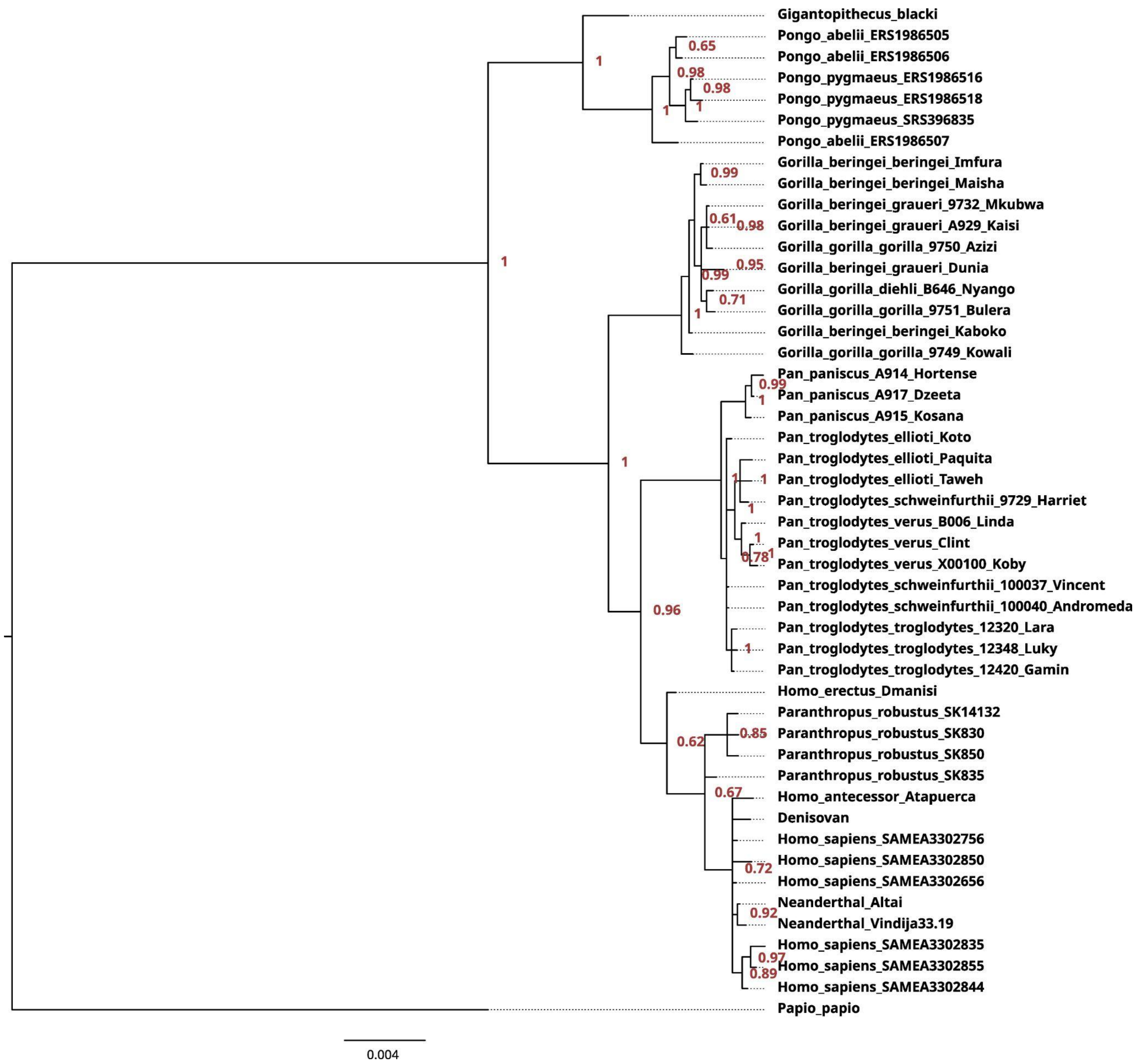


**Figure S27: Bayesian tree generated using MrBayes v3.2.7a and the ‘Independent’ dataset.**

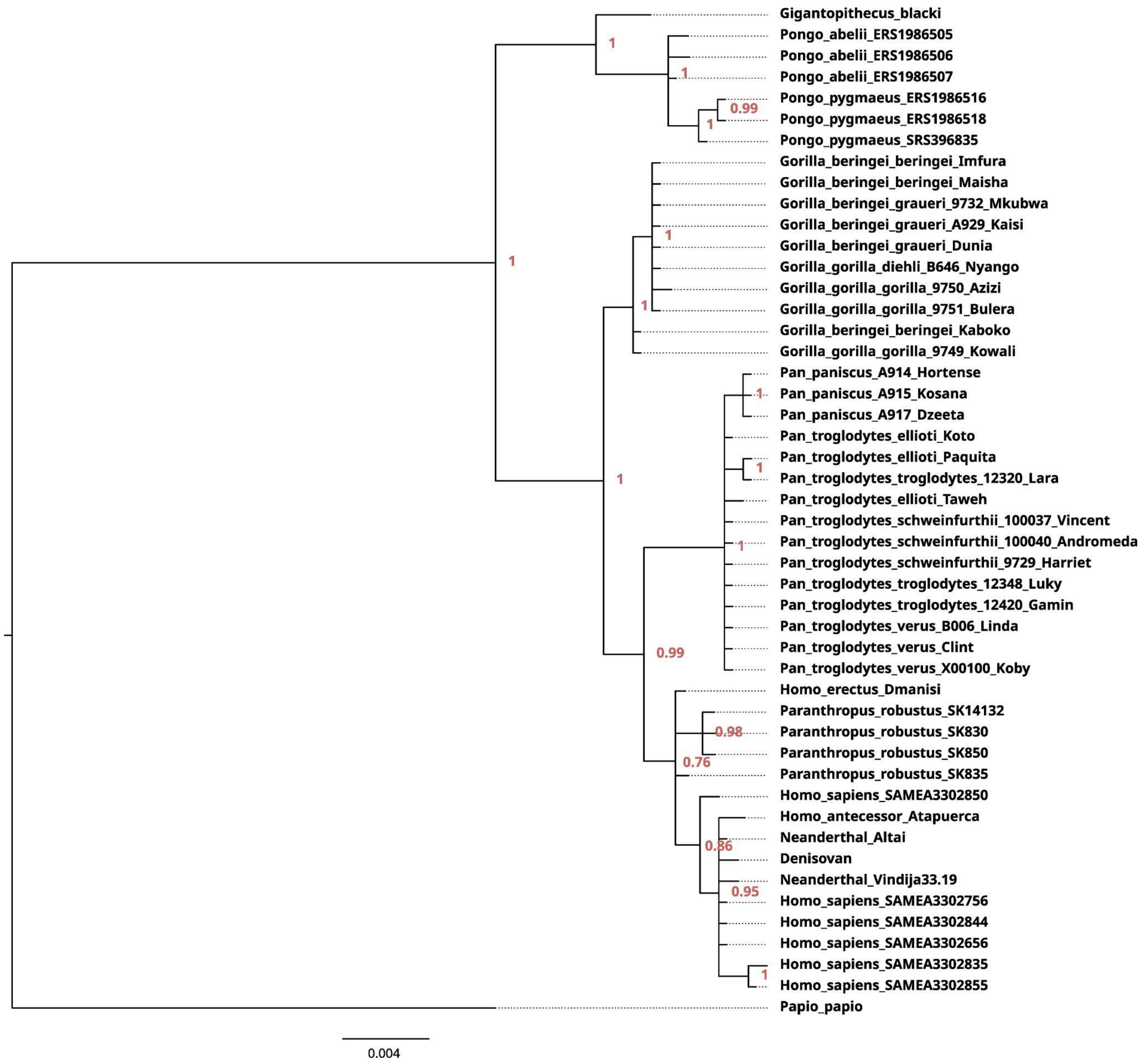


**Figure S28: Bayesian tree generated using MrBayes v3.2.7a and the 'Independent' dataset, reduced to amino acid positions for which at least one ancient sample was available.**

#### 10.4. Genetic Variation Metrics Results

Our workflow identified **29 SNPs** in present day humans that have a missense effect, and fall within the regions that were recovered by the 4 *Paranthropus* individuals. Only 2 of these SNPs have a frequency greater than 1%: **rs7439186** has a frequency of 0.111 in our dataset and **rs113506649** has a frequency of 0.014. All 29 of these sites/SNPs, regardless of frequency, were included in the analysis.

The genetic diversity metrics of the final, filtered VCF file, are as follows:

- Expected Heterozygosity: 0.252
- Observed Heterozygosity: 0.238
- Watterson's Estimator: 6.264
- Total length of corresponding nucleotide sequence: 1272

The resulting sampling measurements from the workflow are presented below:

1) Frequency of times at least one alternative amino acid ***allele*** in any of the sites was sampled: 0.713
2) Number of sites (out of 29 possible) that have at least one alternative amino acid allele identified in a run: (Figure S29 - average is 0.871):
3) Frequency of times at least two alternative amino acid alleles in any of the sites were sampled: 0.223
4) Frequency of times at least one **homozygous** individual for the alternative amino acid allele in any of the sites was sampled: 0.062

The results suggest that - under a scenario where the Paranthropus sample comes from a single population or group with similar levels of heterozygosity as present-day humans - it would not be unexpected to find within-sample genetic variation at the amino acid level (within the observable protein region) as large as what is actually observed in Paranthropus. It should be noted however that although sampling single alternative **alleles** for a variant site is frequent, sampling homozygous individuals for an alternative allele is much less unlikely.

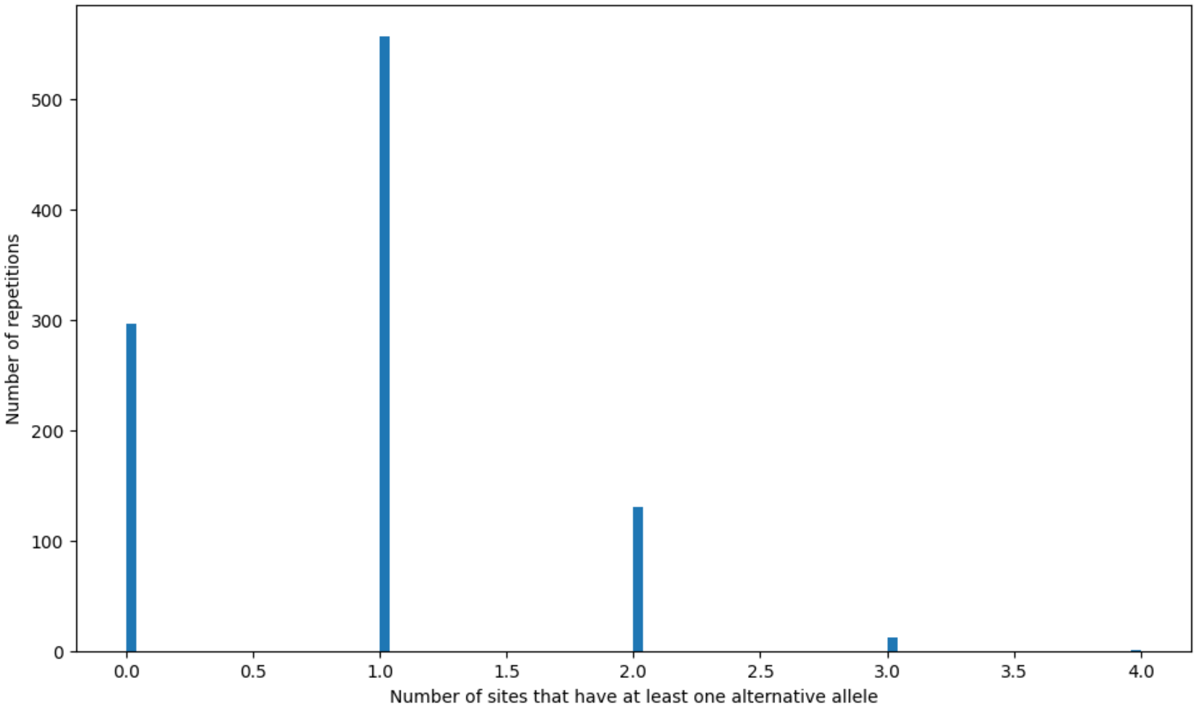


**Figure S25: Histogram showing the number of sites with at least one alternative allele (x axis) and for how many repetitions (y axis).**

### 11. Morphological analyses

#### 11.1. Morphological analysisfor sex assignment

Buccolingual and mesiodistal measurements of ndividuals that have previously been sexed in the literature are identified in Supplementary Document 3, and in the resultant boxplots and bivariate plots (Extended Data Fig. 10, Extended data Fig. 9). The boxplots of P4 dimensions of *P. robustus* indicate that SK 830, a mandibular P4 identified as female in this study, is in the lower half of the size distribution in both MD and BL diameters (Extended Data Fig. 10), and plots close to other previously identified female specimens in the bivariate plot (Extended Data Fig. 9). The P3 MD boxplot highlights the difference in size between two previously identified *P. robustus* females, SK 96 and DNH 7, as well as the large size range of previously identified males. SK 850, a mandibular P3 identified as male in this study, falls within the lower distribution of this boxplot (at the position of the first quartile). The boxplots of M3 dimensions also show a wide range of size variability across previously identified male individuals. SK 835, a maxillary M3 identified as male in this study, but previously classified as female (Dean et al., 2020), falls in the lower half of the distribution in its MD dimension, and at the centre of the BL distribution (Extended Data Fig. 10). SK 835 plots centrally in the bivariate plot (Extended Data Fig. 9).

#### 11.2. Geometric morphometric analysis

We conducted geometric morphometric analyses of the enamel-dentine junction of the $M^3$ SK 835 and of the $P_4$ SK 830 (Figure S26), and compared them with those of southern African *Australopithecus* and *Paranthropus* specimens, as well as African and Asian early *Homo* teeth (Extended Data Fig. 8).

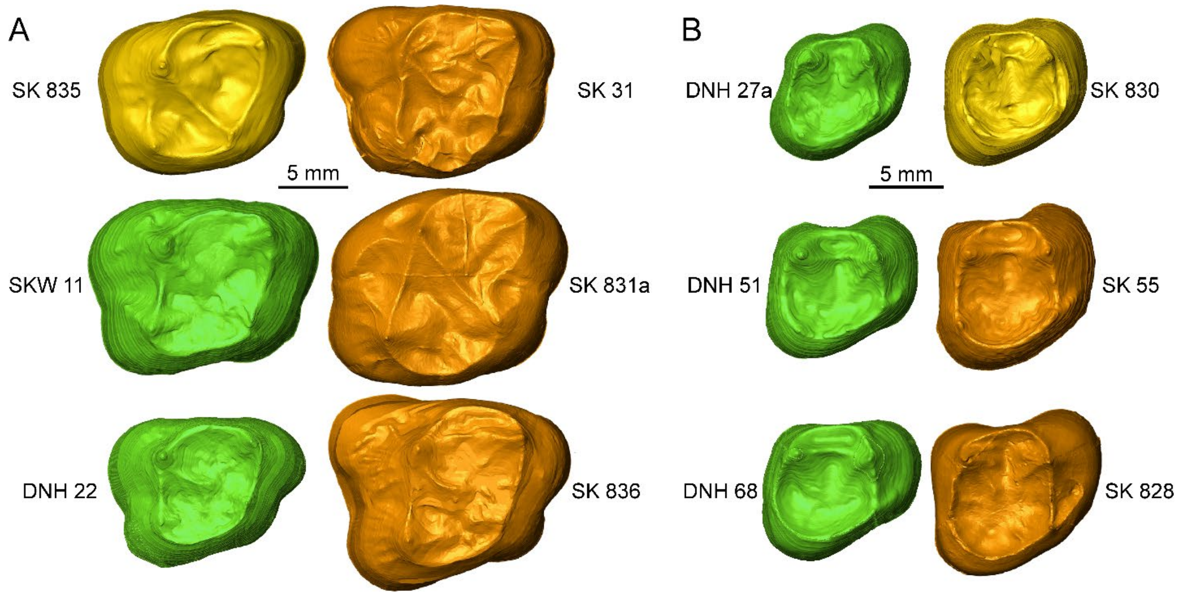


**Figure S26:** The EDJ of the $M^3$ SK 835 (A) and $P^4$ SK 830 (B) from Swartkrans compared with some of the *Paranthropus* specimens from Drimolen (in green; including also the specimen SKW 11 from Swartkrans) and Swartkrans (in orange) used in the geometric morphometric analyses

Results of these analyses discriminate well between the three genera (Extended Data Fig. 8; Table S20) and unambiguously show that SK 835 and SK 830 belong to *Paranthropus* and significantly differ from *Australopithecus* and *early Homo* (Extended Data Fig. 8; Table S21). When only southern African *Paranthropus* specimens are included in the geometric morphometric analyses, two groups can be distinguished, one including mostly specimens from Swartkrans and Kromdraai (except the $M^3$ SKW 11, that differs from most Swartkrans teeth and is more similar to the Drimolen specimens; Figure. S3), and another one constituted by material from Drimolen (Table S22). The shape of the EDJ of the $M^3$ SK 835 is statistically more similar to that of the Drimolen specimens than to those of the Swartkrans and Kromdraai assemblage, that includes the holotype of *P. robustus* (TM 1517), while that of the $P_4$ SK 830 resembles more the latter group and statistically differs from the material from Drimolen (Fig. 4; Table S23).

**Table S20**: Cross-validated CVA results of correctly classified individuals using shape data and *Australopithecus* (AUS), Early and Middle Pleistocene *Homo* (HOM), and *Paranthropus* (PAR) used as a priori groups.

| Tooth | HOM | AUS | PAR | overall accuracy |
|---|---|---|---|---|
| **$M^3$** | 5/6 | 10/11 | 11/11 | 92.9% |
| **$P_4$** | 9/9 | 14/16 | 12/13 | 92.1% |

**Table S21.** Typicality probabilities of the investigated specimens computed for the CVA of shape analyses. For each specimen, typicality probabilities are shown only for the group to which that specimen was affiliated. AUST: *Australopithecus*; HOM: Early and Middle Pleistocene *Homo*; PAR: *Paranthropus*.

| Tooth | Specimen | HOM | AUS | PAR |
|---|---|---|---|---|
| $M^3$ | SK 835 | - | - | 0.38 |
| $P_4$ | SK 830 | - | - | 0.96 |

**Table S22:** Cross-validated CVA results of correctly classified *Paranthropus* individuals using shape data and two a priori groups, one including material from Swartkrans and Kromdraai (SK), and the other one based on specimens from Drimolen (D). For the $M^3$, the specimen SKW 11 was included in group D (see details in Supplementary Information 7).

| Tooth | SK | D | overall accuracy |
|---|---|---|---|
| **$M^3$** | 7/8 | 4/4 | 91.7% |
| **$P_4$** | 9/10 | 3/3 | 92.3% |

**Table S23:** Typicality probabilities of the investigated *Paranthropus* specimens computed for the CVA of shape analyses. For each specimen, typicality probabilities are shown only for the group to which that specimen was affiliated. SK: Swartkrans and Drimolen; D: Drimolen.

| Tooth | Specimen | SK | D |
|---|---|---|---|
| $M^3$ | SK 835 | - | 0.05 |
| $P_4$ | SK 830 | 0.05 | - |

### DATA DEPOSITION NOTE

The mass spectrometry proteomics data have been deposited to the ProteomeXchange Consortium (http://proteomecentral.proteomexchange.org) via the PRIDE partner repository [137] with the dataset identifier **PXD040221.**
