## Supplementary Document 1 for "Enamel proteins reveal biological sex and genetic variability within southern African *Paranthropus*"

Variable sites detected: Comparison of one experimental spectra (top) with its predicted spectra (bottom).



### AHSG - 117

F Q L L K L

Precursor m/z: 381.7416 Charge: +2 Fragmented Bonds: 5/5

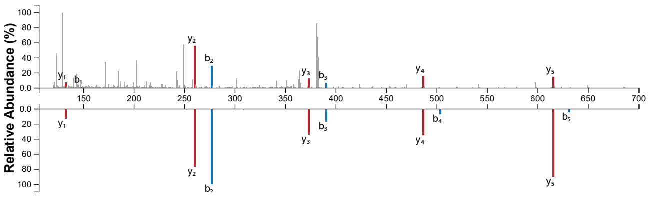


Precursor m/z: 381.7416 Charge: +2 Fragmented Bonds: 5/5

F E L L K L

### ALB - 574

I K K Q T A L V E L V K

Precursor m/z: 457.6254 Charge: +3 Fragmented Bonds: 11/11

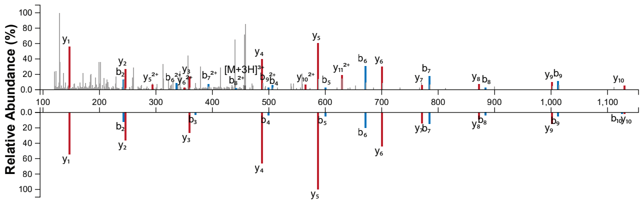


Precursor m/z: 457.6254 Charge: +3 Fragmented Bonds: 10/11

I K K E T A L V E L V K

### ALB - 202

E L L F F A K R

Precursor m/z: 503.2976 Charge: +2 Fragmented Bonds: 6/7

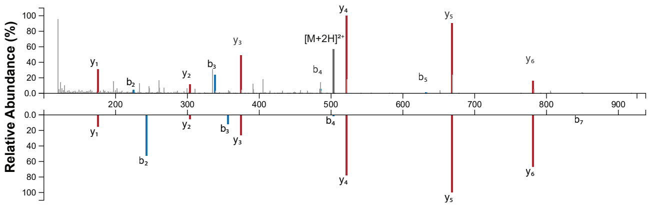


Precursor m/z: 512.3029 Charge: +2 Fragmented Bonds: 6/7

E L L F F A K R

### AMBN - 270

E E V A G G R E D P M

Precursor m/z: 603.2588 Charge: +2 Fragmented Bonds: 10/10

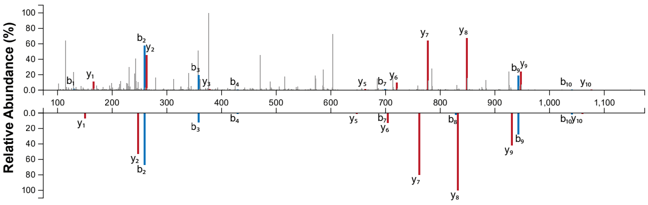


Precursor m/z: 595.2613 Charge: +2 Fragmented Bonds: 10/10

E E V A G G R E D P M

### AMELX - 127

T P I Q H H Q P N

Precursor m/z: 537.7462 Charge: +2 Fragmented Bonds: 8/8

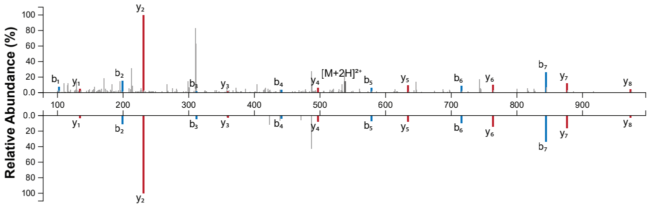


Precursor m/z: 537.7462 Charge: +2 Fragmented Bonds: 8/8

T P I E H H E P D

### AMELX - 135

P N L P P P A Q Q P

Precursor m/z: 531.2611 Charge: +2 Fragmented Bonds: 7/9

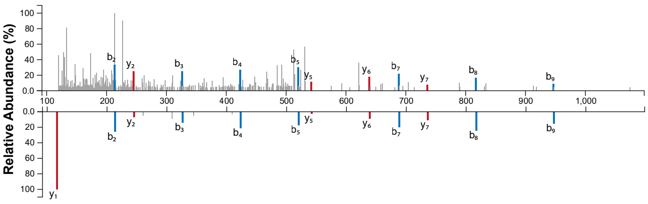


Precursor m/z: 531.2611 Charge: +2 Fragmented Bonds: 7/9

P D L P P P A E E P

### AMELY - 101/109

I P V V P A Q Q P R

Precursor m/z: 532.8030 Charge: +2 Fragmented Bonds: 9/9

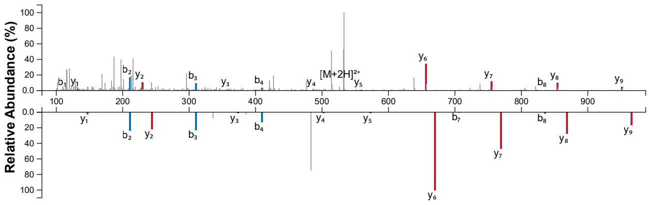


Precursor m/z: 539.8108 Charge: +2 Fragmented Bonds: 9/9

I P V V P A E E P K

### AMELY - 184

L P P I L P D L H L

Precursor m/z: 564.3448 Charge: +2 Fragmented Bonds: 9/9

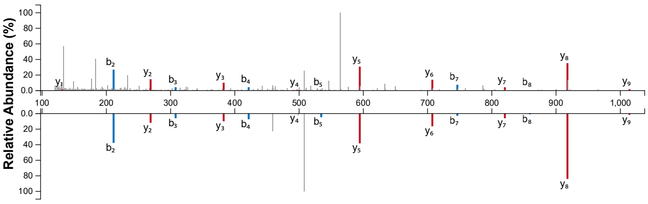


Precursor m/z: 564.3448 Charge: +2 Fragmented Bonds: 8/9

L P P I L P D L H L

Variable sites detected: Comparison of one experimental spectra (top) with its predicted spectra (bottom).



### AMTN - 99

V L P I F V T Q L

Precursor m/z: 515.8128 Charge: +2 Fragmented Bonds: 7/8

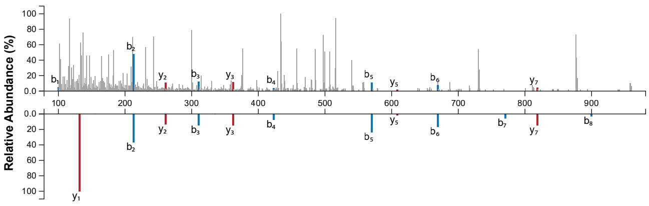


Precursor m/z: 515.8128 Charge: +2 Fragmented Bonds: 7/8

V L P I F V T E L

### COL17A1 - 636

G P M G P R G E P G P P G

Precursor m/z: 606.2717 Charge: +2 Fragmented Bonds: 6/12

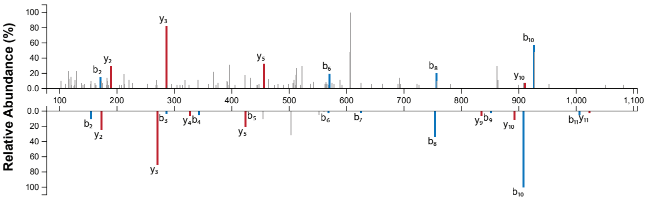


Precursor m/z: 589.2871 Charge: +2 Fragmented Bonds: 10/12

G P M G P K G E P G P P G

### ENAM - 147

P Q P E E E A Q P P Q A

Precursor m/z: 662.2830 Charge: +2 Fragmented Bonds: 9/11

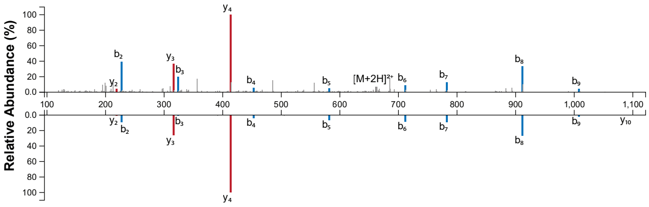


Precursor m/z: 662.2830 Charge: +2 Fragmented Bonds: 9/11

P E P E E E A E P P E A

### ENAM - 190

R P P I S N E E G

Precursor m/z: 500.2407 Charge: +2 Fragmented Bonds: 8/8

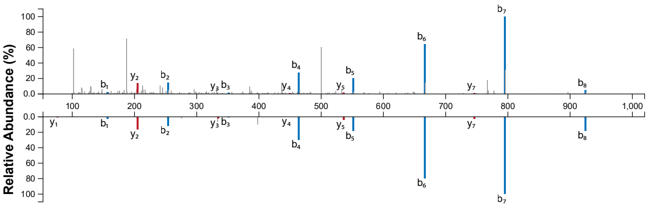


Precursor m/z: 500.2407 Charge: +2 Fragmented Bonds: 8/8

R P P I S D E E G

### ENAM - 212

F G G R P P Y Y

Precursor m/z: 478.7349 Charge: +2 Fragmented Bonds: 7/7

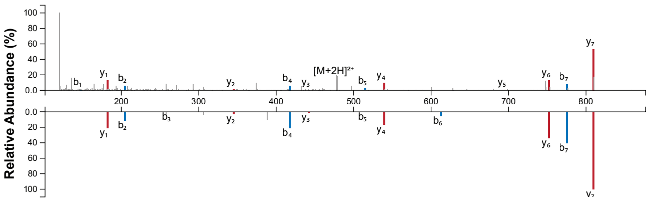


Precursor m/z: 478.7349 Charge: +2 Fragmented Bonds: 7/7

F G G R P P Y Y

### MMP20 - 292

K P S I P D L

Precursor m/z: 385.2263 Charge: +2 Fragmented Bonds: 6/6

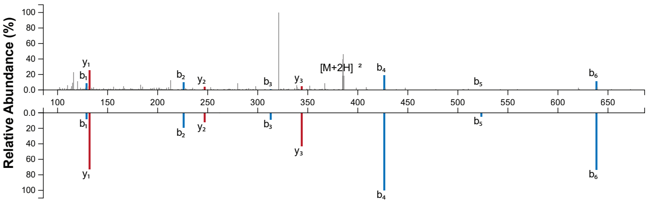


Precursor m/z: 385.2263 Charge: +2 Fragmented Bonds: 6/6

K P S I P D L

### ODAM - 271

D K T D S L R E P

Precursor m/z: 530.7671 Charge: +2 Fragmented Bonds: 8/8

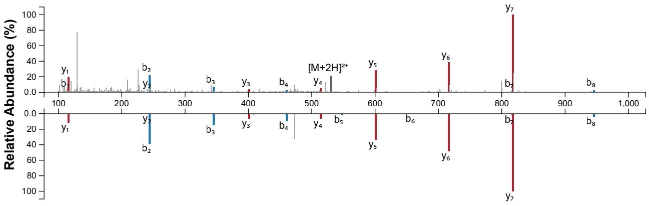


Precursor m/z: 530.7671 Charge: +2 Fragmented Bonds: 7/8

D K T D S L R E P
