## Supplementary figures and images for "Enamel proteins reveal biological sex and genetic variability within southern African *Paranthropus*"

### Supplementary Document 2

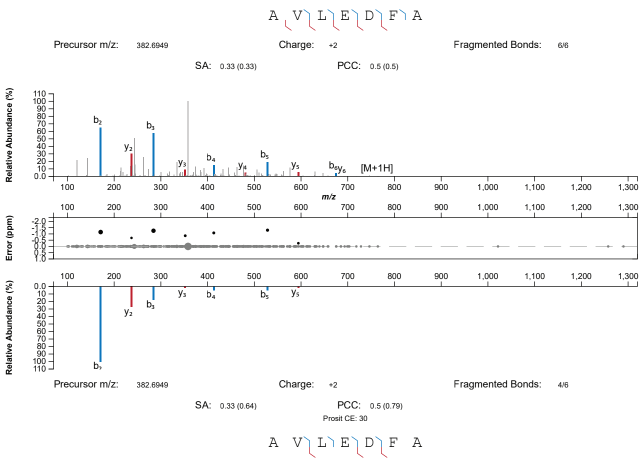

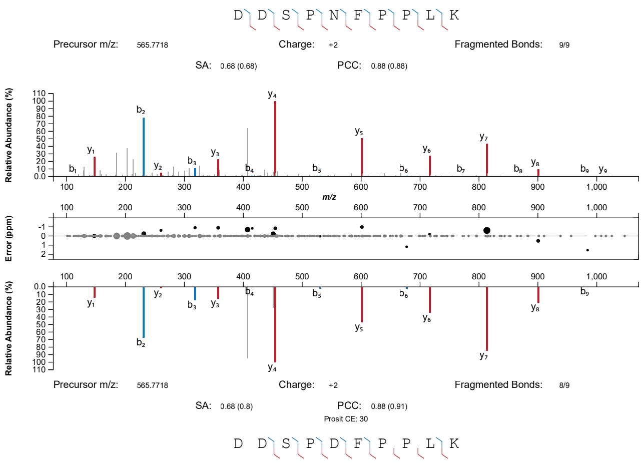

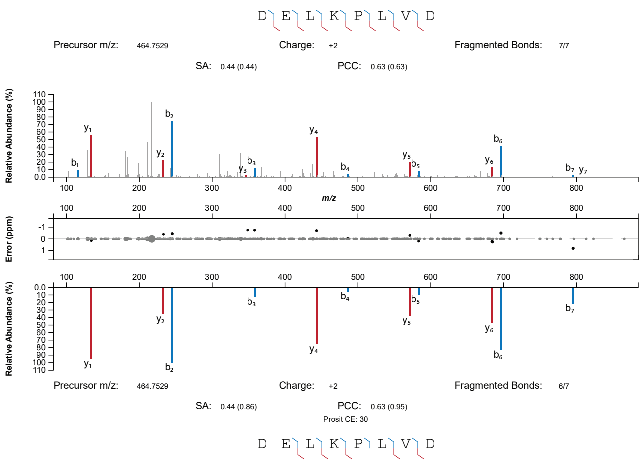

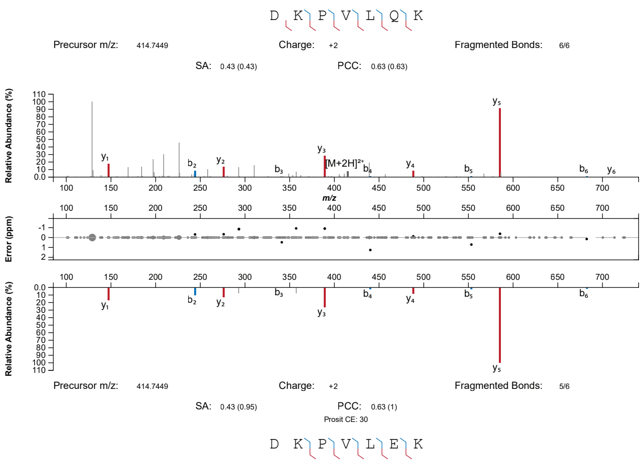

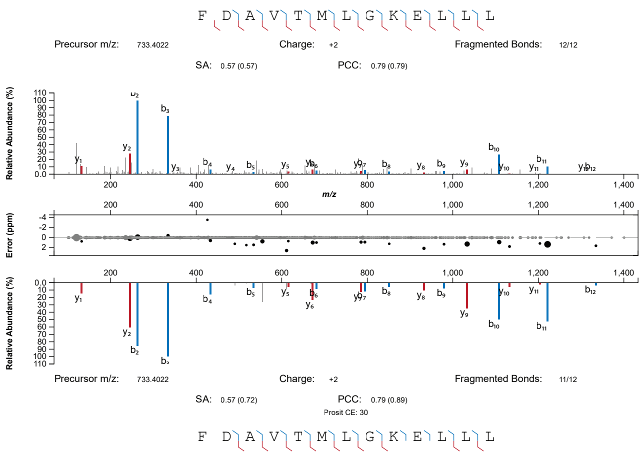

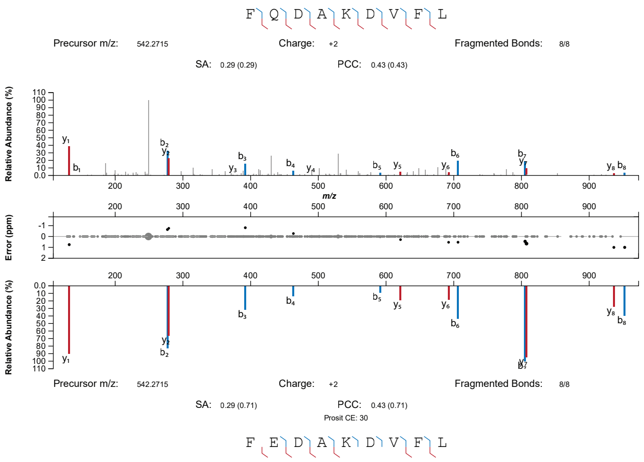

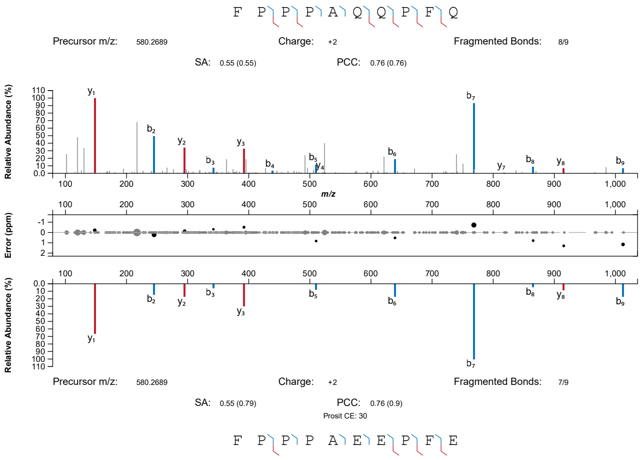

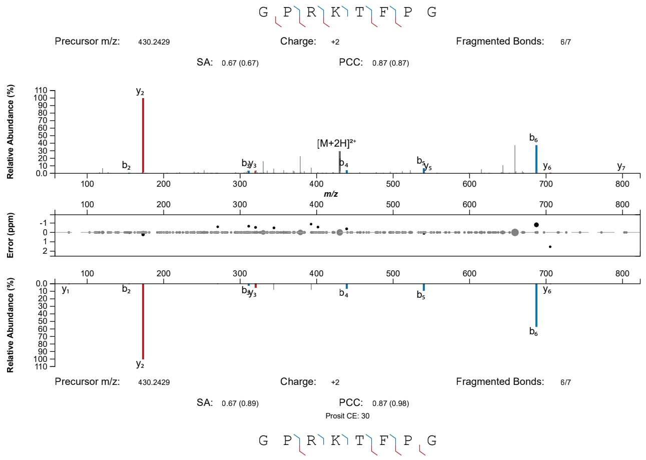

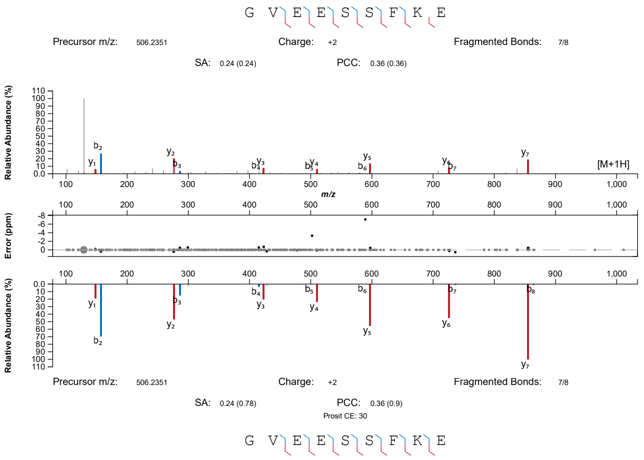

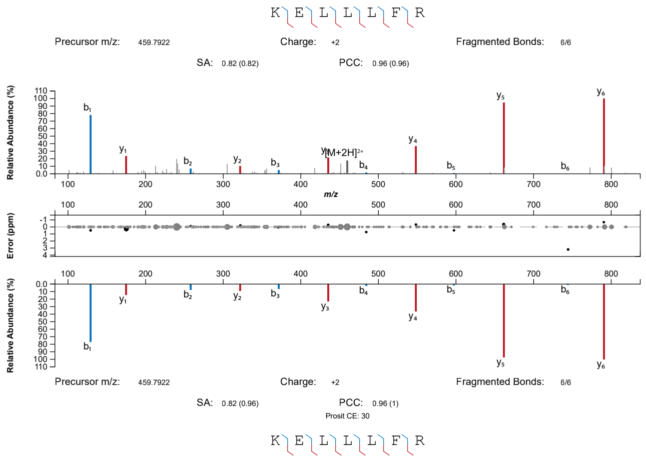

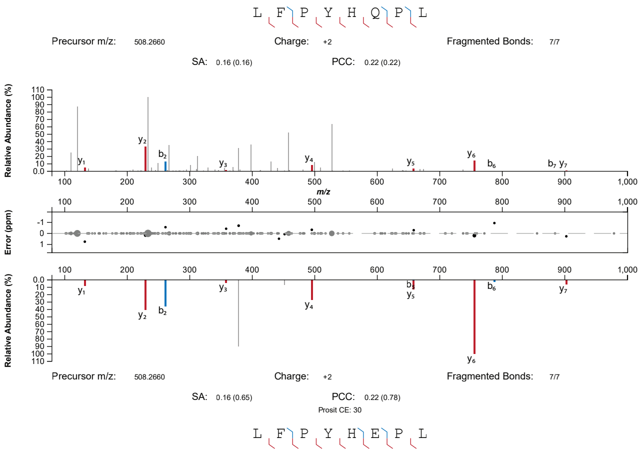

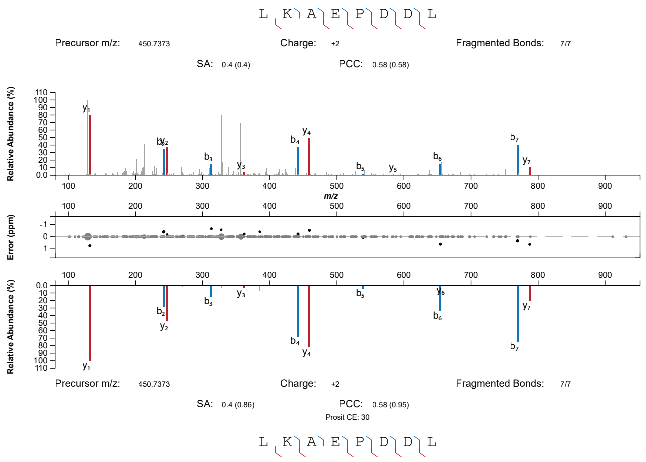

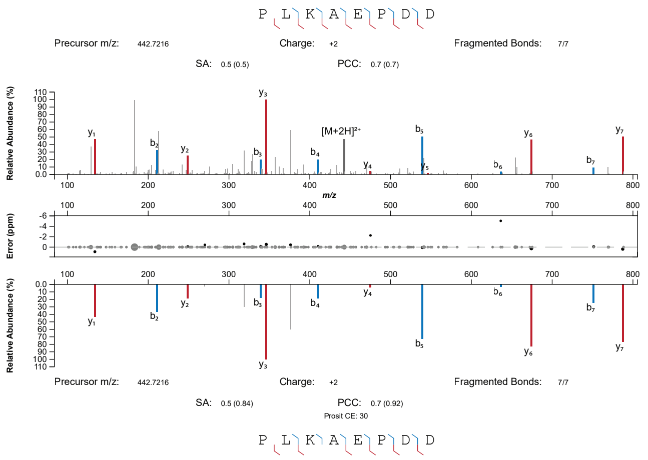

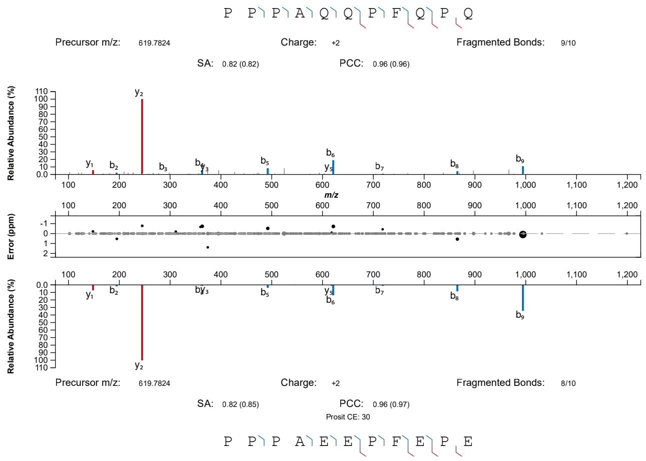

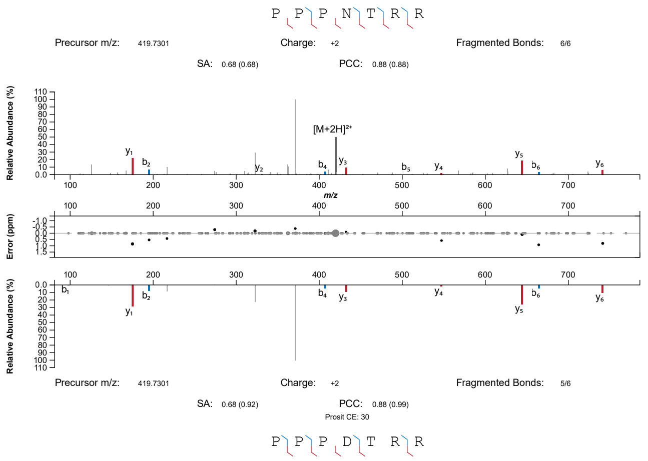

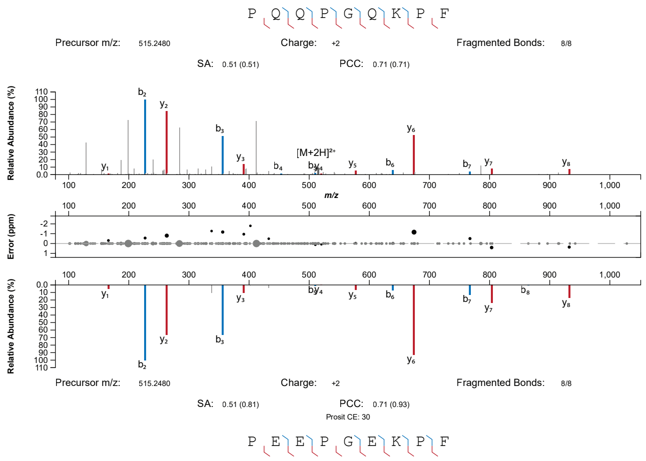

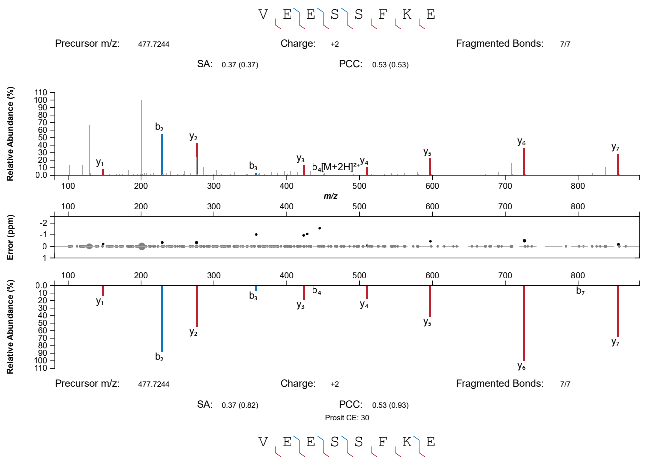

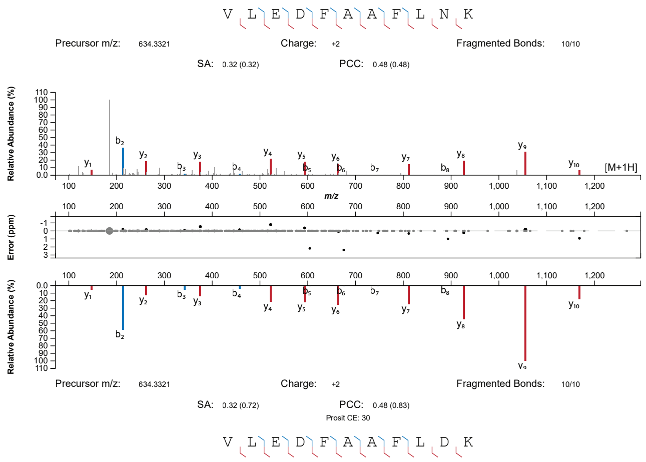
