## Supplementary Document 3 for "Enamel proteins reveal biological sex and genetic variability within southern African *Paranthropus*"

**Supplementary document 3: *Paranthropus robustus* standard dental measurements**

| Sample | Right Upper M3 | | Left Upper M3 | | Upper M3 side not specified | | Right Lower P3 | | Left Lower P3 | | Lower P3 side not specified | | Right Lower P4 | | Left Lower P4 | | Lower P4 side not specified | | Sex | Reference (sex) | Reference (measurements) |
|---|---|---|---|---|---|---|---|---|---|---|---|---|---|---|---|---|---|---|---|---|---|
| | MD | BL | MD | BL | MD | BL | MD | BL | MD | BL | MD | BL | MD | BL | MD | BL | MD | BL | | | |
| SK 6 | | | | | | | | | 10.4 | 12.5 | | | | | 11 | 12.3 | | | male | Grine et al. 2012/Thackeray et al. 2005 | Wood 1991 |
| SK 7 | | | | | | | | | | | | | 11 | 13.4 | | | | | | | Wood 1991 |
| SK 9 | | | | | | | | | | | | | | | 10.8 | 12.8 | | | | | Wood 1991 |
| SK 11 | | | 15.1 | 16.1 | | | | | | | | | | | | | | | male | Lockwood et al. 2007 | Robinson 1956 |
| SK 12 | | | | | | | | | | | | | 10.5 | 12.5 | | | | | male | Keyser 2000 | Robinson 1956 |
| SK 13 | 14.2 | 17.5 | 14.7 | 17.5 | | | | | | | | | | | | | | | | | Wood 1991 |
| SK 16 | | | | | 14.2 | 16.8 | | | | | | | | | | | | | | | Robinson 1956 |
| SK 18a | | | | | | | | | 8.9 | 10.1 | | | | | | | | | | | Wood 1991 |
| SK 23 | | | | | | | 9.6 | 11.7 | 9.6 | 11.5 | | | 11.1 | 13.7 | 11.1 | 14.4 | | | male | Schwartz and Tattersall 2005 | Wood 1991 |
| SK 25 | | | | | | | | | | | | | 11.4 | 13.3 | | | | | male | Schwartz and Tattersall 2005 | Robinson 1956 |
| SK 30 | | | | | | | | | 10.2 | 11.2 | | | | | | | | | | | Wood 1991 |
| SK 31 | 15.5 | 16.5 | | | | | | | | | | | | | | | | | | | Wood 1991 |
| SK 34 | | | | | | | 10.4 | 12.7 | | | | | | | 12.3 | 13.8 | | | male | Grine et al. 2012 | Wood 1991 |
| SK 36 | 15.1 | 17.2 | | | | | | | | | | | | | | | | | | | Wood 1991 |
| SK 41 | | | | | 15.1 | 16 | | | | | | | | | | | | | | | Robinson 1956 |
| SK 46 | 14.5 | 17.2 | 15 | 16.6 | | | | | | | | | | | | | | | male | Schwartz and Tattersall 2005 | Wood 1991 |
| SK 48 | | | 13.5 | 15.9 | | | | | | | | | | | | | | | probable male | Lockwood et al. 2007 | Wood 1991 |
| SK 49 | | | 14.2 | 17 | | | | | | | | | | | | | | | male | Lockwood et al. 2007 | Wood 1991 |
| SK 55b | | | | | | | | | 9.8 | 10.6 | | | | | | | | | | | Wood 1991 |
| SK 72 | | | | | | | | | 10.3 | 12.2 | | | | | | | | | | | Wood 1991 |
| SK 74a | | | | | | | 9.9 | 10.4 | | | | | 11 | 12.1 | | | | | | | Wood 1991 |

| | | | | | | | | | | | | | | | | | | | | | |
|---|---|---|---|---|---|---|---|---|---|---|---|---|---|---|---|---|---|---|---|---|---|
| SK 80 | | | 12.3 | 15.3 | | | | | | | | | | | | | | | | | Wood 1991 |
| SK 81 | | | | | | | | | | | | | | | 10.8 | 13.5 | | | male | Grine et al. 2012 | Robinson 1956 |
| SK 83 | 15.7 | 17.3 | 15.8 | 17.6 | | | | | | | | | | | | | | | male | Keyser 2000 | Wood 1991 |
| SK 88 | | | | | | | | | | | | | | | 12.1 | 14.7 | | | | | Wood 1991 |
| SK 96 | | | | | | | | | 10 | 9.5 | | | | | | | | | probable female | Copeland et al. 2011 | Wood 1991 |
| SK 100 | | | | | | | 10.1 | 12.8 | | | | | | | | | | | male | Grine et al. 2012 | Wood 1991 |
| SK 105 | | | | | 15.6 | 15.7 | | | | | | | | | | | | | | | Robinson 1956 |
| SK 826b | | | | | | | | | | | | | | | 12.3 | 13.5 | | | male | Lockwood et al. 2007 | Wood 1991 |
| SK 827 | | | | | | | | | | | | | | | 10.9 | 11.6 | | | | | Wood 1991 |
| **SK 830** | | | | | | | | | | | | | | | 10.6 | 12.2 | | | | | Robinson 1956 |
| SK 831 | | | 17.2 | 17.8 | | | 9.8 | 11.5 | | | | | | | | | | | | | Wood 1991 |
| **SK 835** | | | 14.1 | 16.5 | | | | | | | | | | | | | | | female | Dean et al. 2020/Copeland et al. 2011 | Wood 1991 |
| SK 836 | | | 16.5 | 18.2 | | | | | | | | | | | | | | | | | Wood 1991 |
| SK 837 | | | | | 15.2 | 17 | | | | | | | | | | | | | | | Robinson 1956 |
| **SK 850** | | | | | | | 9.63 | | | | | | | | | | | | | | This study |
| SK 857 | | | | | | | 10.1 | 12.3 | | | | | | | | | | | | | Wood 1991 |
| SK 858 | | | | | | | | | | | 10 | 13.7 | | | | | 11 | | male | Lockwood et al. 2007 | Frayer 1973 |
| SK 870 | | | 14.2 | 16.8 | | | | | | | | | | | | | | | | | Wood 1991 |
| SK 876 | | | | | | | | | | | 10 | 11.7 | | | | | 10.5 | 12.5 | male | Sillen et al. 1998 | Frayer 1973 |
| SK 1587 | | | | | | | | | | | | | | | 10.8 | 11.7 | | | | | Wood 1991 |
| SK 1588 | | | | | | | | | | | | | 10.7 | 11.8 | | | | | female | Thackeray et al. 2005 | Wood 1991 |
| SK 1593 | | | | | | | 10.4 | 11.6 | | | | | | | | | | | | | Wood 1991 |
| SK 3975 | | | 14.7 | 16.7 | | | | | | | | | | | | | | | | | Wood 1991 |
| SK 3977 | 15.7 | 17.8 | | | | | | | | | | | | | | | | | | | Wood 1991 |

| | | | | | | | | | | | | | | | | | | | | | |
|---|---|---|---|---|---|---|---|---|---|---|---|---|---|---|---|---|---|---|---|---|---|
| SK 14132 | | | | | | | | | | | | | | | | | | | | | Unmeasurable |
| SKW 5 | | | | | | | 9.3 | 11.2 | | | | | 11.1 | 12.5 | 11.4 | 12.7 | | | male | Lockwood et al. 2007 | Grine 1993 |
| SKW 11 | 14.8 | 17 | 14.2 | 16.7 | | | | | | | | | | | | | | | male | Lockwood et al. 2007 | Grine 1993 |
| SKW 29 | 12.7 | | 12.7 | 16.2 | | | | | | | | | | | | | | | male | Lockwood et al. 2007 | Grine 1993 |
| SKX 311 | | | | | | | | | 9 | 11.9 | | | | | | | | | | | Grine 1989 |
| SKX 4446 | | | | | | | | | | | | | 11.9 | 12.5 | | | | | male | Lockwood et al. 2007 | Grine 1989 |
| SKX 21841 | 15.5 | 16.6 | | | | | | | | | | | | | | | | | | | Grine 1989 |
| SKX 32162 | | | | | | | | | | | | | 11.3 | | | | | | | | Grine 1989 |
| TM 1517 | 14.6 | 15.8 | | | | | | | 10.5 | 12.3 | | | | | 11.5 | 13.3 | | | male | Lockwood et al. 2007 | Wood 1991 |
| TM 1600 | | | | | | | | | 10.4 | 12.4 | | | | | | | | | | | Wood 1991 |
| TM 1601 | | | | | | | | | 9.6 | 11.2 | | | | | 10.9 | 12.1 | | | | | Wood 1991 |
| TM 1603 | | | 14.5 | 16.1 | | | | | | | | | | | | | | | | | Wood 1991 |
| DNH 3 | | | 13.6 | 16.3 | | | | | | | | | | | | | | | | | Moggi-Checci et al. 2010 |
| DNH 7 | 12.2 | 14.2 | 12.1 | 14.5 | | | 8.9 | 12.4 | 9.2 | 12.3 | | | 10.1 | 11.9 | 10.3 | 12.6 | | | female | Keyser 2000 | Keyser 2000 |
| DNH 8 | | | | | | | | | 10.2 | 12.6 | | | 11.4 | 13.4 | 11.3 | 13.6 | | | male | Keyser 2000 | Moggi-Checci et al. 2010 |
| DNH 15 | 14.2 | 16.1 | 14.8 | 16.4 | | | | | | | | | | | | | | | | | Moggi-Checci et al. 2010 |
| DNH 19 | | | | | | | | | 11 | | | | | | 12.3 | 13.2 | | | | | Moggi-Checci et al. 2010 |
| DNH 22 | 12.8 | 15.4 | | | | | | | | | | | | | | | | | | | Moggi-Checci et al. 2010 |
| DNH 26 | | | | | | | | | | | | | 10.9 | | | | | | | | Moggi-Checci et al. 2010 |
| DNH 27 | | | | | | | | | | | | | | | 11.2 | 12.9 | | | | | Moggi-Checci et al. 2010 |
| DNH 40 | | | 14.3 | 15.1 | | | | | | | | | | | | | | | | | Moggi-Checci et al. 2010 |
| DNH 51 | | | | | | | 10.9 | 11.3 | | | | | 11 | 12.5 | | | | | | | Moggi-Checci et al. 2010 |
| DNH 54 | | | 14 | 14.2 | | | | | | | | | | | | | | | | | Moggi-Checci et al. 2010 |
| DNH 58 | | | | | | | | | 11.3 | | | | | | | | | | | | Moggi-Checci et al. 2010 |
| DNH 68 | | | | | | | 10 | 12.9 | | | | | 9.9 | | | | | | male | Lockwood et al. 2007 | Moggi-Checci et al. 2010 |

| | | | | | | | | | | | | | | | | | | | | | |
|---|---|---|---|---|---|---|---|---|---|---|---|---|---|---|---|---|---|---|---|---|---|
| DNH 155 | 14.4 | 15.3 | 12.8 | 15 | | | | | | | | | | | | | | | male | Martin et al. 2021 | Martin et al. 2021 |
