## Supplementary Document 4 for "Enamel proteins reveal biological sex and genetic variability within southern African *Paranthropus*"

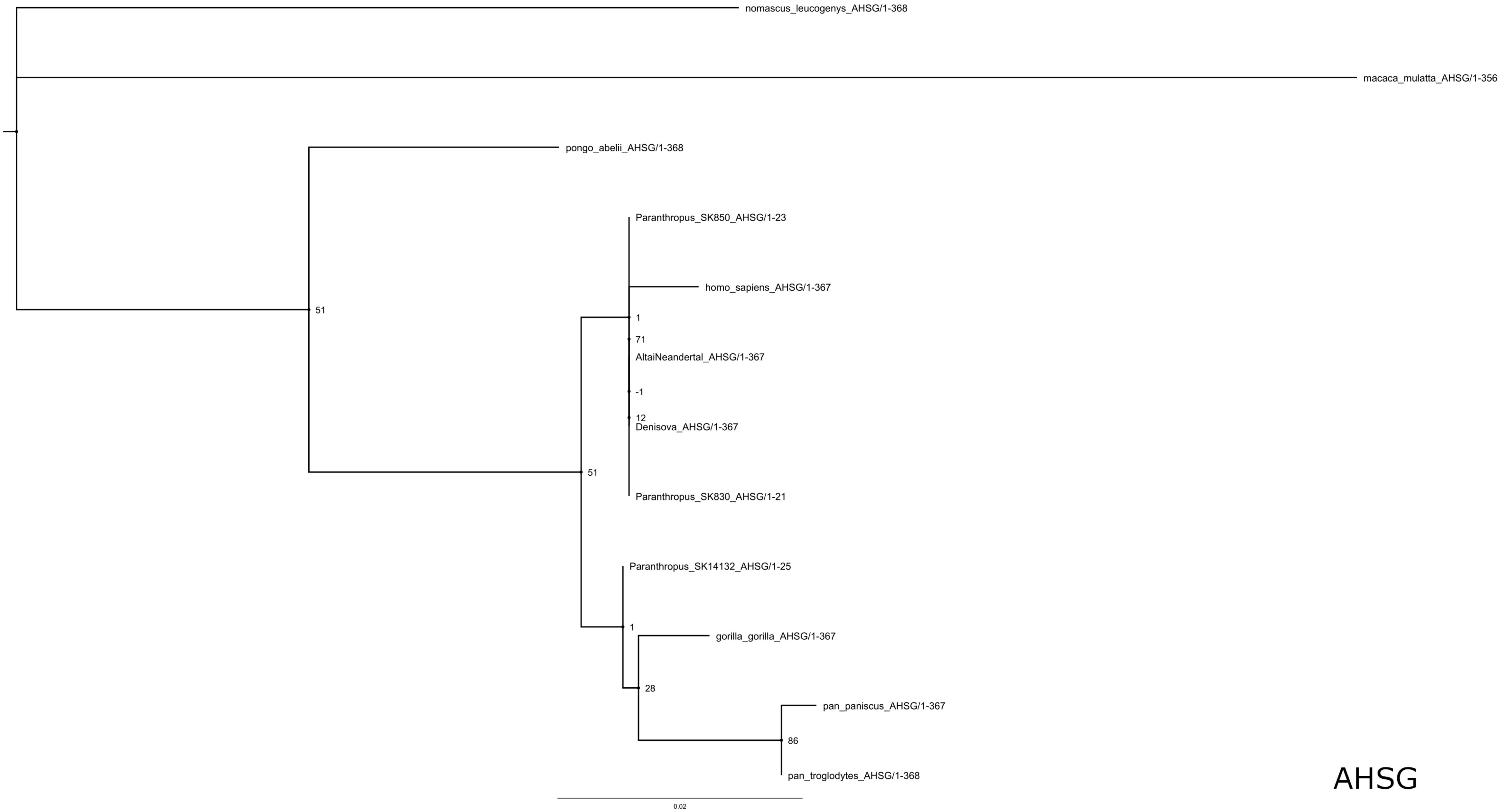

nomascus_leucogenys_AHSG/1-368
macaca_mulatta_AHSG/1-356
pongo_abelii_AHSG/1-368
51
Paranthropus_SK850_AHSG/1-23
homo_sapiens_AHSG/1-367
1
71
AltaiNeandertal_AHSG/1-367
-1
12
Denisova_AHSG/1-367
51
Paranthropus_SK830_AHSG/1-21
Paranthropus_SK14132_AHSG/1-25
1
gorilla_gorilla_AHSG/1-367
28
pan_paniscus_AHSG/1-367
86
pan_troglodytes_AHSG/1-368
0.02
AHSG

### ALB

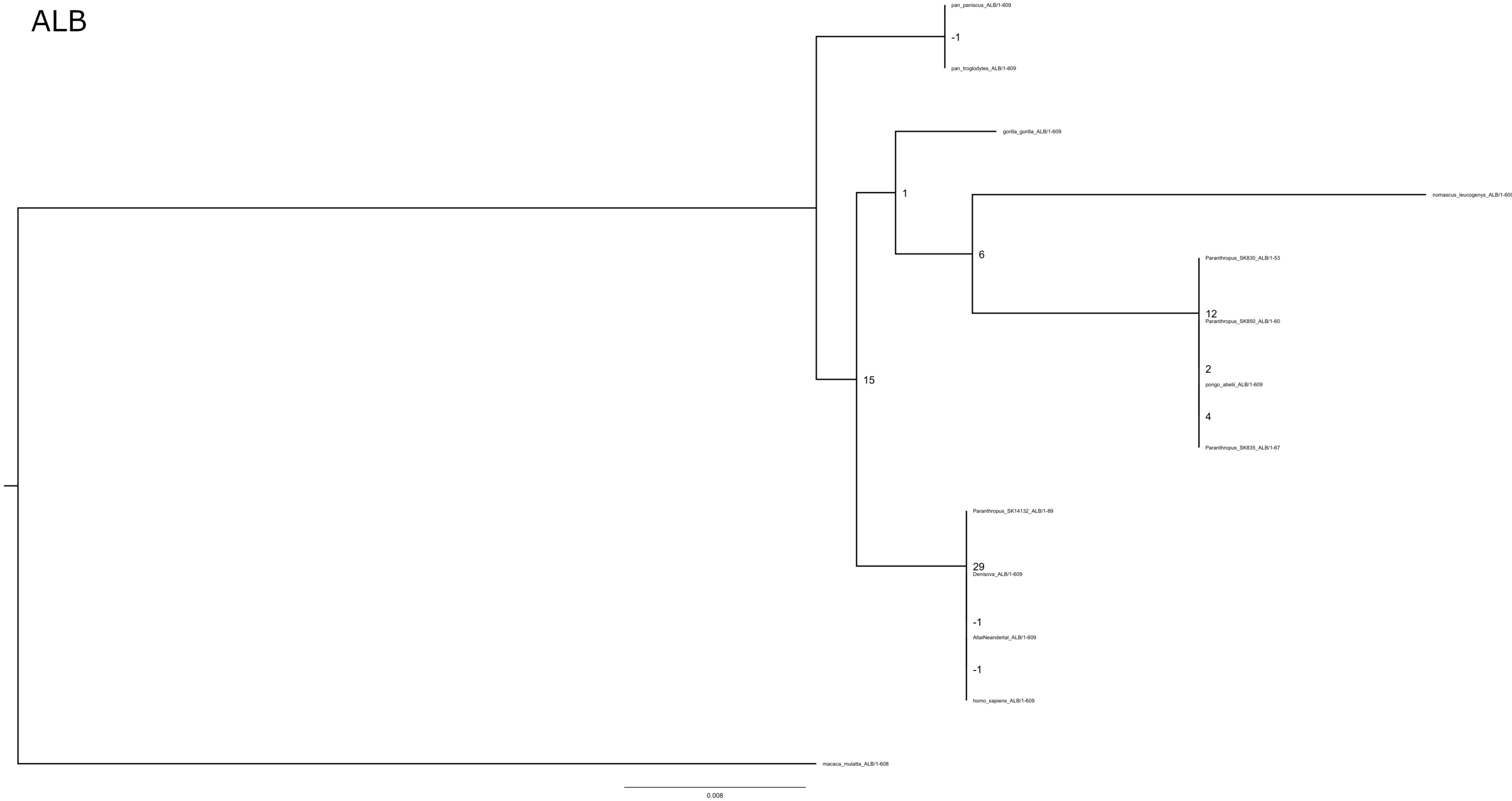
pan_paniscus_ALB/1-609
-1
pan_troglodytes_ALB/1-609
gorilla_gorilla_ALB/1-609
1
nomascus_leucogenys_ALB/1-609
6
Paranthropus_SK830_ALB/1-53
12
Paranthropus_SK850_ALB/1-60
2
pongo_abelii_ALB/1-609
4
Paranthropus_SK835_ALB/1-67
15
Paranthropus_SK14132_ALB/1-89
29
Denisova_ALB/1-609
-1
AltaiNeandertal_ALB/1-609
-1
homo_sapiens_ALB/1-609
macaca_mulatta_ALB/1-608
0.008

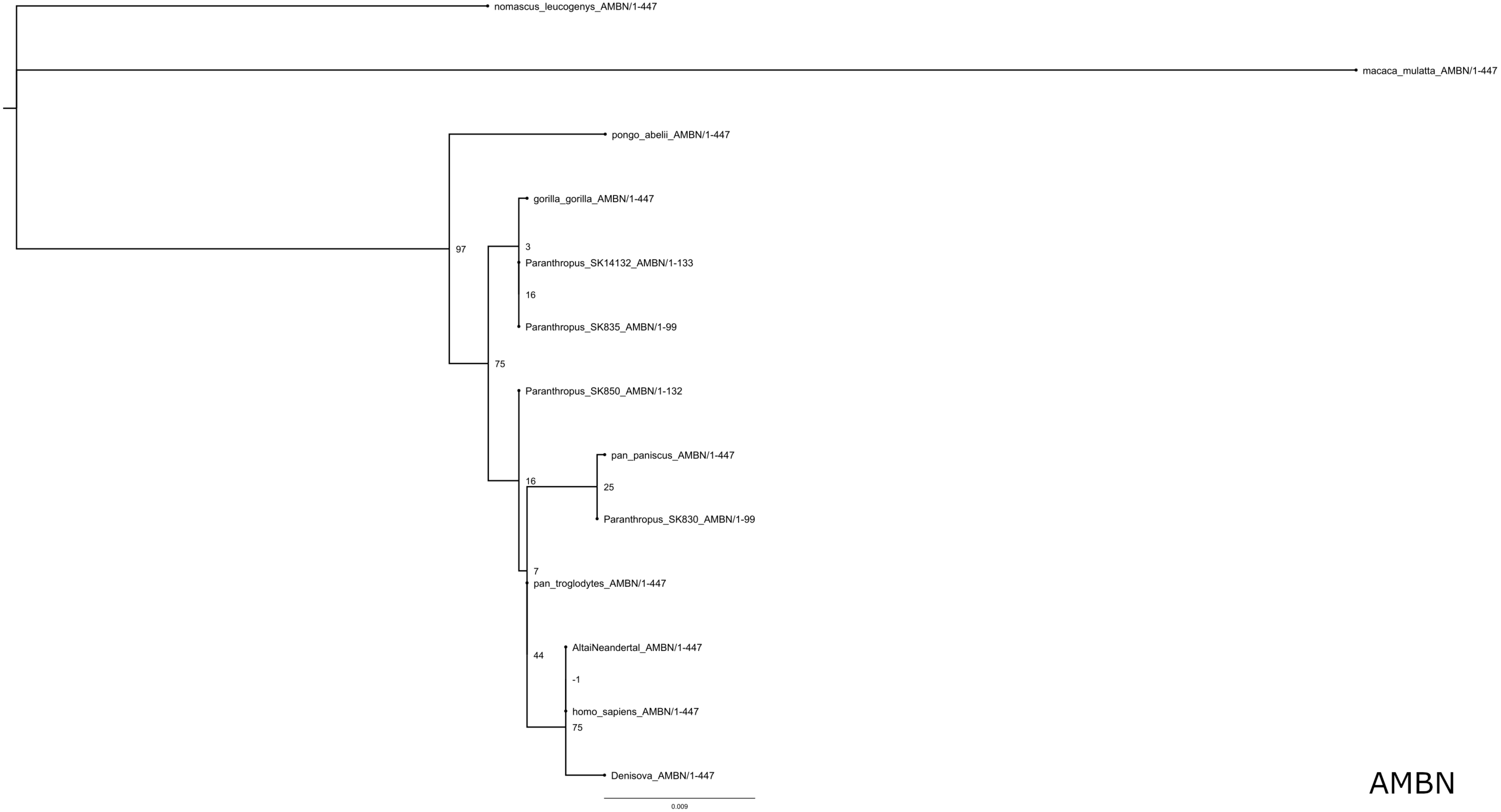

nomascus_leucogenys_AMBN/1-447
macaca_mulatta_AMBN/1-447
pongo_abelii_AMBN/1-447
97
gorilla_gorilla_AMBN/1-447
3
Paranthropus_SK14132_AMBN/1-133
16
Paranthropus_SK835_AMBN/1-99
75
Paranthropus_SK850_AMBN/1-132
pan_paniscus_AMBN/1-447
16
25
Paranthropus_SK830_AMBN/1-99
7
pan_troglodytes_AMBN/1-447
44
AltaiNeandertal_AMBN/1-447
-1
homo_sapiens_AMBN/1-447
75
Denisova_AMBN/1-447
0.009


AMBN

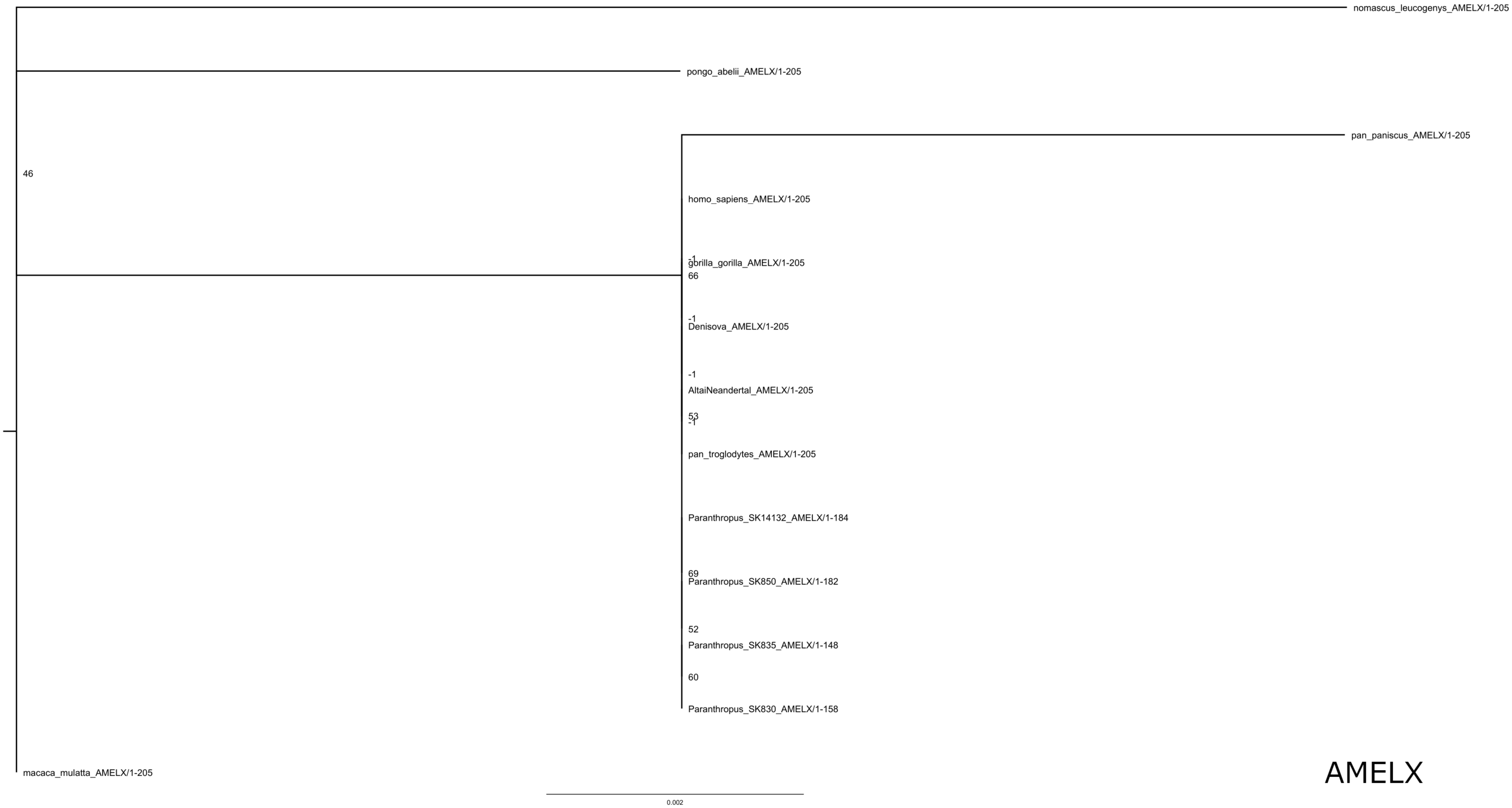
nomascus_leucogenys_AMELX/1-205
pongo_abelii_AMELX/1-205
pan_paniscus_AMELX/1-205
46
homo_sapiens_AMELX/1-205
-1
gorilla_gorilla_AMELX/1-205
66
-1
Denisova_AMELX/1-205
-1
AltaiNeandertal_AMELX/1-205
53
-1
pan_troglodytes_AMELX/1-205
Paranthropus_SK14132_AMELX/1-184
69
Paranthropus_SK850_AMELX/1-182
52
Paranthropus_SK835_AMELX/1-148
60
Paranthropus_SK830_AMELX/1-158
macaca_mulatta_AMELX/1-205
0.002
AMELX

### AMELY

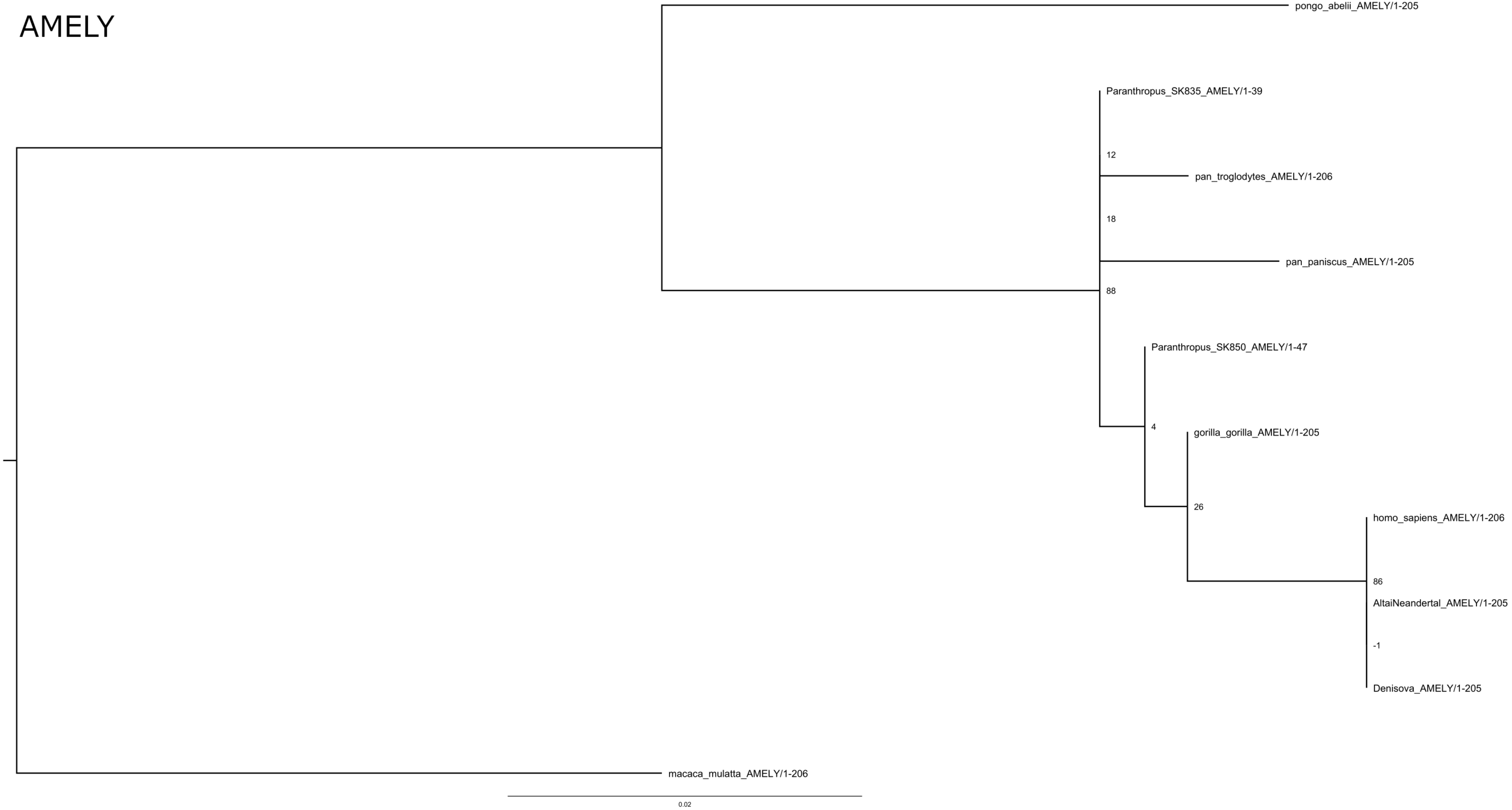

pongo_abelii_AMELY/1-205
Paranthropus_SK835_AMELY/1-39
12
pan_troglodytes_AMELY/1-206
18
pan_paniscus_AMELY/1-205
88
Paranthropus_SK850_AMELY/1-47
4
gorilla_gorilla_AMELY/1-205
26
homo_sapiens_AMELY/1-206
86
AltaiNeandertal_AMELY/1-205
-1
Denisova_AMELY/1-205
macaca_mulatta_AMELY/1-206
0.02

### AMTN

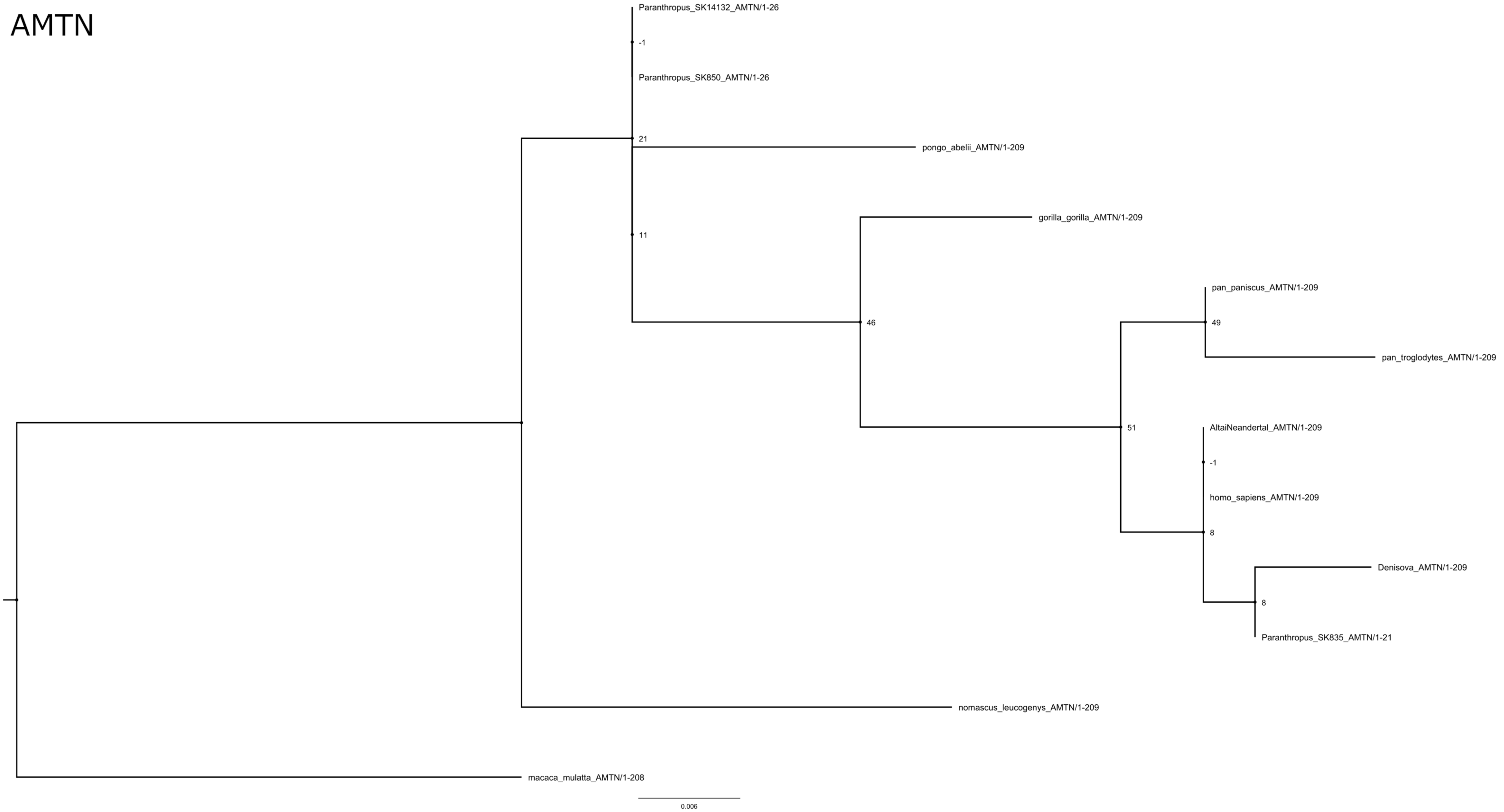
Paranthropus_SK14132_AMTN/1-26
-1
Paranthropus_SK850_AMTN/1-26
21
pongo_abelii_AMTN/1-209
11
gorilla_gorilla_AMTN/1-209
pan_paniscus_AMTN/1-209
46
49
pan_troglodytes_AMTN/1-209
51
AltaiNeandertal_AMTN/1-209
-1
homo_sapiens_AMTN/1-209
8
Denisova_AMTN/1-209
8
Paranthropus_SK835_AMTN/1-21
nomascus_leucogenys_AMTN/1-209
macaca_mulatta_AMTN/1-208
0.006

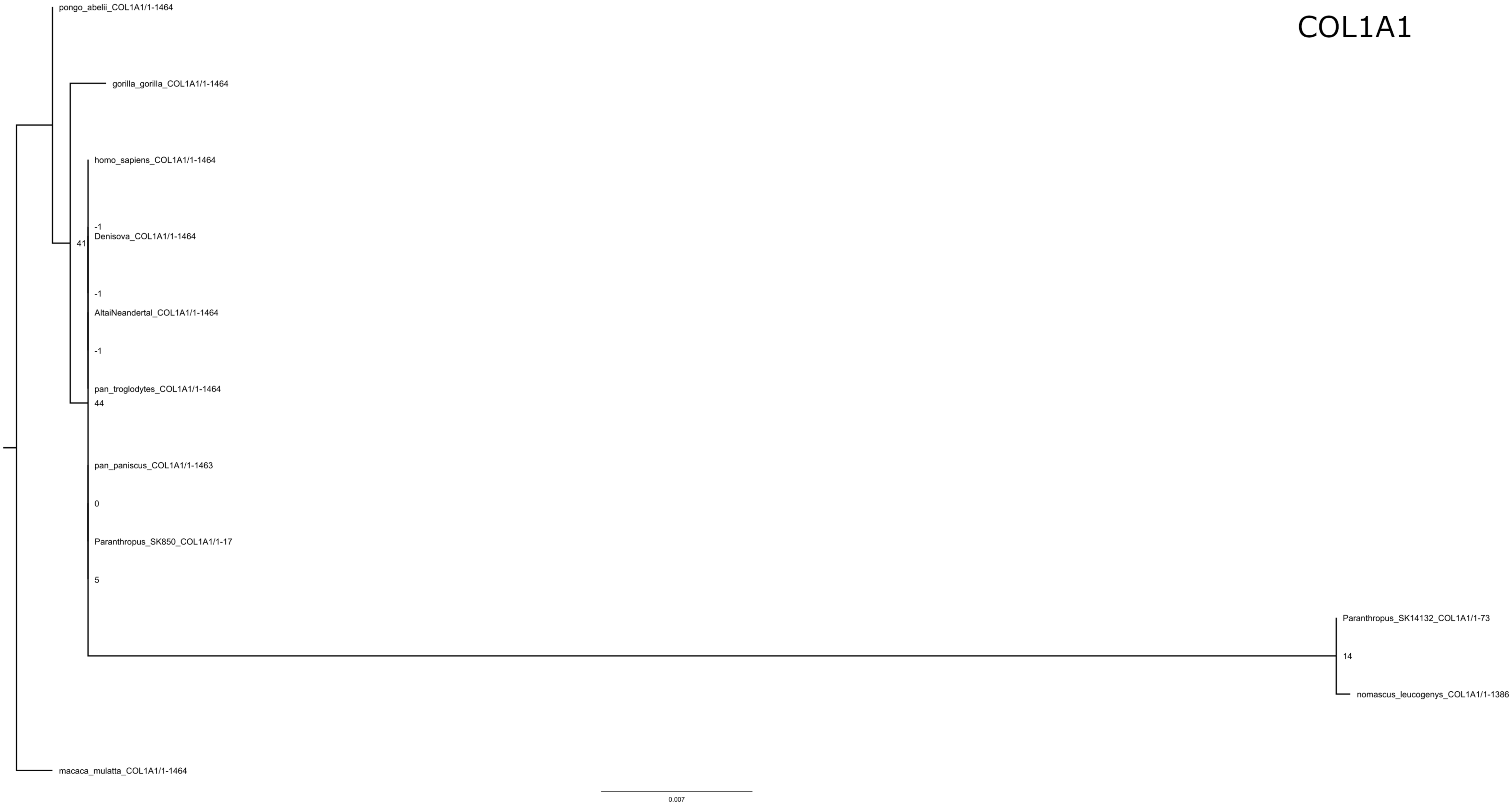
COL1A1
pongo_abelii_COL1A1/1-1464
gorilla_gorilla_COL1A1/1-1464
homo_sapiens_COL1A1/1-1464
-1
Denisova_COL1A1/1-1464
41
-1
AltaiNeandertal_COL1A1/1-1464
-1
pan_troglodytes_COL1A1/1-1464
44
pan_paniscus_COL1A1/1-1463
0
Paranthropus_SK850_COL1A1/1-17
5
Paranthropus_SK14132_COL1A1/1-73
14
nomascus_leucogenys_COL1A1/1-1386
macaca_mulatta_COL1A1/1-1464
0.007

### COL17A1

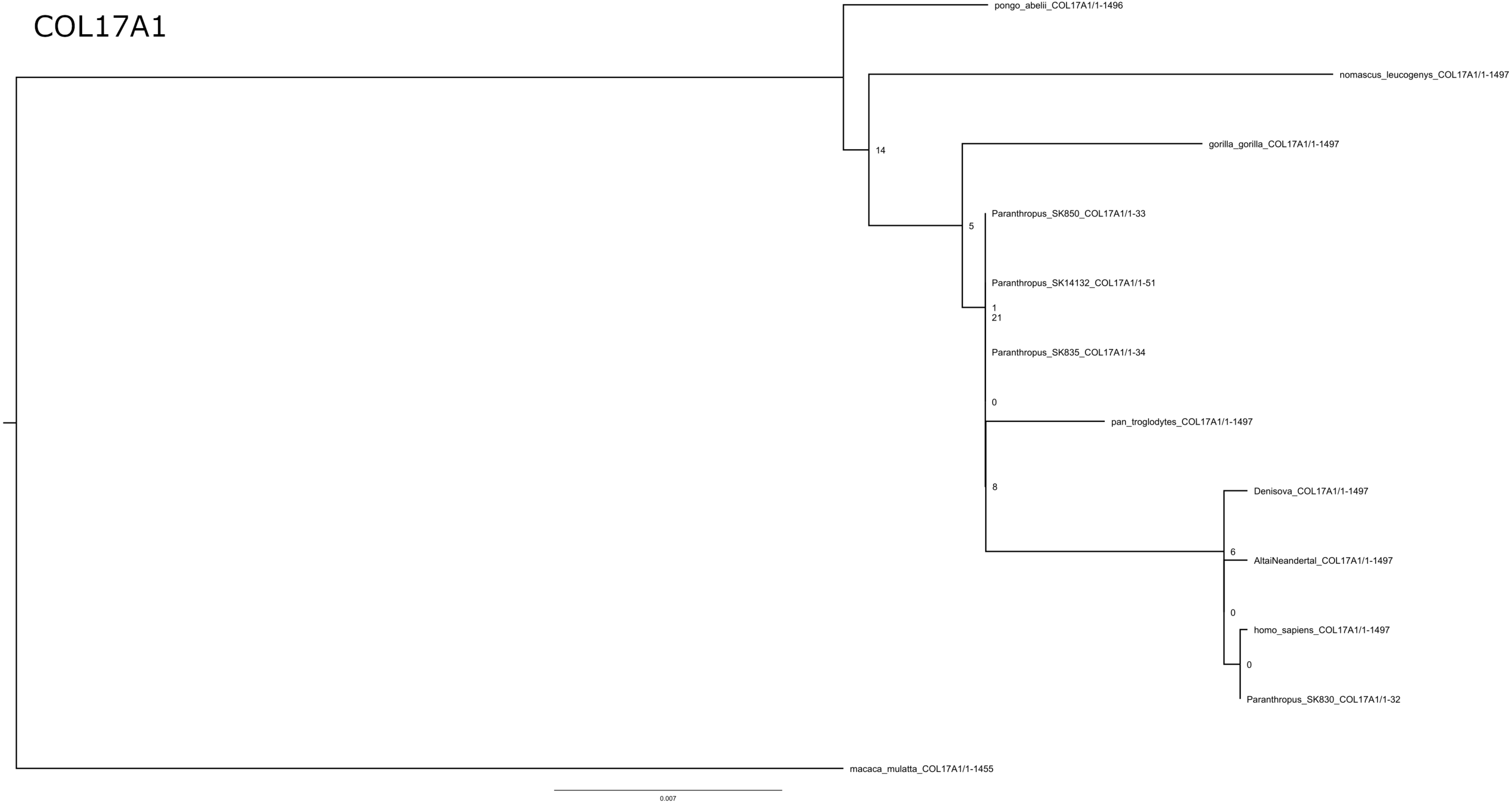

pongo_abelii_COL17A1/1-1496
nomascus_leucogenys_COL17A1/1-1497
14
gorilla_gorilla_COL17A1/1-1497
5
Paranthropus_SK850_COL17A1/1-33
Paranthropus_SK14132_COL17A1/1-51
1
21
Paranthropus_SK835_COL17A1/1-34
0
pan_troglodytes_COL17A1/1-1497
8
Denisova_COL17A1/1-1497
6
AltaiNeandertal_COL17A1/1-1497
0
homo_sapiens_COL17A1/1-1497
0
Paranthropus_SK830_COL17A1/1-32
macaca_mulatta_COL17A1/1-1455
0.007

### ENAM

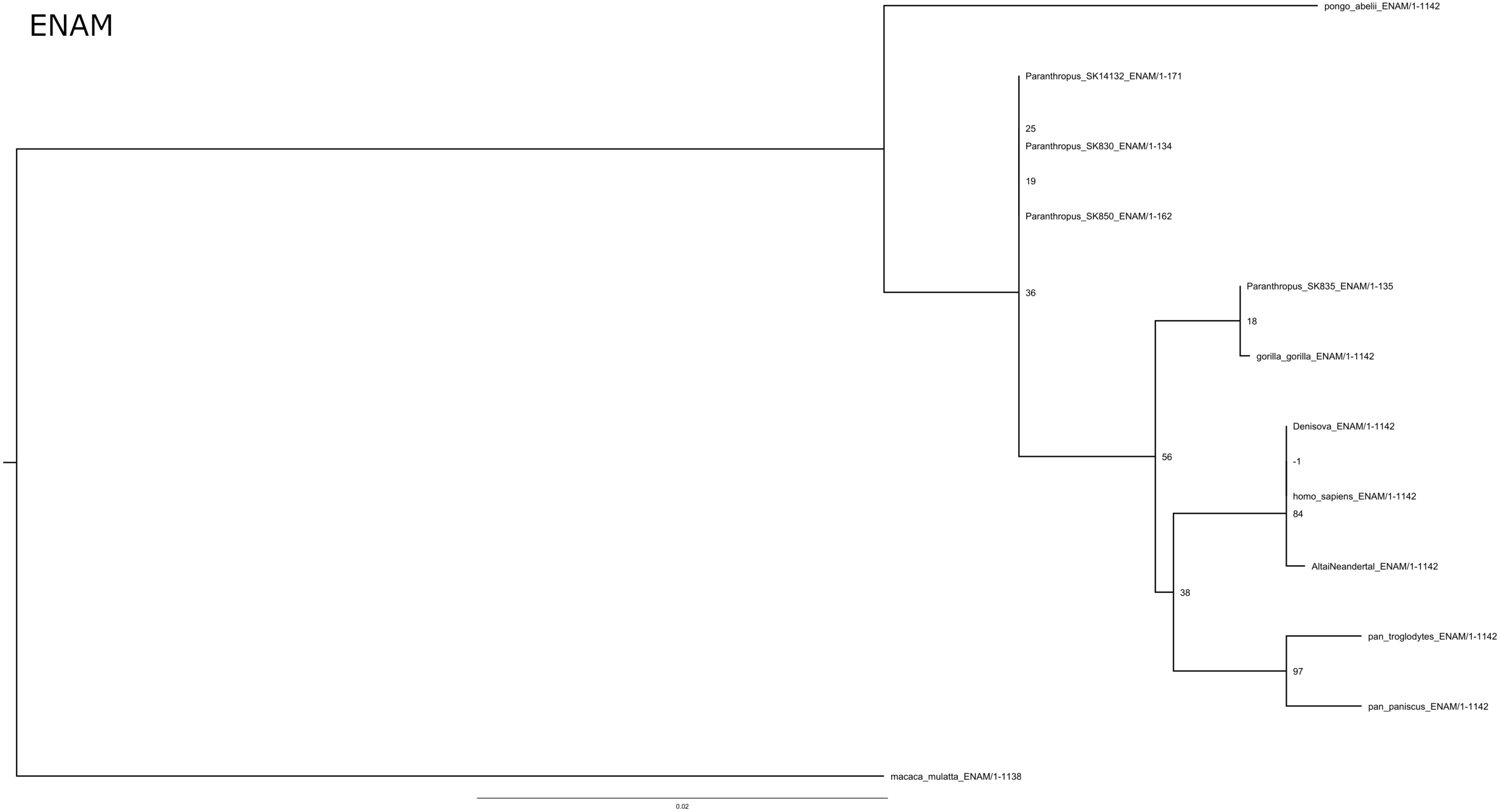
pongo_abelii_ENAM/1-1142
Paranthropus_SK14132_ENAM/1-171
25
Paranthropus_SK830_ENAM/1-134
19
Paranthropus_SK850_ENAM/1-162
36
Paranthropus_SK835_ENAM/1-135
18
gorilla_gorilla_ENAM/1-1142
Denisova_ENAM/1-1142
-1
56
homo_sapiens_ENAM/1-1142
84
AltaiNeandertal_ENAM/1-1142
38
pan_troglodytes_ENAM/1-1142
97
pan_paniscus_ENAM/1-1142
macaca_mulatta_ENAM/1-1138
0.02

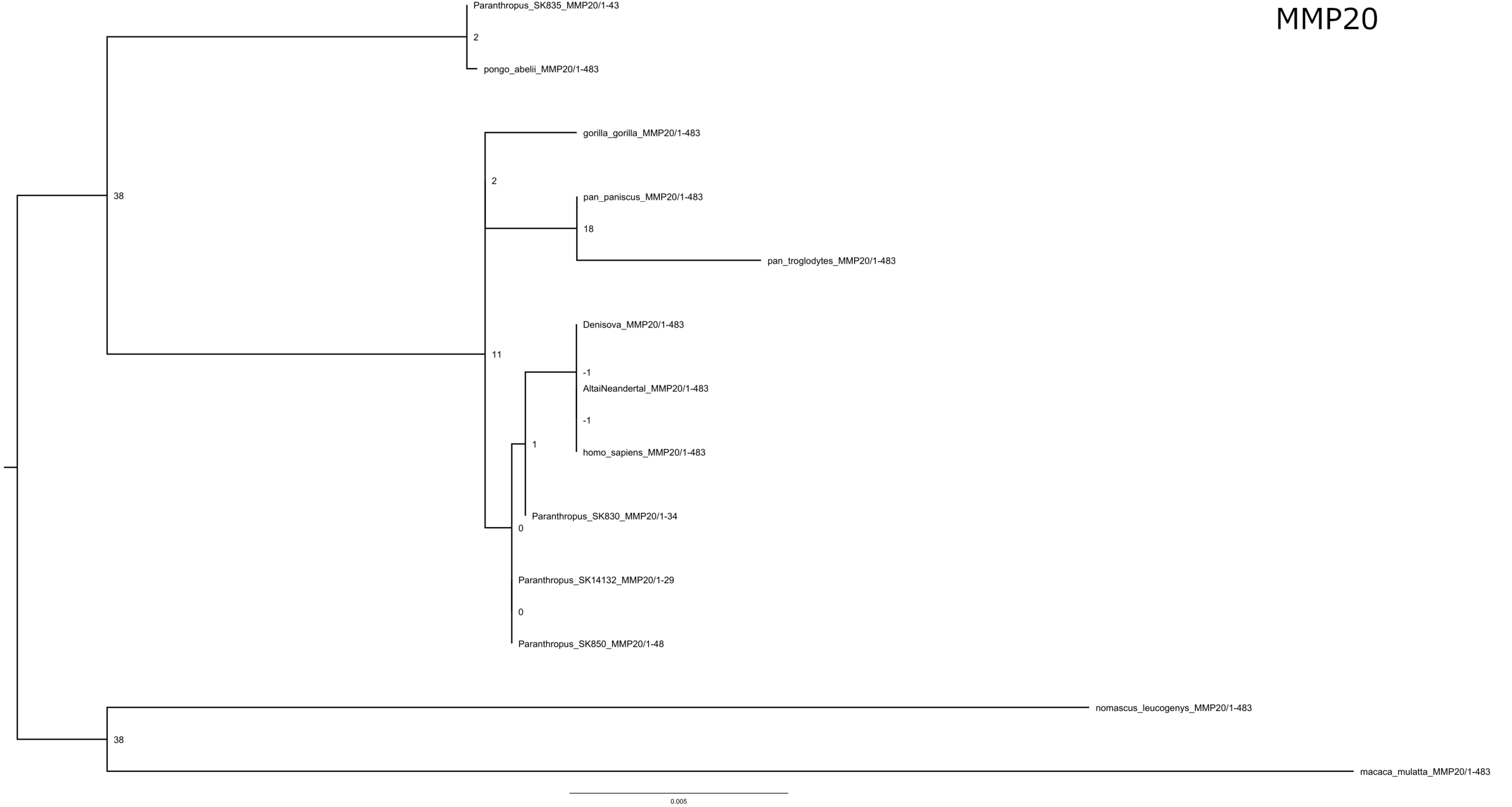
MMP20
Paranthropus_SK835_MMP20/1-43
2
pongo_abelii_MMP20/1-483
38
gorilla_gorilla_MMP20/1-483
2
pan_paniscus_MMP20/1-483
18
pan_troglodytes_MMP20/1-483
Denisova_MMP20/1-483
11
-1
AltaiNeandertal_MMP20/1-483
-1
1
homo_sapiens_MMP20/1-483
Paranthropus_SK830_MMP20/1-34
0
Paranthropus_SK14132_MMP20/1-29
0
Paranthropus_SK850_MMP20/1-48
nomascus_leucogenys_MMP20/1-483
38
macaca_mulatta_MMP20/1-483
0.005

ODAM

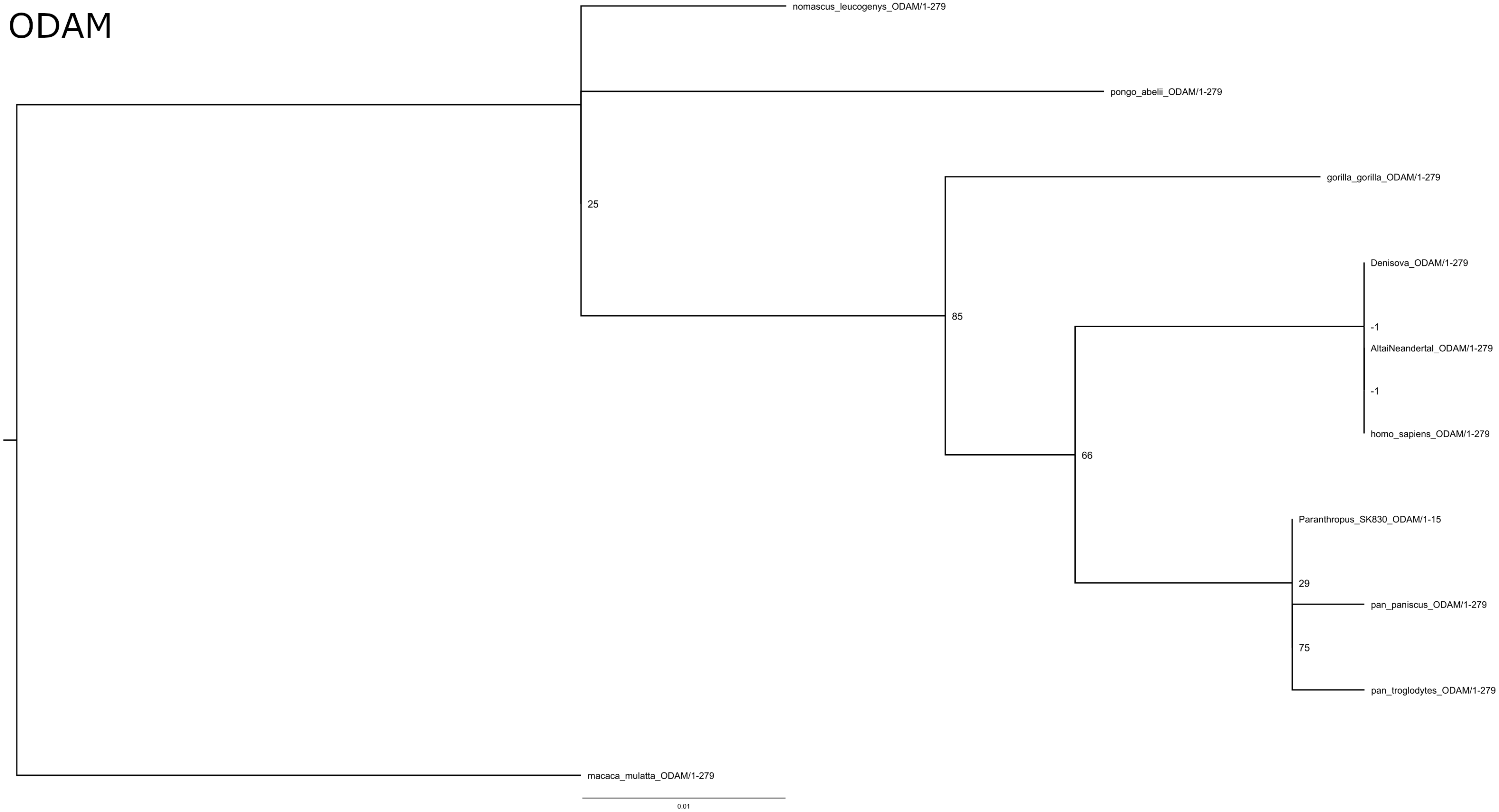
nomascus_leucogenys_ODAM/1-279
pongo_abelii_ODAM/1-279
25
gorilla_gorilla_ODAM/1-279
Denisova_ODAM/1-279
85
-1
AltaiNeandertal_ODAM/1-279
-1
homo_sapiens_ODAM/1-279
66
Paranthropus_SK830_ODAM/1-15
29
pan_paniscus_ODAM/1-279
75
pan_troglodytes_ODAM/1-279
macaca_mulatta_ODAM/1-279
0.01
